# Population Differentiation of *Rhodobacteraceae* Along Coral Compartments

**DOI:** 10.1101/2020.12.31.424895

**Authors:** Danli Luo, Xiaojun Wang, Xiaoyuan Feng, Mengdan Tian, Sishuo Wang, Sen-Lin Tang, Put Ang, Aixin Yan, Haiwei Luo

## Abstract

Coral mucus, tissue and skeleton harbor compositionally different microbiota, but how these coral compartments shape the microbial evolution remains unexplored. Here, we focused on the *Rhodobacteraceae*, which represents a significant but variable proportion (5-50%) of the coral microbiota. We sequenced 234 genomes constituting two divergent populations inhabiting a prevalent coral species *Platygyra acuta*. One population diverged into two clades colonizing the mucus and skeleton respectively. We reconstructed the ancestral gene changing events that potentially drove the split, and found that the affected genes matched well with the distinct physicochemical features of the mucus and skeleton. Specifically, the mucus clade acquired functions involved in the utilization of coral osmolytes abundant in the mucus (e.g., methylamines, DMSP, taurine and L-proline), whereas the skeleton clade uniquely harbored traits that may promote adaptation to the low-energy and diurnally anoxic skeleton (e.g., sulfur oxidation and swimming motility). These between-clade genetic differences were largely supported by physiological assays. Expanded analyses by including relatives isolated from various marine environments suggest that the mucus and skeleton clades may have diversified in non-coral habitats, but they also consolidated a key role of distinct coral compartments in diversifying many of the above-mentioned traits. The second population varied only at a few dozen nucleotide sites across the whole genomes, and the Slatkin-Maddison test supported that dispersal limitation between coral compartments is another key mechanism driving microbial population differentiation. Collectively, our results suggest that different coral compartments represent ecologically distinct and microgeographically separate habitats that drive the evolution of the coral microbiota.

## Introduction

As the home to more than 25% of marine life, coral reefs are an important habitat promoting biodiversity in the ocean [1]. The coral holobiont is a dynamic community of coral polyps, microalgae, fungi, prokaryotes and viruses [2]. Scleractinian corals harbor three major compartments microbial populations inhabit (Fig. S1), namely coral mucus, skeleton and tissue [2]. The coral mucus is a nutrient and energy-rich mixture of materials [3, 4], and its primary structural macromolecules are glycoproteins, polysaccharides and lipids [5, 6]. In addition, mucus contains rich organic osmolytes secreted by the coral host and its endosymbiont *Symbiodiniaceae* [7]. The osmolytes are compatible or counteracting solutes that can be accumulated in the cells in response to the fluctuation of osmotic stresses, thermal stress and hypoxia [8, 9], which includes methylamines [e.g., glycine betaine (GBT), choline, choline-O-sulfate (COS), sarcosine and trimethylamine N-oxide (TMAO)], methyl sulfonium [e.g., dimethylsulfoniopropionate (DMSP) and dimethyl sulfoxide (DMSO)], amino acid derivatives (e.g., taurine, L-proline and ectoine), sugars (e.g., trehalose and arabinose) and polyols (e.g., glycerol) [9]. Of these, GBT, taurine and DMSP represent the major osmolytes of corals [10]. The coral animals eliminate or accumulate the osmolytes quickly to equilibrate the fluctuated salinity caused by, for example, tides, air exposure, evaporation, and heavy rainfalls [11]. These processes together lead to the enrichment of wasted osmolytes in mucus [12], which serve as organic nutrients to support the resident bacteria [13, 14]. In contrast, the skeleton is a low-energy environment embedded in the coral tissue, where up to 99% of the photosynthetically active radiation is either scattered or entrapped by the *Symbiodiniaceae* located at coral tissue [15]. The porous structure of the skeleton is mainly constructed with inorganic aragonite crystals along with a small fraction (≤ 0.1% by weight) of the organic matrix [16, 17] which limits skeleton-associated microbes [18]. Although the light penetration is low, there are still endolithic algae that reside inside the skeleton, whose photosynthesis and respiration generate a diel fluctuation of the oxygen and pH levels in the skeleton [15]. Likewise, the *Symbiodiniaceae* in the coral cells causes similar diel changes of the tissue [19].

The fact that different coral compartments are featured with distinct physicochemical conditions indicates that coral is an ideal system for studying microbial niche adaptation. Previous studies suggest that the community structure of the bacterial associates are largely shaped by the coral anatomy [20, 21]. Among the three compartments, for example, the mucus harbors a higher proportion of compartment-specific bacteria than the tissue [22] and the skeleton accommodates a microbial community of greater diversity compared to the tissue and mucus [15, 23]. While these studies illustrated potentially important differences in terms of the microbial identities and functions across the coral compartments, they provided limited taxonomic resolution in functional differentiation along coral compartments. Since functional differences can be ascribed to both environmental heterogeneity and phylogenetic diversity, a robust association of metabolic functions with ecological niches may be established when the phylogenetic distance of lineages under comparison is minimized. Leveraging the single nucleotide polymorphism (SNP) distribution at the core genome and gene frequency at the accessory genome, the culture-based population genomics is able to uncover the mechanisms driving population differentiation and niche adaptation [24–27]. To our knowledge, this approach has not been used to investigate coral-associated microbial populations.

Among the numerous bacterial groups found on corals, *Ruegeria* is among the top three genera that are most widely associated with coral species [28]. *Ruegeria* is a genus affiliated with alphaproteobacterial family *Rhodobacteraceae*. Members of *Ruegeria* likely establish mutualistic interactions with the coral hosts. For instance, they are constantly associated with both broadcast-spawning and brooding corals in their early developmental stages [29], and they are one of the key bacterial taxa that can be vertically transferred from parents to larvae in the brooding coral *Pocillopora damicornis* [30]. Some *Ruegeria* strains also help the larval settlement in octocoral species [31]. In adult corals, *Ruegeria* strains can inhibit the growth of pathogenic *Vibrio* species [32]. Nevertheless, some *Ruegeria* members may be potential opportunistic pathogens, which show increased abundance when coral hosts are under stress [33–35].

*Platygyra acuta*, the scleractinian coral sampled in the present study, is known for its high tolerance to temperature and salinity stresses, as well as its resistance to bleaching in winter [36]. It is not only the dominant species in Hong Kong coastal waters, but also a globally important germplasm resource to understand the survival of marginal reefs [37, 38]. We harvested two *Rhodobacteraceae* populations from *P. acuta*: one was a member of *Ruegeria* (hereafter, the *Ruegeria* population) and the other cannot be assigned to any described genus in the *Rhodobacteraceae* (hereafter, the *Rhodobacteraceae* population). Both populations are members of the *Roseobacter* group, a phylogenetic group encompassing the most taxonomic diversity found in the *Rhodobacteraceae* [39]. The two populations consisted of 20 and 214 isolates, respectively. Members of the *Ruegeria* population harbored a great amount of genetic variation, and their phylogenetic clustering and metabolic differences matched well with the coral compartment differentiation. While members of the *Rhodobacteraceae* population varied at only a few dozen SNP sites across the whole genomes, they nevertheless displayed a microbiogeographic pattern across coral compartments, which is a signature of population differentiation due to dispersal limitation.

## Materials and Methods

Samples of *P. acuta* and ambient seawater were collected by SCUBA diving in Hong Kong waters from April 2017 to February 2018. Bacteria were isolated from the three coral compartments including mucus, skeleton and tissue, and also from the ambient seawater. We identified two populations representing divergent lineages of *Rhodobacteraceae* suitable for subsequent population genomic analyses according to two criteria: i) there were a large number of isolates within each population and ii) multiple coral compartments were sampled in each population. One population is a member of the ubiquitous coral-associated genus *Ruegeria* (20 isolates), and the other cannot be assigned to any described genus and was thus termed the *Rhodobacteraceae* population (214 isolates). Genomes of all these 234 isolates were sequenced with BGISEQ-500 PE100, assembled and annotated. The strain HKCCD4315 in the *Ruegeria* population was additionally sequenced using the PacBio Sequel platform to obtain a closed genome.

The *Ruegeria* population was initially analyzed to understand its phylogenetic and population genetic structure. The orthologous gene families were identified with OrthoFinder v2.2.1 [40] and the maximum likelihood phylogenomic tree was constructed based on the concatenated single-copy gene alignments at the amino acid level using IQ-TREE v1.6.5 [41]. Two clades (namely, clade-M and clade-S) matched well with the coral mucus and skeleton compartments, respectively, along with an additional clade locating at the outgroup position. The pairwise 16S rRNA gene identity and the whole-genome average nucleotide identity (ANI) between- and within-clades were compared to help determine whether speciation has already occurred between these clades [42]. Phylogenetic separation does not necessarily lead to allelic isolation between the phylogenetic clusters [43, 44]. We therefore investigated the population structure of clade-M and clade-S with the coancestry analysis implemented in fineSTRUCTURE v2.0.7 [45], and characterized the potential gene flow barrier by comparing the between-clade to within-clade relative frequency of recombination to point mutation (*ρ/θ*) [46] and relative effect of recombination to mutation (r/m) [46], and by determining the fixation index (F_st_) across the core genome shared by clade-M and clade-S [47]. We further tested whether the genetic separation of the two clades occurred at the accessory genome by measuring the similarity of genome content among strains with the Jaccard index and subsequently clustering it with the complete-linkage method [27].

The population differentiation between clade-M and clade-S of the *Ruegeria* population was further investigated at the functional level. We sought to identify signature genes of potential importance in niche adaptation from both core and accessory genome. In the core genome, novel allelic replacements via homologous recombination with external species at certain core loci represent a potentially important and prevalent adaptive mechanism in marine bacterial populations [44, 48]. This type of core genes was also identified here. Briefly, the substitution rate at synonymous (silent and thus largely neutral) sites (*d_S_*) across all single-copy core genes and across all pairwise comparisons between isolates from clade-M and clade-S was clustered using the K-means method, which resulted in a cluster composed of genes each showing unusually large *d_S_* values, the genetic signature of an evolution history subjected to novel allelic replacement [48]. For the identified core genes, gene trees were built and compared to the genome tree to help determine which clade was subjected to novel allelic replacement. In terms of the accessory genome, we focused on the genes largely specific to clade-M (or clade-S) and tried to resolve two competing hypotheses: whether they resulted from acquisition events at the last common ancestor (LCA) of clade-M (or clade-S) or loss events at the LCA of clade-S (or clade-M). Specifically, the gene gain and loss history of clade-specific genes were inferred based on the gene copy number distribution among extant genomes of the *Ruegeria* population using BadiRate v1.35 [49]. and checked manually based on the distribution pattern of pseudogenes in the context of gene clusters [50]. For genes whose functions were potentially relevant to niche adaptation and which differentiated the clade-M and clade-S at the DNA level, their potential functional outcomes were tested using physiological assays, though we did not have direct evidence connecting the genetic and phenotypic differentiation.

To test whether the clade-M and clade-S may have evolved from a single ancestor initially colonized on the coral or these two clades may have already split in marine habitats unrelated to corals and subsequently independently colonized coral compartments, we sampled and sequenced 22 closely related *Ruegeria* isolates mostly from other marine habitats in Hong Kong, constructed an expanded phylogenomic tree, and inferred the evolutionary history of the ecological relevant genes using the above-mentioned methods. For the flagellar gene cluster, its evolutionary history cannot be simply resolved based on the gene copy number distribution pattern. In this case, pseudogenes may provide unequivocal evidence for the presence of this gene cluster in the ancestral populations, and they were identified following a previous study [50]. We additionally estimated the potential time of the split between clade-M and clade-S based on synonymous substitution rate, spontaneous mutation rate and potential generation time in the wild, and the goal of this analysis to show its antiquity compared to the coral ages.

For the *Rhodobacteraceae* population, the genetic uniformity enabled the Slatkin-Maddison test [51] to measure the degree of compartmentalization on population subdivision. We also estimated the time of its origin and compared it to the potential age of the coral animal. All the methodological details were described in Supplementary Text 1.

## Results and Discussion

### Features of a Ruegeria population isolated from different coral compartments

The phylogenomic tree showed that the 20 closely related *Ruegeria* strains isolated from *P. acuta* formed three clades (Fig. 1A). Among these, the six clade-S members were all isolated from the skeleton, and the six clade-M members were all collected from mucus except for one from ambient seawater (Fig. 1A). The outgroup clade consisted of seven skeleton-associated strains and one tissue-associated strain. As these strains were isolated from the same coral species, interpretation of any strain-level genomic differences ruled out the host species effect. Therefore, our finding of the phylogenetic clustering by their source compartments indicates that adaptation to the distinct coral compartments likely drives the evolutionary diversification of this *Ruegeria* population [23].

**Figure 1.**
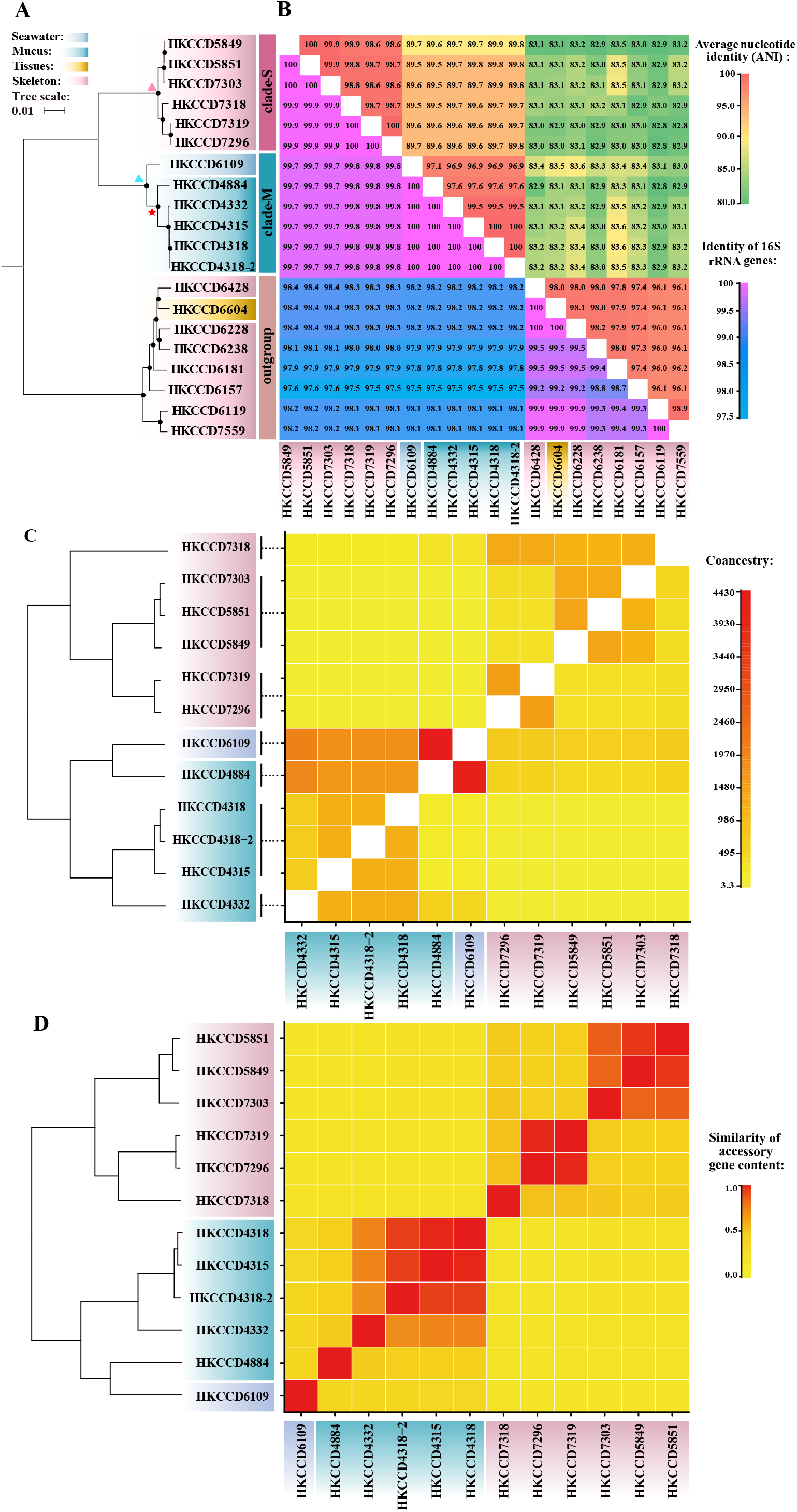
The phylogeny and population differentiation of the *Ruegeria* population. **(A)** The rooted maximum-likelihood phylogenomic tree of the 20 strains isolated from the coral species *Platygyra acuta* (Accession numbers were showed in Table S6). This tree is rooted with the mid-point rooting method. Solid circles at the nodes indicate that the support value (IQ-TREE v1.6.5 ultrafast bootstraps) of the branch is 100%. The LCA of clade-M, the LCA of the five mucus strains within the clade-M, and the LCA of clade-S are shown as blue triangle, the red star, and the pink triangle, respectively. Strains isolated from different coral compartments are highlighted with different colors. The clade-M and clade-S are also labeled. **(B)** The heatmap of the whole-genome average nucleotide identity (ANI) and heatmap of the pairwise identity of 16S rRNA genes of the 20 strains. **(C)** fineSTRUCTURE coancestry matrix of the 12 isolates from clade-M and clade-S, with warmer colors representing more ancestry shared between strains under comparison. Strains assigned to the same fineSTRUCTURE population are connected with a vertical bar on the left of the matrix, and the dendrogram shows the clustering of the fineSTRUCTURE populations. **(D)** Heatmap of accessory gene content similarity measured by Jaccard distance, with warmer colors representing the higher similarity between strains. The left dendrogram was generated based on the complete-linkage clustering method.

We further investigated the ANI and 16S rRNA gene identity for these 20 strains, which are standard measures typically used for bacteria species delineation [42, 52]. The genetic clusters based on ANI or 16S rRNA genes are consistent with the phylogenetic clustering (Fig. 1B). Within each clade, both ANI and 16S rRNA gene identities are greater than the level often used to delineate a bacterial species at 95.0% [52] and 98.7% [42], respectively. Between clade-M and clade-S, ANI is lower than 89.9% (Table 1, Fig. 1B), but the average 16S rRNA gene identity is greater than 99.7% (Table 1, Fig. 1B). Considering that the 16S rRNA genes may not provide sufficient resolution at the species level [53], we hypothesized that each clade may represent a distinct species and that the speciation process may have already completed. We tested this hypothesis as follows.

**Table 1.** Genetic variation of the *Ruegeria* population and the *Rhodobacteraceae* population.

| Population | Clade | #strains | 16S rRNA gene identity<br>(mean ± sd) | ANI<br>(mean ± sd) | #SNP/Mbp | #SNP |
| --- | --- | --- | --- | --- | --- | --- |
| *Ruegeria* | clade-M | 6 | 100.00 ± 0.00 | 98.24 ± 1.32 | 36,976 | 175,214 |
| | clade-S | 6 | 99.96 ± 0.04 | 99.05 ± 0.57 | 16,606 | 75,989 |
| | outgroup | 8 | 99.52 ± 0.36 | 97.08 ± 0.94 | 131,482 | 564,323 |
| | clade-M versus clade-S | 12 | 99.76 ± 0.03 | 89.67 ± 0.1 | 106,078 | 502,661 |
| | clade-M versus outgroup | 14 | 98.00 ± 0.25 | 83.04 ± 0.16 | 116,843 | 553,672 |
| | clade-S versus outgroup | 14 | 98.10 ± 0.25 | 83.18 ± 0.21 | 117,499 | 537,689 |
| *Rhodobacteraceae* | - | 214 | 100.00 ± 0.00 | 99.99 ± 0.64 | 10 | 43 |

### Differentiation of the Ruegeria population along coral compartments at the core genome

The branching pattern of a phylogenetic tree cannot always represent the population structure as the rate and influence of recombination may significantly affect the evolution of bacterial species [43]. To elucidate whether the population subdivision occurred between clade-M and clade-S, fineSTRUCTURE v2.0.7 [45] was applied to the core genome alignment of the two clades. As a result, 12 strains from clade-S and clade-M were assigned to seven subpopulations, three in clade-S and four in clade-M (Fig. 1C). The proportion of co-ancestry shared within each clade is much greater than that between clades (Fig. 1C), suggesting a strong barrier to homologous recombination between the two clades. The overall consistent branching pattern between the coancestry populations and the phylogenetic groups indicates that clade-M and clade-S diverged into two species. The decreased gene flow between clades is also supported by decreased *ρ/θ* ratio and r/m ratio between the two clades compared to those within clades (Supplementary Text 2.1, Table S1). Besides, F_st_, a measurement of population differentiation, was estimated across the core genome, and the genome-wide high F_st_ values (≥ 0.5) revealed that the speciation between the two clades is approaching completion (Fig. S2, Supplementary Text 2.1).

Recombination at the core genomic regions with distant lineages can be a key mechanism driving the differentiation of the *Roseobacter* population [44]. Such allelic replacements are expected to leave a strong signature of nucleotide substitution rate at the largely neutral synonymous sites (*d_S_*) in the affected genes [54–56], manifested as unusually large *d_S_* values at those loci for between-clade comparisons. We have recently developed a pipeline to pull out those loci through a clustering of the *d_S_* values between all possible pairs of genomes and across all shared single-copy gene families (Fig. S3, Supplementary Text 1.6) [48]. This procedure led to the identification of an outlier cluster consisting of 167 single-copy gene families each showing unusually large between-clade *d_S_* values (Table S2, Fig. 2). Novel allele replacement at these loci likely occurred at the last common ancestor (LCA) of either clade-M (marked with a blue triangle in Fig. 1A) or clade-S (marked with a pink triangle in Fig. 1A), and the inferred evolutionary history of allelic replacements was discussed in the following sections (Fig. S4, Fig. S5-S11).

**Figure 2.**
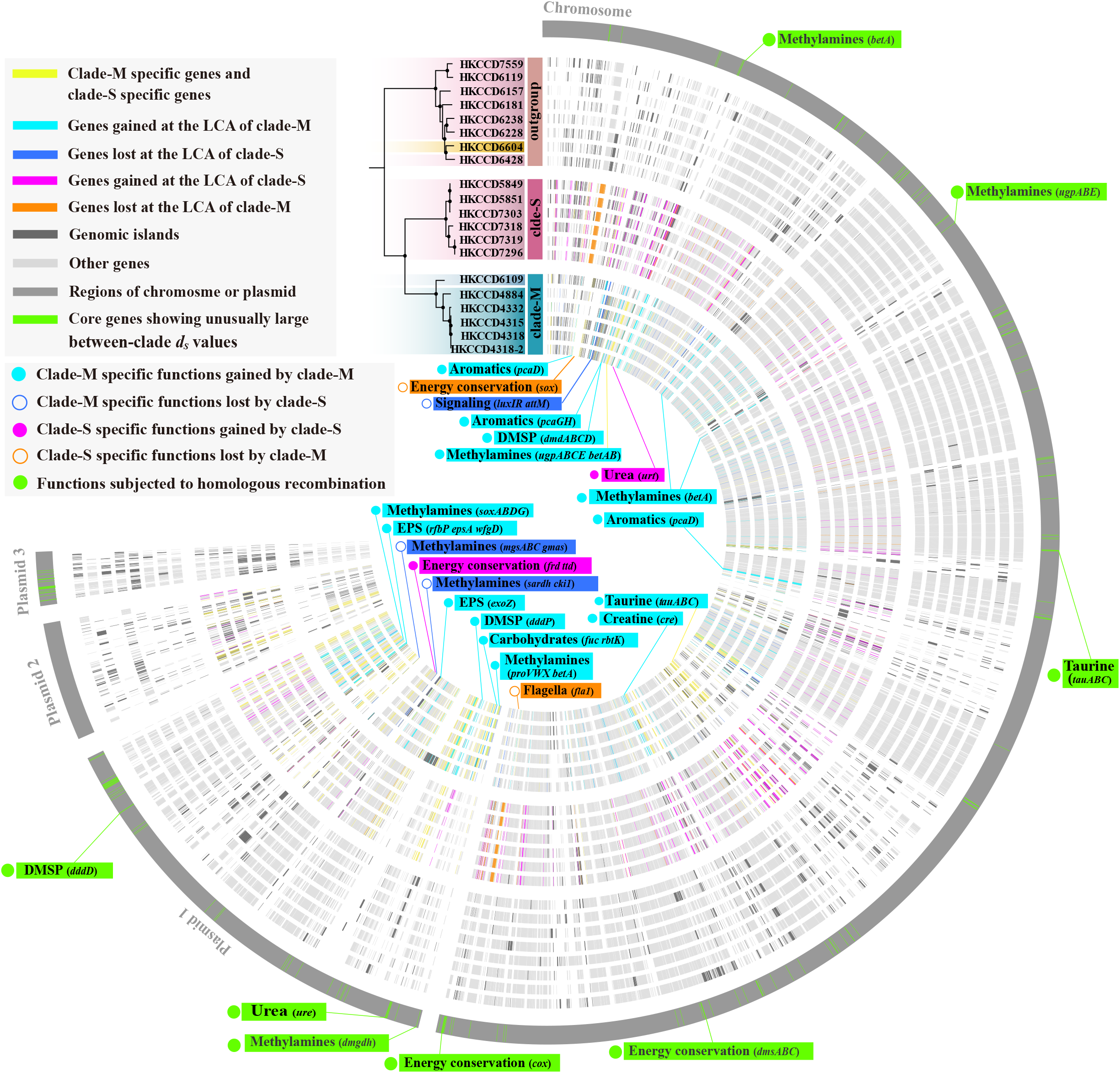
The genomic differentiation of the *Ruegeria* population. **(A)** The phylogeny and pangenome of the *Ruegeria* population generated using Circos v0.64 [103]. The gene families are arranged in order of the closed genome of strain HKCCD4315. The circular tracks depict the genomes which are arranged in order of the phylogeny and shown in grey. From the inner to the outer circle: (1)-(6) six genomes of clade-M. The genes gained at the LCA of clade-M are shown in cyan, and those lost at the LCA of clade-S are shown in blue. (7)-(12) Six genomes of clade-S. The genes gained at the LCA of clade-S are shown in magenta, and those lost at the LCA of clade-M are shown in orange. (13)-(20) Eight genomes of outgroup. The 536 clade-M specific genes and the 365 clade-S specific genes are shown in yellow in all tracks. The genomic islands (GIs) are showed in dark grey in all tracks. Other genes are shown in light grey in all tracks. (21) The genomic region of the chromosome and three plasmids are shown in grey, and the core genes showing unusually large synonymous substitution rate (*d_S_*) are shown in green.

### Differentiation of the Ruegeria population along coral compartments at the accessory genome

The above analyses demonstrated the genetic separation of clade-M from clade-S across the core genome. We further investigated whether population differentiation also occurred in the accessory genome (i.e., genes not shared by all members of the clade-M and clade-S). The clustering dendrogram based on the gene presence/absence pattern (Fig. 1D) showed that the similarity of accessory gene content was higher within each clade than that between the two clades. The congruent branching pattern between the gene content dendrogram (Fig. 1D) and the core gene-based phylogenomic tree (Fig. 1A) suggests that population differentiation has occurred at the accessory genome. Among the 3,202 accessory gene families, 194 and 157 families are universally and exclusively present in clade-M and clade-S members, respectively. Using a relaxed definition of “clade-specific gene families” where genes are present in at least two-thirds of the strains in one clade but present in no more than one-third of the strains in the other clade, we found 536 and 365 gene families specific to the clade-M and clade-S (Table S3, Table S4 and Fig. 2), respectively, suggesting that those two clades have diversified in many clade-specific functions.

Guided by the above principles, we inferred the evolutionary history of the clade-specific gene families. For the 536 clade-M specific genes, 224 and 64 gene families were inferred to be acquired at the LCA of clade-M and lost at the LCA of clade-S, respectively (Table S3). Since the basal position of the clade-M is occupied by a seawater strain (Fig. 1A), we further inferred that 122 additional gene family gains occurred at the LCA (marked with a red star in Fig. 1A) of the remaining five members which are exclusively mucus-associated (Table S3). For the 365 clade-S specific genes, 233 and 97 gene families were inferred to be gained at the LCA of clade-S and lost at the LCA of clade-M, respectively (Table S3). Thus, both gene gains and losses likely played an important role in differentiating the accessory genome. Among the 901 clade-specific gene families, 126 are associated with genomic islands (GIs) (Fig. 2), suggesting that GIs likely facilitated population differentiation. In the following sections, we elaborated on some clade-specific gene functions that may be of particular interest in terms of their potential roles in facilitating population differentiation.

### Metabolic adaptation of the mucus population to the eutrophic mucus niche

#### Utilization of methylamine-related osmolytes

A series of chemically related methylamines, such as glycine betaine (GBT), choline-O-sulfate (COS), glycerophosphocholine (GPC), choline and sarcosine, are compatible solutes and detected as cellular components of cnidarians including corals [57, 58]. Among these, GBT is the most prevalent osmolyte in corals [59]. GBT varies from 19 to 130 mmol/L in concentration among different coral species [58] and represents up to 90% of detected compatible solutes or 16% of the total N storage in some GBT-rich corals [10, 59]. Another two methylamines, trimethylamine N-oxide (TMAO) and dimethylglycine (DMG), are also marine osmolytes that can be transformed from choline and GBT (Fig. 3) [9], but have not been reported in coral studies.

**Figure 3.**
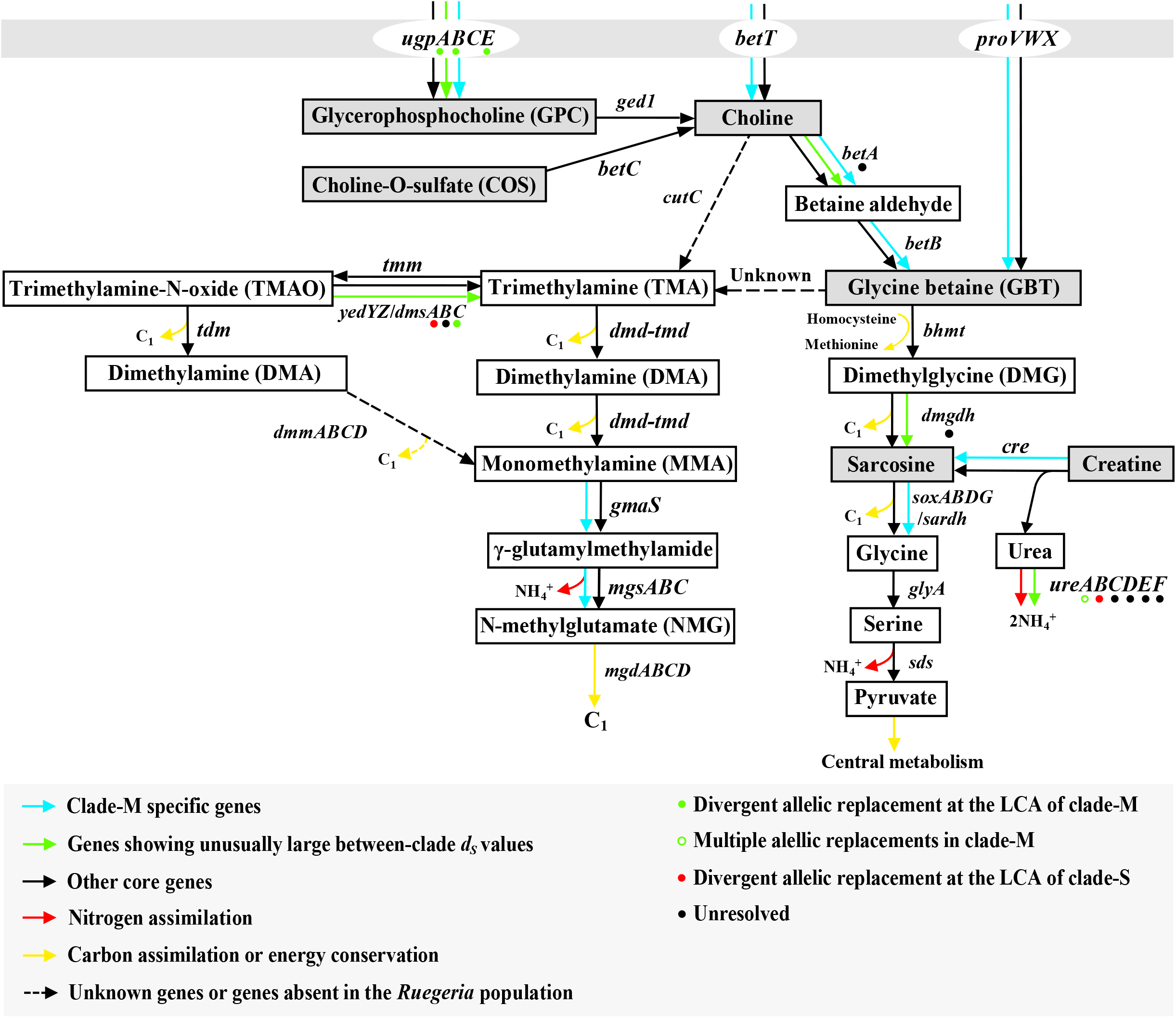
The catabolic pathways of methylamine-related coral osmolytes in the *Ruegeria* population. The compounds in grey blocks are osmolytes in coral. The solid arrows represent the genes present in the *Ruegeria* population. The arrows in dashed lines represent the genes missing from the *Ruegeria* population. The cyan arrows represent the clade-M specific genes. The green arrows denote the outlier core genes showing unusually large between-clade *d_S_* values. Divergent allelic replacement at the LCA of clade-M and clade-S are marked with solid green and red circles, respectively. The open circle in green represents multiple allelic replacements within clade-M. The black circles indicate that the allelic replacement history cannot be resolved based on the available data. The black arrows show the core genes shared by all clade-M and clade-S members. The red arrows indicate the reactions related to nitrogen assimilation. The yellow arrows indicate the reactions related to carbon assimilation. There are no clade-S specific genes found in these pathways. Only the dominant substrates of the promiscuous transporters are shown in the figure. The conversion from GBT to TMA was previously revealed by a metabolomics study on intestinal microbiota [67], but the underlying gene for this reaction has not been known.

Many of the clade-M specific genes are likely involved in the utilization of these methylamines. The methylamines may be transported into the bacterial cytoplasm through multiple transporters. For example, GBT may be taken up through the ABC transporter ProVWX [60]. The GBT precursors, GPC and choline, may be assimilated through the ABC transporter UgpABCE [61] and the BCCT transporter BetT [62], respectively. Clade-M members specifically contain an extra copy of all three transporters (Table S3, Fig. 3). After transportation into the cytoplasm, the metabolic pathways for GBT, GPC and choline are largely overlapped (Fig. 3). Choline is either incorporated into phosphatidylcholine (PC) or subjected to demethylative degradation [63, 64]. For the former, the key gene encoding choline kinase (*cki1*, Table S3) for assimilation is specific to clade-M. For the latter, the choline is oxidized to GBT aerobically before demethylation (Fig. 3). The key genes including the choline dehydrogenase (*betA*, Table S3) and betaine aldehyde dehydrogenase (*betB*, Table S3) are part of the clade-M specific genes (Fig. 3).

The model strain of the *Roseobacter* group, *Ruegeria pomeroyi* DSS-3, was previously shown to degrade GBT and produce intermediate metabolites including DMG, sarcosine and glycine sequentially [64]. This canonical pathway allows roseobacters to use GBT as a C, N and energy source (Fig. 3, the right-hand pathway located downstream of GBT) [64]. It starts from the demethylation of GBT to DMG catalyzed by betaine-homocysteine methyltransferase (*bhmt*) (Fig. 3) [64]. In the present *Ruegeria* population, however, a truncated *bhmt* gene (HKCCD4315_03106) which lacks the vitamin B_12_-binding domain is present in the core genome [65]. This truncated *bhmt* was also identified in the *Roseobacter* group member *Phaeobacter inhihens* (previously known as *Phaeobacter gallaeciensis*), and was considered nonfunctional for this reaction as the bacterium poorly used GBT as a C source [66]. Likewise, none of the six isolates assayed here were able to thrive on choline or GBT as a sole C and N source (Fig. 4A-2, 4A-3) or as a sole C source (Fig. 4B-3, 4B-5). When provided with the intermediate metabolites downstream of GBT, such as DMG (Fig. 4A-4) and sarcosine (Fig. 4A-5), these isolates thrived, further supporting that the utilization of GBT is blocked at the step catalyzed by BHMT. Specifically, DMG is demethylated to sarcosine through dimethylglycine dehydrogenase (*dmgdh*), and sarcosine can be demethylated to glycine through either sarcosine dehydrogenase (*sardh*) or sarcosine oxidase (*soxABDG*). Sarcosine also connects to another coral osmolyte creatine [57]; while creatine is not a downstream metabolite of GBT, it is hydrolyzed to sarcosine and urea through creatinase (*cre*) and the products follow the sarcosine demethylation pathway and ureolysis (*ureABCDEF*), respectively (Fig. 3).

**Figure 4.**
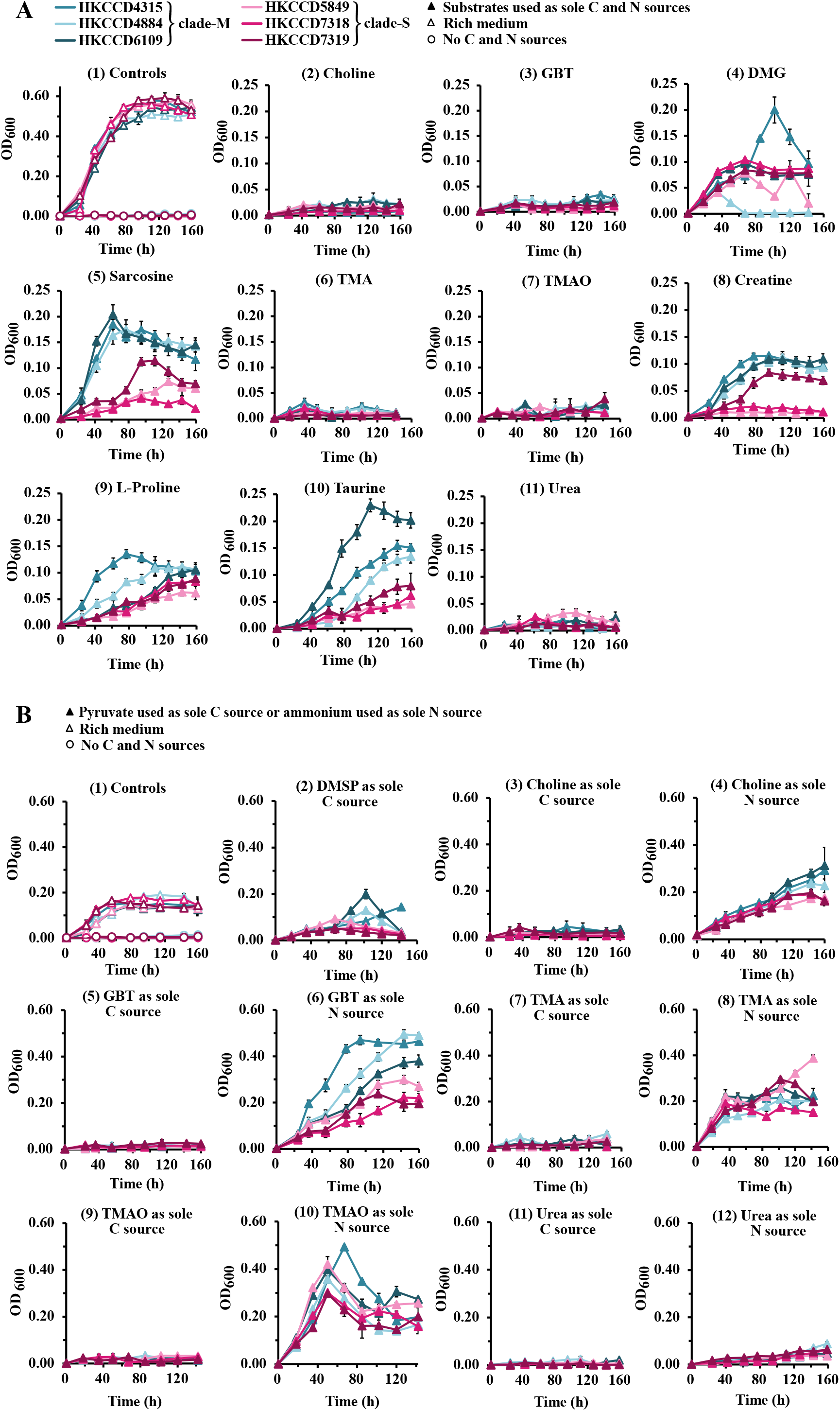
Growth experiments of three clade-M strains and three clade-S strains. **(A)** The minimal medium is supplemented with different substrates. (1) Control experiments. In the negative control, bacteria are cultured in the minimal medium without C and N source (open circles). In the positive control, bacteria are cultured in the rich medium, in which the peptone, glucose and yeast extracts are added as mixed C and N sources (open triangles). Growth experiments of the six strains on 5 mM of choline (2), GBT (3), DMG (4), sarcosine (5), TMA (6), TMAO (7), creatine (8), L-proline (9), taurine (10) and urea (11), each used as a sole C and N source (solid triangles). Three replicates are performed for each condition and error bars denote standard deviation. **(B)** Different substrates are used as either a sole C source or a sole N source. (1) Control experiments. In the negative control, bacteria are cultured in the minimal medium without C and N source added (open circles). In the positive control, bacteria are cultured using pyruvate as the C source and ammonium as the N source, respectively (open triangles). Growth experiments of the six strains on 5 mM of DMSP (2), choline (3), GBT (5), TMA (7), TMAO (9) and urea (11) as the sole C source and each with ammonium (10 mM) as the N source (solid triangles). Growth experiments of six strains on 5 mM of choline (4), glycine betaine (6), TMA (8), TMAO (10) and urea (12) as the sole N source and each with pyruvate (5 mM) as the C source (solid triangles). Three replicates are performed for each condition and error bars denote the standard deviation.

On the other hand, all assayed isolates can use choline (Fig. 4B-4) and GBT (Fig. 4B-6) as sole N sources, suggesting that they may acquire N from choline and GBT through an alternative pathway. One possibility is that GBT is initially transformed to trimethylamine (TMA) which can be used as a sole N source (Fig. 4B-8). This reaction was experimentally demonstrated in a metabolomics study on intestinal microbiota [67], but the genetic basis has not been known (Fig. 3). TMA can be sequentially demethylated to dimethylamine (DMA) and monomethylamine (MMA) through dimethylamine/trimethylamine dehydrogenase (*dmd-tmd*) [68]. Then MMA is further oxidized to release ammonium via a pathway comprising the gamma-glutamylmethylamide synthetase (*gmaS*), N-methylglutamate synthase (*mgsABC*), and N-methylglutamate dehydrogenase (*mgdABCD*) [69]. TMAO is another methylamine osmolyte that is chemically related to choline. It can be reduced to TMA through either anaerobic dimethyl sulfoxide/trimethylamine oxide reductase (*dmsABC*) [70] or aerobic methionine sulfoxide reductase (*yedYZ*) [71], and then enters the TMA demethylation pathway (Fig. 3).

While the above-mentioned genes involved in methylamine-related osmolytes are common to all members of the *Ruegeria* population, many of them have at least one additional copy specific to the clade-M members. The latter includes *ugpABCE* for GPC uptake, *betABT* and *cki1* for choline acquisition and catabolism,*proVWX* for GBT uptake, *cre* for converting creatine to sarcosine, *sardh* and *soxABDG* for sarcosine demethylation, and *gmaS* and *mgsABC* for MMA oxidation (Fig. 3). The majority of these genes were inferred to be acquired at the LCA of clade-M (12 out of 23) or lost at LCA of clade-S (four out of 23), while the remaining genes were acquired at the LCA of the five clade-M mucus members (seven out of 23) (Table S3). Among the core gene copies, the *ugpABE*, *betA*, *dmgdh*, and *dmsABC* were subjected to novel allelic replacements, though different component genes in each gene cluster went through different evolutionary history of homologous recombination (Fig. 3, Table S5). In line with these between-clade differences at the genetic level, the clade-M exhibited a significantly higher ability than clade-S to utilize the abundant osmolytes in corals, such as using choline (*p* < 0.05, One-way Repeated-Measures ANOVA; the same test used below unless stated otherwise; Fig. 4B-4) and GBT (Fig. 4B-6) as a sole N source, and using sarcosine and creatine as a sole N and C source (Fig. 4A-5 and 4A-8). While the osmolytes less common in corals (e.g. DMG, TMA and TMAO) were also used by the *Ruegeria* population, no significant differences between clade-M and clade-S were observed (Fig. 4A-4, 4B-8 and 4B-10). Detailed results of the physiological assays can be found in Supplementary Text 2.2.

To sum up, the above-mentioned genes were related to the metabolism of methylamine-related osmolytes, and most of them underwent either gene copy number variation or novel allelic replacements at the LCA of clade-M or clade-S. We therefore propose that the acquisition of genetic traits in utilizing methylamine-related coral osmolytes contributed to the diversification of clade-M from clade-S.

#### Utilization of other osmolytes in coral

Here, we highlighted three additional coral osmolytes that differentiated the clade-M from the clade-S: L-proline, DMSP and taurine [10, 58, 72]. L-proline is a potential substrate of the transporters ProVWX and BetT with relatively lower affinity compared to choline and GBT, respectively [60, 62]. This is supported by the growth assay (Supplementary Text 2.2), in which the clade-M members showed significantly higher growth yields and growth rates in utilizing L-proline than the clade-S members (Fig. 4A-9). DMSP can be degraded via either demethylation or cleavage pathway, yielding methanethiol (MeSH) and dimethyl sulfide (DMS) as the final products, respectively [73]. The demethylation pathway is catalyzed by *dmdABCD*, which was found exclusively in clade-M members and inferred to be acquired at the LCA of clade-M (Table S3). The cleavage pathway is mediated by a number of non-homologous genes, among which *dddP* (Table S3, Fig. 2) is exclusively present in the plasmid of clade-M members and *dddD* (Table S5) is part of the outlier core genes showing unusually large between-clade *d_S_* values, though the donor of the novel *dddD* allele is unclear based on our phylogenetic analysis (Fig. S4 versus Fig. S6). Using DMSP as a sole C source, the clade-M members accumulated a higher growth yield than the clade-S members (Fig. 4B-2). In terms of taurine, the genes encoding the taurine transporter (*tauABC*) are part of the outlier core genes that show unusually large between-clade *d_S_* values (Table S5). Phylogenetic analyses showed an independent evolutionary history of the three-component genes (Fig. S4 versus Fig. S7). For example, the *tauC* phylogeny suggests that the LCA of the five clade-M mucus members (marked with a red star in Fig. 1A) was subjected to divergent allele replacement, whereas the *tauA* phylogeny indicates that the seawater strain (HKCCD6109) and clade-S were the recipients of novel alleles. Notably, a second copy of the *tauABC* genes was found in four out of the six clade-M members (Table S3). These genetic differences are again supported by the growth assays, in which the clade-M members showed significantly higher growth yields in the utilization of taurine as a sole C and N source compared to the clade-S members (Fig. 4A-10). Most of the above-mentioned genes were acquired at the LCA of clade-M (7 out of 11) and the remaining four genes were acquired at the LCA of four clade-M members (Table S3), further supporting the utilization of coral osmolytes may be the ecological adaption of clade-M bacteria.

In addition to osmolyte utilization, the clade-M also acquired genes related to using other mucus related substrates (i.e. carbohydrates and aromatic compounds, Supplementary Text 2.3), as well as genes involved in quorum sensing (QS) system and biofilm formation (Supplementary Text 2.4), which may further facilitate the clade-M members to adapt to the nutrient-rich and densely-populated coral mucus habitat.

### Metabolic adaptation of skeleton population to anoxic and low-energy skeleton niche

#### Energy conservation in the periodically anoxic skeleton

Members of the *Roseobacter* group commonly conserve energy through the oxidation of inorganic reduced sulfur and carbon monoxide through the *sox* and *cox* gene clusters, respectively [74]. We identified a complete *sox* gene cluster in all members of the clade-S but in only one strain (HKCCD4884) of the clade-M (Table S4), and inferred that this *sox* gene cluster was lost at the LCA of clade-M. Additionally, two *cox* gene clusters encoding the form I (HKCCD4315_03676-03684) and form II (HKCCD4315_03987-03970) carbon monoxide dehydrogenase (*codh*) are shared by both clades. Among these, the form I *cox* gene cluster (Table S5), which is indispensable for carbon monoxide oxidation [75], showed unusually large between-clade *d_S_* values, suggesting the different genomic origins of carbon monoxide oxidation between the two clades. The phylogenetic trees of the component genes comprising the form I *cox* cluster consistently showed that the LCA of clade-S was subjected to divergent allele replacements at the *cox* gene cluster (Fig. S4 versus Fig. S8). Together with the presence of *sox* gene cluster, the clade-S members appear to use more versatile energy conservation strategies to adapt to the energy-limited coral skeleton niches.

The dissolved oxygen produced by the photosynthesis of symbiotic and endolithic algae during daytime is continuously consumed by the holobiont members through respiration [15], making the skeleton a diurnally anoxic environment [76]. Consistently, the clade-S acquired a gene cluster (*frdABCD* and *ttdAB*, Table S4) at LCA which likely enables the skeleton-associated strains to respire fumarate and L-tartrate under anoxic conditions. Besides, a gene cluster allowing for anaerobic respiration of dimethyl sulfoxide (DMSO) and TMAO showed an unusually large *d_S_* value (Supplementary Text 2.5, Fig. S9). These compounds could serve as the terminal electron acceptors for energy conservation in the diurnally anoxic skeleton [77].

#### Motility

Bacteria can be motile in multiple ways, among which the swimming motility enables bacteria to move in the liquids, while swarming and twitching allow the bacteria to be motile on solid surfaces (e.g., coral skeleton) [78]. In the *Roseobacter* group, the swimming motility is widely observed, but swarming motility has not been reported [79, 80]. Both swimming and swarming may be conferred by any of the three evolutionarily related flagellar gene clusters (FGCs), termed *fla1, fla2* and *fla3* [79, 80]. The putative twitching motility resulting in a dendritic-shaped phenotype was observed in seven *Roseobacter* species, but the genetic basis has not been clear [79].

All clade-S members except for HKCCD7318 carry the complete set of *fla1* consisting of 36 genes (Fig. 5A). In contrast, two DNA segments encompassing 17 continuous genes and part of the *flgI* gene (Table S4) are missing from the clade-M members (Fig. 5A), and they were inferred to be lost at the LCA of clade-M. These lost genes are involved in the assembly of multiple components of the flagellum, including components of type III secretion system (*flhA, flhB*, *fliQ*, and *fliR*), P- and L-rings (*flgA* and *flgH*), motor proteins (*motB*), basal body proteins (*flgB*, *flgC*, *flgG* and *fliI*), hook (*flgE*, *flgK*, *flgL* and *fliE*) and flagellum-specific ATP synthase (*fliI*). The strain HKCCD7318, an exception of the clade-S members, also carries a truncated *fla1* cluster, but encompasses five more genes encoding components of type III secretion system and basal body compared to the clade-M members (Table S4, Fig. 5A). Our experimental assays showed that the clade-S members with a complete set of *fla1* (HKCCD5849 and HKCCD7319) displayed larger swimming circles than the clade-M members and the HKCCD7318 which carried a truncated *fla1* cluster (Fig. 5B). No swarming or twitching motility was observed in all tested members (Fig. 5B). Previous studies on *Rhodobacter sphaeroides* showed that the expression of the *fla1* cluster is positively regulated under anaerobic conditions which could be part of the aerotaxic response of this bacterium to oxygen or alternative electron acceptors (e.g., DMSO and TMAO) in response to the anoxic condition [81]. This helps explain why the complete *fla1* in some clade-S members is maintained. As mentioned above, the coral skeleton becomes periodically anoxic, and the maintained motility may facilitate scavenging oxygen and alternative electron acceptors in the skeleton with compacted aragonites.

**Figure 5.**
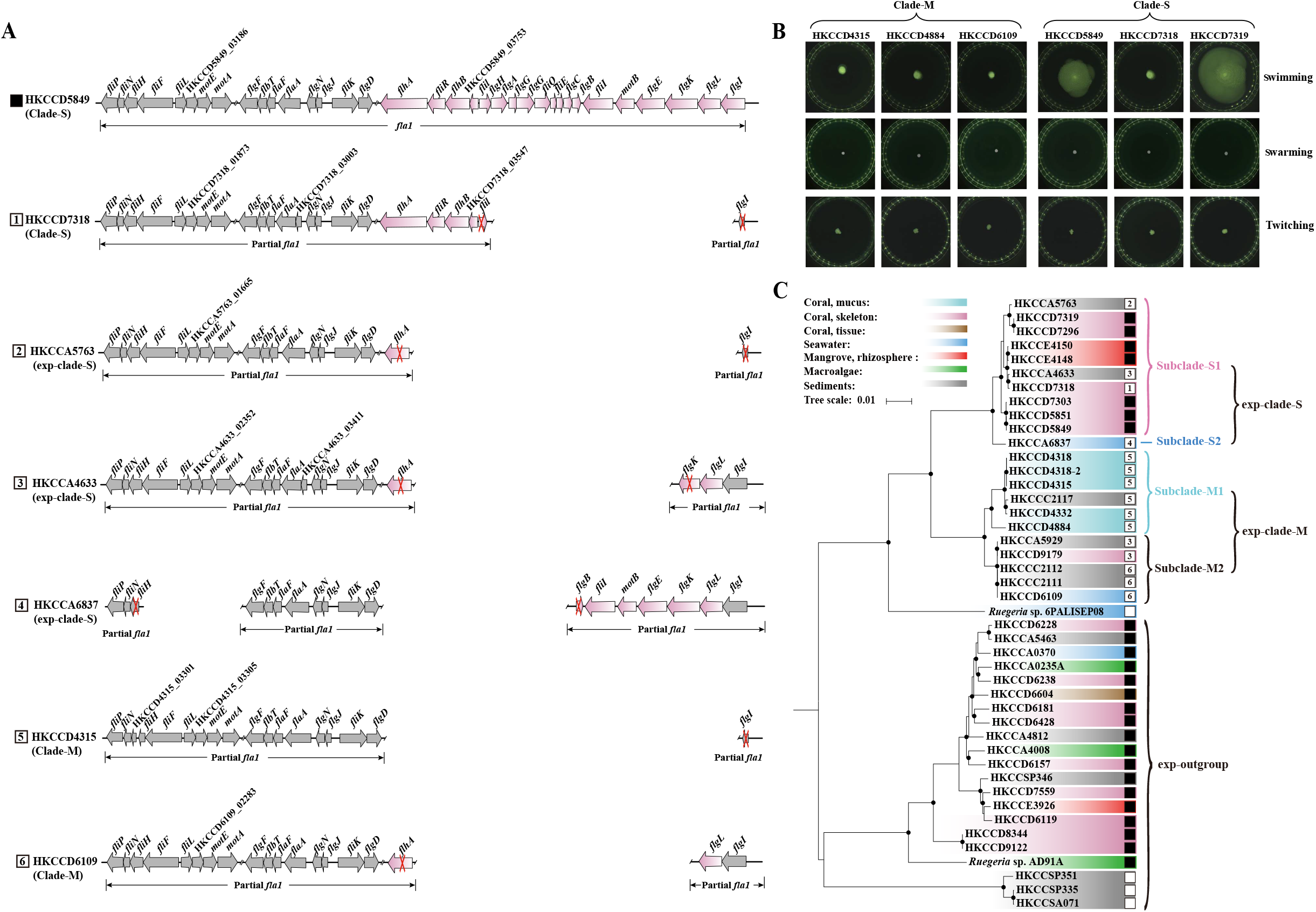
The phylogenetic distribution and pseudogene characterization of the flagellar gene cluster *fla1* across the expanded *Ruegeria* population composed of isolates from both coral and non-coral marine habitats, along with the motility assays of select isolates. **(A)** The gene arrangement of the *fla1* clusters in members of exp-clade-M and exp-clade-S. The pink arrows represent the clade-S specific genes, and the grey arrows denote core genes shared by exp-clade-M and exp-clade-S, respectively. The pseudogenes are marked with red crosses on the arrows. **(B)** The motility assays of three clade-M strains and three clade-S strains. Swimming and swarming motilities are tested on 0.3% (w/v) soft agar and 0.6% (w/v) agar for eight days, respectively; twitching motility is tested on 1.0% (w/v) agar in the humidified box for 10 days. **(C)** An expanded genome phylogeny of the *Ruegeria* population based on 44 strains including those collected from various ecological niches. The exp-clade-M, exp-clade-S and exp-outgroup expand from the clade-M, clade-S and outgroup in Fig. 1A. Solid circles at the nodes indicate that the ultrafast bootstrap support value of the branch is 100%. The scale bar indicates the number of substitutions per site. The root of the tree is determined using a larger phylogenomic tree (Fig. S12) composed of 454 *Ruegeria* and related strains [50]. The strains containing the complete *fla1* cluster are marked with black boxes. The strains with partial *fla1* clusters are marked with white boxes, and different types of partial *fla1* clusters in exp-clade-M and exp-clade-S are distinguished by the numbers in the white boxes.

#### Possible role of the urea remineralization potential

Coral reef ecosystem is generally poor in N [82]. Urea, mainly from the excretion of zooplankton, fish and other animals, is an important source of N [82, 83] for the coral host [84, 85], *Symbiodiniaceae* [85] and bacteria [86] in the coral holobiont. Urea is hydrolyzed to ammonium and inorganic carbon by urease and results in the rise of pH and carbonate concentration [87]. Ureolysis by bacteria is known to facilitate calcification by increasing pH and supplying carbonate in environments such as soil, estuarine, coastal seawater, animal rumen and gut [88–91]. In the coral skeleton, the calcification process can also be enhanced by ureolysis [92, 93].]

A gene cluster encoding the urea transporter (*urtABCDE*, Table S4) is specific to the clade-S, and four (*urtABCD*) out of the five subunits were likely lost at the LCA of clade-M (Table S4). Besides, the genes (*ureABCDEF*, Table S5) encoding the urease for urea hydrolysis are part of the core genes showing unusually large between-clade *d_S_* values. Phylogenetic analyses suggested that either clade-S or clade-M acquired alleles from a different genetic origin, depending on the component gene (Fig. S4 versus Fig. S10). These results suggest that the clade-M and clade-S have differentiated on the urea decomposition, and the clade-S may show a higher potential to take advantage of this function as it kept the transporter for urea uptake. In that case, the conservation of genetic traits in urea mineralization by skeleton-associated members might be beneficial for the calcification of the coral host.

Interestingly, our growth assay showed that both clade-M and clade-S members grew poorly on urea under the tested conditions and did not exhibit differential growths (Fig. 4A-11, 4B-11 and 4B-12). We argue that generating N source is not the primary purpose of the *Ruegeria* population to use urea. This is because urea may not be a preferred N source of bacteria due to the high energy cost in using it [94], especially when the coral skeleton is enriched with dissolved inorganic nitrogen with its concentration exceeding that in ambient seawater by a factor of 10 [95]. Besides using urea as N source [96], bacteria also benefit from ureolysis for survival in acidic environments [97] and chemotaxis to the host [98], which are induced by acid exposure [97] and enrichment of nickel (Ni) [99], respectively. However, the environmental inducers of urease expression (i.e., acidic pH and high level of Ni) were not screened in the present study, which might explain the overall poor utilization of urea by the *Ruegeria* population and the absence of between-clade differences. We hypothesize that ureolysis in the skeleton may be a conditionally expressed function in response to undetected environmental inducers, which may increase the calcification efficiency of coral host meanwhile. Future experiments are needed to validate this hypothesis.

### The role of different coral compartments in driving the Ruegeria population evolution

The metabolic differences between the clade-M and clade-S provide a wealth of evidence that the diversification of the two clades is likely driven by heterogeneous conditions associated with different coral compartments. However, the splitting time of the two clades was estimated to be 7.52 million years ago, indicating that the split of clade-M and clade-S far predated the birth of their host coral which likely emerged several decades to a century ago [100]. The long evolutionary time thus leaves the origin of the clade-M and clade-S an unanswered question: did they evolve *de novo* on coral, or had they already occurred in other environments before their colonization on the different compartments of the coral host?

In the former case, different compartments is the direct force driving speciation. The bacteria have a chance to undergo a long evolution time on corals even beyond the longevity of coral, because bacteria could be transmitted vertically between coral generations (Fig. S11) [101]. In this model, the LCA of the clade-M and clade-S colonized the ancient coral host, then the clade-M and clade-S diverged *in situ* through the acquisition or loss of metabolic traits and thrived in their preferred coral compartments along with the growth cycle of the coral host. The predominant bacterial clade in one compartment might still be able to survive in another as they are spatially close, but with a relatively lower abundance due to compartment-specific selection for its own inhabitants. Next, members of both clades have a chance to be transmitted together with the coral gametes and the planula, and thrive in their optimum compartments after the coral recruit is settled and starts to secret mucus and develop the skeleton.

If the second hypothesis is true (Fig. S11), distinct coral compartments may not be the initial force driving the speciation between the clade-M and clade-S. Instead, the two clades might have already diverged in other environments and repeatedly colonized on different compartments of the coral host throughout their evolutionary processes. The environmental heterogeneity of different coral compartments might impose distinct selective forces on the localized bacteria and enrich the best-adapted clade members, and this in turn may accelerate the diversification of the two clades.

To help resolve these competing hypotheses, we included additional 24 strains that are closely related to members of the *Ruegeria* population. Among these, 22 co-occurred with the coral-associated *Ruegeria* population in the Hong Kong coastal ecosystems, but they were isolated from a variety of ecological niches including brown algal ecosystem niches (e.g. seawater, sediments and algal tissue), mangrove rhizosphere, coastal seawater, intertidal sediments and another batch of the *P. acuta* coral sample (Table S6). According to the updated phylogenomic tree (Fig. 5C), the newly added strains expanded all three major clades of the *Ruegeria* population (Fig. 5C). Members in the expanded clade-M (hereafter “exp-clade-M”) were equally partitioned into two subclades, in which subclade-M1 is dominated by the mucus-associated members (equivalent to the clade-M in Fig. 1A) and subclade-M2 consists of members largely from non-coral niches (Fig. 5C). This pattern prevented us from inferring the ancestral habitat of the LCA of exp-clade-M. Members in the expanded clade-S (hereafter “exp-clade-S”) were unequally grouped into two subclades, with the singleton subclade-S2 derived from seawater and the subclade-S1 composed of phylogenetically mixed strains inhabiting the coral skeleton and other non-coral niches (Fig. 5C). Thus, we cannot conclusively determine the habitat of the LCA of the exp-clade-S. In the case of the expanded outgroup clade, the coral-associated strains are embedded in the early-branching lineages that are not derived from corals (Fig. 5C), suggesting that the ancestral habitat of the LCA of this expand outgroup clade was not related to coral. When these three clades were viewed collectively, together with the deeply-branching singleton clade represented by a seawater strain *Ruegeria* sp. 6PALISEP08 located at the middle of the tree (Fig. 5C), the available isolates’ habitat information and phylogenetic structure are more consistent with the hypothesis that the original skeleton clade, mucus clade, and outgroup clade (Fig. 5C) each independently evolved from ancestors colonizing non-coral habitats.

Since the cladogenesis event between the original clade-M and clade-S was likely completed in habitats other than the coral compartments, some between-clade differences of metabolic potential could have existed before ancestral lineages independently transited to coral mucus and skeleton. We therefore sought evidence that allows singling out the metabolic differences imposed by different coral compartments from those by non-coral environments. In brief, we screened the gene families that show clade-M (or clade-S) specific distribution but are not prevalent in exp-clade-M (or exp-clade-S), because the evolution of these genes is more likely driven by coral compartments. We then inferred the evolutionary history of the eligible gene families along the genome-based phylogeny of the 44 strains (Fig. 5C).

This new analysis revealed that over half of the osmolyte utilization genes specific to clade-M discussed above were acquired at the LCA of the subclade-M1 in the expanded phylogeny (Table S7). These genes include *ugpABCE* for GPC uptake, *betAB* for choline oxidation, *gmaS* and *mgsABC* for MMA oxidation, *cre* for creatine degradation, *tauABC* for taurine uptake, and *dddP* for DMSP lysis. This is evidence that the coral mucus environment represents an important selective force shaping the genome of mucus-associated bacteria. This expanded analysis, however, rendered the evolution of the remaining osmolyte degradation genes specific to clade-M uncoupled from the coral mucus habitat, since these genes were prevalent among the exp-clade-M members and were largely acquired at the LCA of the exp-clade-M (Table S7). Similarly, the expanded analysis uncorrelated the evolution of urea transporter genes, sulfur oxidation genes, and fumarate and L-tartrate respiration genes to the coral skeleton habitat, since these genes were commonly found in the exp-clade-S members and were either acquired at the LCA of exp-clade-S or lost at the LCA of exp-clade-M (Table S8). In the case of the type I flagellar gene cluster (*fla1*) comprising 18 genes, it is difficult to infer its evolutionary history based on the gene presence and absence pattern. This is because while a few members in the exp-clade-S including most skeleton-associated members possess a complete set of the genes, the basal branch of this clade and all members of exp-clade-M each contain a subset of the genes (Fig. 5). In the strains carrying an incomplete *fla1* cluster (Fig. 5C), the genes adjacent to the missing part of the *fla1* were mostly pseudogenized (Fig. 5A). Because pseudogenization is unequivocal genetic evidence of ongoing gene loss [50], the missing part of *fla1* is more likely a result of gene loss (Table S8). Therefore, the conservation of a complete *fla1* cluster in most skeleton-associated members indicates that the coral skeleton may act as a selective force to maintain *fla1* in the clade-S.

### Limited migration of a genetically uniform Rhodobacteraceae population across coral compartments

For the *Ruegeria* population, we provided evidence that different compartments of the coral host likely act as distinct selective pressures that drive the evolution and adaptation of bacterial genomes. Given a considerable amount of genetic diversity harbored in that population, however, it remains unknown whether different compartments of the coral host also act as a physical barrier of gene flow among bacteria colonizing different compartments. We next investigated this question by using a genetically uniform *Rhodobacteraceae* population, which is composed of 214 isolates sampled from four different coral individuals of the same coral species *P. acuta* (Table S6), each collected from a different location in Hong Kong (Fig. S1). This new population represents a distinct lineage from all known *Roseobacter* group members (Fig. S4), showing identity of 97.7% at the 16S rRNA gene to its closest relative, *Rhodobacteraceae bacterium* MA-7-27 (GenBank Assembly Accession Number: GCA_003688285.1). Members of this population share identical 16S rRNA genes, show the pairwise ANI at ~99.99% (Table 1), and differ by only 43 SNPs across around 4.39 Mbp core genome sequences shared by the 214 strains according to kSNP3 [102] (Table 1). Phylogenomic construction based on these SNPs showed that the isolates are not clustered according to the coral compartments or the sampling locations (Fig. S13). The extremely high genetic identity among isolates indicates a very recent origin of this population within the past several hundreds of years [130 and 94 years ago for the Wong Wan Chau (WWC) and Ngo Mei Chau (NMC) subpopulations, respectively. Supplementary text 1.11], which likely spanned up to a few generations of the coral animals in Hong Kong waters [100] and thus provided a unique opportunity to test whether dispersal limitation may have a role in driving population differentiation.

The Slatkin-Maddison test [51] is designed to measure the extent of compartmentalization according to the number of migrations producing the observed phylogenetic placement (Supplementary Text 1.10). By assuming that the common ancestor of closely related strains occupied a single compartment, a migration event is inferred when a closely related strain is found in a different compartment. The small probability that the estimated number of migrations from the real data is greater than the expected number of migrations calculated from randomly generated population structures by permutating the strains across the phylogenomic tree reflects the high degree of compartmentalization. The WWC subpopulation (Fig. 6A) and the NMC subpopulation (Fig. 6B) each consist of two or more compartments, and thus are qualified for the Slatkin-Maddison test. This test showed that in both subpopulations fewer migrations were inferred than expected by chance (29 migrations in the WWC subpopulation, *p* < 0.01, Fig. 6C; 12 migrations in the NMC subpopulation, *p* < 0.001, Fig. 6D), indicating that members from distinct compartments of the same coral individual are compartmentalized and that microbial population differentiation along coral compartments likely started from limited migration between compartments.

**Figure 6.**
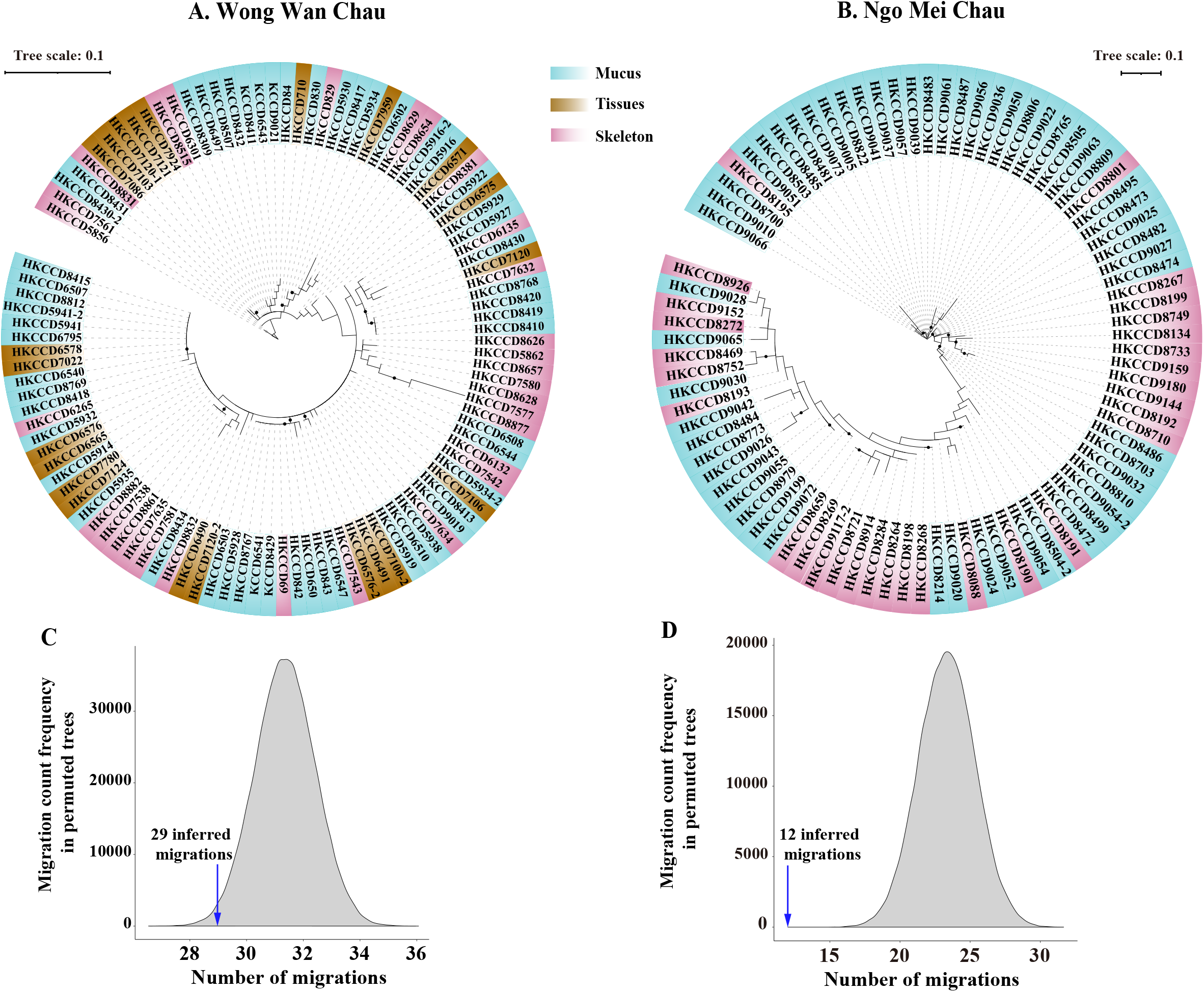
Compartmentalization of two subpopulations of the *Rhodobacteraceae* population each from a distinct coral individual. **(A-B)** Phylogenomic trees constructed by IQ-TREE v1.6.5 using the core SNPs identified with kSNP3 for the population associated with the coral individual collected at Wong Wan Chau and the other associated with the coral individual at Ngo Mei Chau. Strains isolated from different coral compartments are highlighted with different colours. Solid circles at the nodes indicate that the ultrafast bootstrap support value of the branch is ≥ 80%. The scale bar indicates the number of SNPs per variable site. **(C-D)** The frequency distribution of the number of migrations required to produce each of the 100,000 permuted trees in the two subpopulations. The blue arrow denotes the inferred number of migrations based on the real data.

## Concluding remarks

Coral hosts harbor multiple compartments that differ in many physicochemical properties, including nutrient quality and quantity, energy sources and redox levels among others. Here, we provided the first evidence that this heterogeneity drives the population evolution of bacterial residents. Of the two *Roseobacter* group populations examined here, one (the *Rhodobacteraceae* population) differs at only a few SNP sites across the whole genomes but has already shown dispersal limitation across the different compartments, whereas the other (the *Ruegeria* population) has already diverged into distinct species varying in multiple functional traits. For the latter, gene gains and losses at the accessory genome along with novel allele replacements at the core genome are likely the underlying processes driving their differentiation. While the clade-M acquired novel genes to make use of mucus-related compounds (e.g., methylamines and other coral osmolytes, carbohydrates and aromatics) and to enable them to compete well with other bacteria in the densely populated coral mucus; clade-S maintained metabolic traits lost by clade-M, such as oxidation of inorganic sulfur and carbon monoxide, motility, and potential urea decomposition, and also acquired new traits to distinguish themselves from the outgroup, which are of vital importance to survive in skeletons. Although some traits may have diversified before the colonization of the coral host, the differentiation in a few important traits including motility and utilization of mucus-derived compounds indicates skeleton and mucus environments may be important selective forces shaping the *Ruegeria* population evolution.

Previous coral microbiome studies successfully characterized the prevalence of *Rhodobacteraceae* members in coral environments at the community structure level, but the role of each lineage on corals, and more specifically, on different coral compartments has not been appreciated. Population genomics is a powerful tool to trace how specific lineages evolve along different coral compartments, and brings us the knowledge of how bacteria interact with coral through neutral processes due to dispersal limitation and adaptive choices of local groups. By taking advantage of population genomics, our analysis provides the first insights that evolution of the ecologically relevant *Ruegeria* population is driven by the specific environmental factors of distinct coral compartments. Some of the local adaptation strategies are win-win choices for the bacteria and coral host. For example, clade-M members may remove some potentially toxic aromatics trapped in the coral mucus, and clade-S members possess genetic potential to decompose urea, which might confer benefits to the coral host by promoting calcification of the skeleton when the environment acidifies. These predictions, if confirmed, would be the building blocks to further understand the whole nature of the bacteria-coral association and would help to improve our strategies in coral conservation by more carefully considering the role of bacteria in maintaining the health of coral holobiont.

## Acknowledgments

We thank Ryan Ho-Leung Tsang for the coral sample collection, Tsz-Yan Ng for guiding coral sample processing, Xiao Chu and Shuangfei Zhang for their help on bacteria cultivation, Hao Zhang and Minglei Ren for their help in data analysis, and Xinqin Lin for her advice in experimental design. We thank Xiao Chu, Minglei Ren and Zhichao Zhou for providing isolates from the brown algae ecosystem, sediments and mangrove ecosystem. This work was supported by the Shenzhen Science and Technology Committee (JCYJ20180508161811899), the Hong Kong Environment and Conservation Fund (15/2016), the Hong Kong Research Grants Council General Research Fund (14163917), and the Hong Kong Research Grants Council Area of Excellence Scheme (AoE/M-403/16).

## Conflict of Interest

The authors declare no competing commercial interests concerning the submitted work.

## Data Availability

The assembled genomic sequences and raw reads of the coral-associated *Ruegeria* population, the seven *Ruegeria* isolates from non-coral marine habitats, and the coral-associated *Rhodobacteraceae* population are made publicly available at NCBI under GenBank assembly accession number PRJNA596594, PRJNA682389 and PRJNA596592, respectively.

## Notes

### Competing Interest Statement

The authors have declared no competing interest.

