## Supplementary Information for "Population Differentiation of *Rhodobacteraceae* Along Coral Compartments"

This PDF file includes:

Text 1. Supplementary methods

Text 2. Supplementary results

Figures S1 to S13

Supplementary references

**Text 1. Supplementary methods**

1.1 Coral sample collection and processing

1.2 Bacterial isolation

1.3 Genome sequencing, assembly and annotation

1.4 Ortholog prediction and phylogenomic tree construction

1.5 Analysis of population structure in core genomes

1.6 Inference of novel allelic replacement with external lineages in core genomes

1.7 Differentiation in the accessory genome and inference of evolutionary history

1.8 Identification of pseudogenes in the *fla1* flagellar gene cluster

1.9 The physiological assays

1.10 Test of compartmentalization and dispersal limitation

1.11 Estimating the origin time for the *Rhodobacteraceae* and the *Ruegeria* populations

**Text 2. Supplementary results**

2.1 Population differentiation at the core genomes of the *Ruegeria* population

2.2 The *Ruegeria* population differentiation at the physiological level

2.3 Metabolic potential for utilizing other substrates by the mucus clade of the *Ruegeria* population

2.4 Metabolic potential of the mucus clade in the *Ruegeria* population underlying microbial interactions in the densely-populated mucus habitat

2.5 Adaptation of the skeleton clade in the *Ruegeria* population to the periodically anoxic skeleton habitat

**Text 1. Supplementary methods**

*1.1 Coral sample collection and processing*

Coral samples of *Platygyra acuta* were collected by SCUBA diving in Hong Kong water at Kiu Tsui Chau (N 22°22'04.4" E 114°17'42.0") on 24th April 2017, Wong Wan Chau (N 22°31'31.2" E 114°19'00.1") on 12th January 2018 and Ngo Mei Chau (N 22°31'47.2" E 114°19'02.9") and Chek Chau (N 22°30'03.3" E 114°21'22.7") on 25th February 2018 (Fig. S1A). One coral rubble (2-8 cm in diameter) was sampled from each colony using a rock chisel, separated in zip-lock bags with their ambient seawater, kept in a low-temperature oven, and carefully transported to the laboratory. One sample of ambient seawater was collected by 50 mL centrifuge tube at each site.

Separation of coral compartments followed an established procedure [1, 2]. In brief, coral fragments were washed three times with filtered ambient seawater for 10 seconds with stirring to disrupt the exogenous microbial contaminants from the ambient seawater or sediments. Mucus samples were collected by exposing coral fragments to the air in the clean bench and waiting until the mucus started to drip from the coral surface. A total of 150 μL dripping mucus was collected using sterile syringes and transferred to 1.5 mL sterile centrifuge tubes. The collected mucus was centrifuged at 2,000 rpm for five minutes. The cell debris on the bottom was discarded and the transparent supernatant was kept.

Tissue samples were collected by spraying the coral surface using a Waterpik. Tissue suspensions of 50 mL were collected with sterile zip-lock bags, and centrifuged at 12,000 rpm for 15 min under 4 °C. The pellet was then suspended in 1 mL of autoclaved artificial seawater (ASW). While procedures to collect clean mucus and skeleton were established [2], the accurate method for collecting clean coral tissue remains unavailable due to the intersecting structure of the coral compartments. For example, the mucocytes are part of the coral issue layer (Fig. S1B), which keeps secreting mucus [3]. Besides, the tissue is embedded in the corallites (Fig. S1B), which are part of the skeleton where the polyp sits and retracts, so the removal of tissue would inevitably disturb the coral skeleton [4]. These anatomical features make the complete separation of tissue from mucus and skeleton not possible by current methods, such as airbrush [5], Waterpik [1] and centrifugation [6].

The core coral skeleton pieces of ~2 cm in diameter were carefully separated. To avoid cross-contamination from the tissue, only the skeleton pieces located more than 2 cm apart from the tissue layer were kept. Then the skeleton pieces were crushed into a slurry with sterilized mortar and pestle with 1 mL ASW added. The slurry was filtered through a 100 μM mesh to remove large fragments.

*1.2 Bacterial isolation*

The collected coral compartments were serially diluted and immediately transferred to marine basal medium (MBM) agar plates. The MBM marine agar was prepared as the following recipe (per liter): 8.47g of Tris-HCl, 0.37 g of $NH_4Cl$, 0.0022 g of $K_2HPO_4$, 11.6 g of NaCl, 6 g of $MgSO_4$, 0.75 g of KCl, 1.47 g of $CaCl_2 \cdot 2H_2O$, 2.5 mg of FeEDTA [pH 7.5], 1 mL of vitamins [7], and 15g of agar. Taurine was added as the carbon source at the concentration of 0.5 mM. The ambient seawater was treated in the same way as the samples of coral compartments, serially diluted and spread over agar plates. Agar plates were incubated at 28 ℃ for at least 48 h. Colonies were randomly selected and subject to streaking three times on 2216E marine agar [BD Difco, USA] for purification.

The 16S rRNA gene was amplified using colony polymerase chain reaction (PCR) with 27F primer (5'-AGAGTTTGATCCTGGCTCAG-3') and 1492R primer (5'-GGTTACCTTGTTACGACTT-3'). By following the protocol, Chelex 100 resin [Bio-rad, USA] was used to prepare biomass samples, and the recipe of PCR was prepared using Premix Taq [Takara Bio, USA]. The PCR was performed according to the following procedure: denaturing at 95℃ for 5 minutes, followed by 32 cycles (95℃ for 45 seconds, 55℃ for 45 seconds and 72℃ for 90 seconds) and a final extension at 72℃ for 10 minutes. The amplicons were sequenced using 27F primer. The primers and the bases with low sequencing quality at the ends of the amplicons were removed, and the remaining 600 bp were kept. The taxonomic information was obtained by comparing the partial 16S rRNA gene sequences with those of all reported type strains using EzBioCloud [8]. The partial 16S rRNA gene sequences were clustered to form operational taxonomic units (OTUs) at the 98.7% identity level, which is used to delineate a bacterial species [9]. Two *Rhodobacteraceae* OTUs each containing 12 (the *Ruegeria* population) and 214 isolates (the *Rhodobacteraceae* population) covering two or more coral compartments were chosen for population genomic analyses. For the *Ruegeria* population, an additional closely related OTU with 8 strains was included as outgroup.

*1.3 Genome sequencing, assembly and annotation*

For each of the 234 isolates comprising the two populations, genomic DNA was extracted using TaKaRa MiniBEST Bacteria Genomic DNA Extraction Kit [Takara Bio, USA]. The quality of each extracted DNA sample was verified spectrophotometrically using NanoDrop™ 2000 [Thermo Fisher, USA] ($A_{260}/A_{280}$ >1.8, $A_{260}/A_{230}$ > 2.0 and $A_{260}$ >$A_{270}$). Whole-genome sequencing was performed using the BGISEQ-500 PE100 platform (Table S6) in Qingdao Huada

Gene Biotechnology Co., Ltd. The untrimmed adapters associated with raw reads were identified with BBMerge implemented in BBmap v37 [10]. Next, adapters and low-quality reads were trimmed using Trimmomatic v0.33 [11], reads each with less than 40 bp were discarded, and the quality of the remaining reads was checked with FastQC v.0.11.4 (https://www.bioinformatics.babraham.ac.uk/projects/fastqc/). Contigs were assembled based on the high-quality paired-end reads using SPAdes v3.9 [12] with default parameters. Only those with a length of over 1,000 bp and with a k-mer coverage over five were kept for further analyses. CheckM v0.9.7 [13] was used to assess the quality of assemblies, and statistics were calculated with QUAST v4.5 [14]. The genome of HKCCD6109 in the *Ruegeria* population showed a 50% heterogeneity (Table S6) by CheckM v0.9.7, suggesting potential DNA contamination from very close relatives. To check the potential contamination, we re-purified and re-sequenced the sample HKCCD6109 as described above. The new version of genome assembly of HKCCD6109 was estimated to have completeness of 99.7% and heterogeneity of 50%. The old and new version of the assembled genome size is 4,472,332 bp and 4,522,316 bp, respectively, and they differ at eight nucleotide sites across the aligned regions (3,595,109 bp). Four of these sites are located together, and the other four are located randomly on the chromosome. Apparently, differences at the former four sites cannot be ascribed to sequencing error, and the possibility that the old HKCCD6109 culture contains very closely related contamination cannot be ruled out. Note that our physiological assays (Supplemental Text 2.2) involving HKCCD6109 used the old version.

Gene prediction was carried out using Prokka v1.14.6 [15]. The functions of the predicted protein-coding genes were further annotated using NCBI Conserved Domain Database (CDD) [16], RAST Annotation Server [17], and eggNOG [18].

To obtain a closed genome as a reference for the *Ruegeria* population, PacBio Sequel was used to sequence the strain HKCCD4315 isolated from coral mucus. Unicycler [19] was used to assemble the complete genome of strain HKCCD4315 based on short reads from BGISEQ-500 PE100 and long reads from PacBio Sequel. The gene prediction and functional annotation were performed as described above.

*1.4 Ortholog prediction and phylogenomic tree construction*

Orthologous gene families were identified using OrthoFinder v2.2.1 [20] among the strains in each population. Members of each single-copy gene family shared by all tested strains were aligned at the amino acid sequence level using MAFFT v7.215 [21]. Gaps of alignments were trimmed using trimAl v1.4.rev15 with parameters "-automated1 -resoverlap 0.55 -seqoverlap 60" [22]. PartitionFinder2 v2.1.1 [23] was implemented to determine the best-fit evolutionary model for each family. The maximum likelihood phylogeny was constructed based on the concatenated sequences of trimmed alignments and model selection results using IQ-TREE v1.6.5 [24] with 1,000 ultrafast bootstrap replicates.

*1.5 Analysis of population structure in core genomes*

For the *Ruegeria* population, ANI for each genome pair was calculated using FastANI [25]. The whole-genome alignment of 12 strains comprising the clade-M and clade-S of was produced using progressiveMauve v2.3.1 [26] with default settings. The core genomic regions shared by the 12 strains were extracted. To measure the relative rate and effect of recombination relative to point mutation among the population, ClonalFrameML v1.1 [27] was implemented with the core genome alignments and the phylogenomic tree as inputs. The same analyses were conducted for members of clade-M and those of clade-S separately.

To infer the population subdivision, we carried out the coancestry analysis. We generated the haplotype data using SNPs from the core genomic alignments and the recombination map files following the instructions of ChromoPainter [28]. Chromosome painting was implemented to calculate the co-ancestry between pairwise strains using ChromoPainter. Next, the fineSTRUCTURE [29] assigned strains to subpopulations based on the co-ancestry matrix using a model-based clustering method. Assuming each individual as a recipient of DNA from the remaining individuals (i.e., donors), the “chromosome painting” algorithm reconstructs the genome of each recipient with chunks of DNA from other donors. Then the painting results were summarized as a “co-ancestry matrix”, which represents ancestral relationships among individuals. Based on the co-ancestry matrix, individuals were assigned to subpopulations by the Markov chain Monte Carlo (MCMC) algorithm in the fineSTRUCTURE. Both the burn-in and the MCMC step were run for 100,000 iterations to ensure convergence. The thin interval was specified as 100. Two independent inferences were performed with the same parameters to confirm the population assignment. The population structure was visualized with the R script “fineRADstructure.R” [30].

To provide further information about the population differentiation between clade-M and clade-S, we calculated the $F_{st}$ values using Arlequin v3.5 [31] with 1,000 permutations. SNPs within and between two clades were extracted and coordinated to the genome of strain HKCCD4315 (a closed genome with one chromosome and three plasmids). The calculation of $F_{st}$ was performed with sliding windows of 10,000 bp moving in 5,000 bp steps across the core genome alignment.

*1.6 Inference of novel allelic replacement with external lineages in core genomes*

To detect genes subject to homologous recombination from external lineages, we employed a recently developed approach based on the synonymous substitution rate ($d_S$) [32]. The synonymous mutations are largely neutral since they do not cause changes in amino acid sequences. However, replacement with a divergent allele via recombination can import many synonymous variants, which leads to an anonymously large $d_S$ value between recipient genomes and the unaffected genome at this locus, compared to the remaining loci in other genomic regions. Thus, if a gene family shows that pairwise $d_S$ values between two clades are enormously large while pairwise $d_S$ values within each clade are small, it can be inferred that the allelic replacement occurred at the last common ancestor (LCA) of either clade.

Other evolutionary mechanisms may also affect the synonymous substitution rate at protein-coding genes, but they are expected to produce different $d_S$ patterns. In marine bacteria, nitrogen (N) limitation and carbon (C) limitation act as selective pressures, driving genomic G+C content to decrease and increase, respectively [33, 34]. As all genomic sites are subject to these selective pressures, synonymous sites in all genes would be affected indiscriminately. Thus, these selective pressures are not likely to affect a small proportion of gene families showing unusually large $d_S$ values. Codon usage bias imposed by translational selection is another potential force affecting the synonymous substitution rate. Different expression levels among genes lead to a varied preference for alternative synonymous codons in fast-growing microbes [35, 36]. For highly expressed genes, a stronger codon usage bias is expected to maximize the translational speed or accuracy [37], which leads to a reduced synonymous substitution rate in these genes. For genes at the regular or reduced expression level, the codon usage is a result of stochastic mutation [38, 39]. In sum, codon usage bias is not likely to give rise to outlier gene families with anonymously large $d_S$ values.

Based on these principles, we conclude that gene families with unusually large $d_S$ values are most likely subject to recombination. In practice, pairwise $d_S$ values were calculated using the YN00 program in PAML v4.9 [40] for each single-copy gene shared by the two clades. To identify core gene families showing the above-described $d_S$ pattern, the K-means clustering method was used to cluster gene families based on pairwise $d_S$ values. The number of optimal clusters was determined using R package 'NbClust' [41], which provides a variety of indices for cluster validity.

The above approach can identify which gene families were subjected to novel allelic replacements and infer the candidate ancestral branches where allelic replacements occurred. However, to determine the exact ancestral branch and the potential donor lineages, gene trees were constructed for orthologs showing unusually large between-clade $d_S$ values and compared to a phylogenomic tree. For the genome tree construction, a total of 202 published genomes are phylogenetically related to the *Ruegeria* population according to a preliminary phylogenomic tree of all publicly available *Roseobacter* genomes (data not shown) in NCBI Genbank before October, 2019. Next, a maximum phylogenomic tree was constructed using IQ-TREE v1.6.5 [24] based on the concatenation of 120 conserved genes [42] from the 202 published *Roseobacter* genomes, 20 newly sequenced genomes of the *Ruegeria* population and one randomly chosen genome from the genetically uniform *Rhodobacteraceae* population consisting of 214 strains. The phylogenomic tree was rooted using three *Oceanicella* genomes and one *Monaibacterium* genome based on their phylogenetic position recorded in the Genome Taxonomy Database (GTDB-release95) [42, 43]. For the gene tree construction, the putative orthologs were identified from the above 202 genomes using the BLASTP v2.6.0 [44] program with an *E*-value of 1e-5, and the best hit from each genome was kept. Next, MAFFT v7.215 [21] was used to align the protein sequences, and TrimAl v1.4.rev15 [22] with parameters "-automated1 -resoverlap 0.55 -seqoverlap 60" was used to trim poorly aligned sites. The maximum likelihood gene trees were subsequently constructed for each gene family based on the trimmed alignments using IQ-TREE v1.6.5 [24] with ModelFinder [45] assigning the best substitution model and with 1,000 ultrafast bootstrap replicates. To root the gene trees, outgroup lineages were chosen according to the phylogenetic placement of the 223 genomes (202 published *Roseobacter* group members, 20 strains of the *Ruegeria* population and one strain from the *Rhodobacteraceae* population) in the above phylogenomic tree. After the genome tree and the gene trees were constructed, the recombination history was manually checked by comparing gene trees to the phylogenomic tree.

*1.7 Differentiation in the accessory genome and inference of evolutionary history*

The gene presence/absence matrix of the accessory gene families was summarized as input. The Jaccard index is a measure of similarity between sample sets [46] and thus can be applied to assess gene content similarity between two strains. It is defined as the size of the intersection divided by the size of the union of the gene content in two strains, and is defined as the size of the intersection divided by the size of the union of the gene content in two strains:

$$J(S_1, S_2) = \frac{|S_1 \cap S_2|}{|S_1| + |S_2| - |S_1 \cap S_2|}$$

where $S_1$ and $S_2$ denote two strains. Thus, the Jaccard index between pairwise strains was calculated to represent gene content similarity and visualized using R.

The presence/absence of genes alone cannot reveal the evolutionary history of accessory genomes, which may provide insights into how different coral compartments drive the evolution of the two clades (clade-M and clade-S). Among the accessory genomes, the population differentiation may be largely driven by the clade-specific gene families, which are defined here as genes that are present in at least two-thirds of the strains in one clade but are present in no more than one-third of the strains in the other clade. The clade-specific genes could result from either gene gain in one clade or gene loss in the other clade, depending on the presence/absence of the gene family in the last common ancestor (LCA) shared by the two clades. As clade-M and clade-S have a closely related lineage which serves as outgroup (eight genomes), inference of the ancestral state of the LCA of the two clades could be assisted by the analysis of the phyletic pattern of the gene family in the outgroup lineage. For example, a clade-M specific gene family might be acquired at the LCA of clade-M or lost at that of clade-S. If this gene family is prevalent among the outgroup members, it was likely present at the LCA shared by clade-M and clade-S, and a reasonable inference is that this family was lost at the LCA of clade-S. Besides the ancestral branch leading to the LCA of the two clades, the evolutionary gain or loss events could also occur at the branches after the LCA. Apparently, we are more interested in the events occurring at the branch leading to the LCA of clade-M or that of clade-S, as these events may have been driving the speciation and ecological differentiation between the two clades.

In practice, the gene gain and loss history for clade-specific gene families was inferred using BadiRate v1.35 [47]. The inference was based on the parsimony rule, and the number of gene copies in each clade-specific gene family was summarized as the inputs with the parameters "--ep CSP -rmodel BDI -bmodel FR". Then the gene gain and loss history were inferred based on the predicted copy number at the ancestral node of each clade.

The genomic islands (GIs) often contain many horizontally transferred genes [48, 49]. GIs were identified for each member of the *Ruegeria* population using IslandViewer 4 [50], with the chromosome of the most closely related strain *Ruegeria* sp. AD91A (Accession number: GCA_003443535.1) was chosen as a reference genome according to the genome tree shown in Fig. S4. The clade-specific genes, the outlier genes showing unusually large $d_S$ values, and the GIs were mapped to the pangenome of the *Ruegeria* population (Fig. 2), which was visualized with Circos v0.64 [51].

*1.8 Identification of pseudogenes in the fla1 flagellar gene cluster*

The pseudogenes were identified using the program suite Psi-Phi [52] and following a modified procedure described in a recent study [53]. We used the 20 genomes of the *Ruegeria* population and another 22 closely related *Ruegeria* genomes sampled from other niches in Hong Kong coastal ecosystems (Table S6) as the pool of protein for pseudogene identification. Using the Psi-Phi program, the annotated proteins of each genome were searched against the complete nucleotide sequence of every other genome using TBLASTN [54]. The pseudogenes were recognized based on the reduced length of the protein (shorter than 80% of protein query), low BLAST *E*-values (< 1e-15), and the occurrence of premature stop codons derived from disruptive mutation.

*1.9 The physiological assays*

To compare the capability of clade-M and clade-S strains in utilizing relevant substrates, three strains from each clade were chosen to grow on a defined minimal medium with added substrates as a sole source of N and carbon C [55]. The minimum medium was modified from a carbon-free marine ammonium mineral salts (MAMS) [56] by removing the original N source ($NH_4Cl$), and thus contained the following compounds (per liter): 20 g of NaCl, 1 g of $MgSO_4 \cdot 7H_2O$, 0.2 g of $CaCl_2 \cdot 2H_2O$, 2 mg of $FeSO_4 \cdot 7H_2O$, 20 mg of $Na_2MoO_4 \cdot 2H_2O$, 0.36 g of $KH_2PO_4$, 2.34 g of $K_2HPO_4$, 1 mL of SL-10 trace metals solution [55] and 1 mL of vitamins [7].

The following organic substrates (5 mM) were added as a sole source of N and C: choline (Fig. 4A-2), glycine betaine (GBT, Fig. 4A-3), dimethylglycine (DMG, Fig. 4A-4), sarcosine (Fig. 4A-5), trimethylamine (TMA, Fig. 4A-6), trimethylamine-N-oxide (TMAO, Fig. 4A-7), creatine (Fig. 4A-8), L-proline (Fig. 4A-9), taurine (Fig. 4A-10) and urea (Fig. 4A-11). The minimum medium without any added N and C source was used as a negative control (Fig. 4A-1). The positive control was set up with a rich medium to evaluate if the optimal growth was consistent among the tested strains (Fig. 4A-1). The rich medium contained 5 g peptone and 1 g yeast extract per liter as mixed C and N sources.

If a substrate can barely support the tested strains as a sole source of N and C, it was further examined as a sole N source and a sole C source separately. Besides, dimethylsulfoniopropionate (DMSP) was also tested as the sole C source. For the sole C source assay, 10 mM $NH_4Cl$ was added as the N source (Fig. 4B-2, 4B-3, 4B-5, 4B-7, 4B-9 and 4B-11). For the sole N source assay, 5 mM sodium pyruvate was added as the C source (Fig. 4B-4, 4B-6, 4B-8, 4B-10 and 4B-12). To control the potential growth bias introduced by the $NH_4Cl$ or pyruvate, positive control was set up with 10 mM $NH_4Cl$ as a sole N source and 5 mM sodium pyruvate as a sole C source (Fig. 4B-1), and negative control was set up with no added C and N source (Fig. 4B-1).

Three replicates for each treatment were conducted in 50 mL tubes at 28 °C, and growth was examined by measuring the optical density at 600 nm ($OD_{600}$). The growth rate was calculated using the data points in the exponential phase. The relative growth yield was represented by the $OD_{600}$ of the final points in the exponential phase of the growth curve. Both the growth rate and growth yield were compared between members from clade-M and those from clade-S. The differences between clades were statistically evaluated with One-way Repeated Measures ANOVA, with $p < 0.05$ indicating that the growth rates and the relative growth yield between clades are significantly different [57].

The swimming, swarming, and twitching motility of the clade-M and clade-S members were tested on agar plates with 2216E marine broth medium [BD Difco, USA]. Overnight cultures of each strain were inoculated in 2216E marine broth medium with 1:200 dilution and incubated at 28°C under a shaking speed of 200 rpm until the $OD_{600}$ of the culture reached 0.6-0.8. For the swimming test, 0.3% (w/v) soft agar plates were point-inoculated with 3 μL of the fresh cell suspension and incubated at 28°C for eight days. The zone of swimming motility on the surface of agar was measured. The swarming assay was conducted following the same procedure but with the concentration of agar plates replaced by 0.6% (w/v). For the twitching assay, cell cultures were stab-inoculated to the bottom of the Petri dish of 1.0 % (w/v) agar plate with a sterile toothpick and incubated in the humidified box at 28 ℃ for 10 days, and the zone of twitching motility at the interface of agar and Petri dish was measured. All tests were done in triplicates.

*1.10 Test of compartmentalization and dispersal limitation*

The genetically uniform *Rhodobacteraceae* population of 214 strains varying at only a few dozen nucleotide sites across the whole genomes were partitioned into four subpopulations, members of each exclusively isolated from distinct coral individuals. These four coral individuals were collected from four different sampling locations (Fig. S1A). The two subpopulations from Wong Wan Chau (WWC) and Ngo Mei Chau (NMC) each contain isolates cultured from multiple compartments (Table S6), and thus they are amenable for calculating the number of migrations between compartments.

The tree-based Slatkin-Maddison test [58] implemented in HyPhy v2.5 [59] was employed to evaluate the compartmentalization of the two subpopulations. For each subpopulation, the maximum likelihood phylogenetic tree was constructed using IQ-TREE v1.6.5 [24] based on the core SNPs identified by kSNP v3.0, a software quickly and accurately identifying SNPs among hundreds of genomes in an alignment-free approach [60]. Next, strains were labeled according to their compartment of isolation (i.e., mucus, tissue and skeleton). The strains isolated from ambient seawater of coral were not considered. Since the identical siblings from the same compartment could amplify the signal of compartmentalization [61], only one of the siblings with branch length zero from the same compartment was kept and the remaining ones were pruned from the phylogenetic tree. To calculate the number of migrations observed in each subpopulation, the pruned phylogenetic tree with compartment information was subject to standard Slatkin-Maddison test, followed by 100,000 permutations of population structures to generate a normal distribution representing the number of migrations expected by chance. Whether isolates were significantly compartmentalized was determined by comparing the observed and expected number of migrations

*1.11 Estimating the origin time for the Rhodobacteraceae and the Ruegeria populations*

To estimate the origin time of the two *Rhodobacteraceae* subpopulations (WWC and NMC), we followed the formula:

$$S = \mu * G * L * T$$

where $S$ is the number of point mutations that have occurred in a population, $\mu$ is the base-substitution mutation rate (per nucleotide site per cell division), $G$ is the growth rate in the field (number of cell divisions per year), $L$ is the number of nucleotide sites (i.e., average genome size), $T$ is the evolutionary time (years). Because both WWC and NMC subpopulations showed genetic monomorphism which varies only at a few dozen SNP sites, the number of point mutations can be approximated by the number of SNP sites (27 SNPs in WWC subpopulation and 21 SNPs in NMC subpopulation). Since the mutation rate is not available for these subpopulations, we turned to the model roseobacter *Ruegeria pomeroyi* DSS-3 [62], whose unbiased spontaneous mutation rate ($1.39 \times 10^{-10}$ per site per generation) was determined using the mutation accumulation experiment followed by whole genome sequencing of the mutant lines [63]. Likewise, the growth rate is also not known for these subpopulations, so we used the published data (averaging to one cell division per day) previously determined for pelagic roseobacters in several coastal waters [64].

In the case of the *Ruegeria* population, which showed much greater diversity, purifying selection at the protein sequence level may have acted to purge diversity at the nonsynonymous (amino acid altering) nucleotide sites, leaving 342,831 SNPs at the synonymous (silent) nucleotide sites useful for time estimation. In addition, a few core genes were subjected to allelic replacement by homologous recombination with externally divergent species (Supplemental Text 1.6). As these genes showed unusually large synonymous substitution rates, they cannot be used for time estimation. This led to the exclusion of 25,211 synonymous SNPs occurring in these core genes. Next, 14,124 triallelic and 251 tetrallelic synonymous SNPs were identified, and for a conservative estimate each of these SNPs was assumed to be introduced through a single event of point mutation. Further, homologous recombination may also have occurred between members within the population, which is best characterized by identifying homoplasious bi-allelic SNPs [65], though a small proportion of homoplasious bi-allelic SNPs can be caused by convergent mutations [66, 67]. Similarly, 34,567 homoplasious bi-allelic SNPs each were treated as a single point mutation. The remaining biallelic 267,678 synonymous SNPs were either autapomorphic or synapomorphic, each best explained by a single point mutation event. Following this rationale, we estimated a total of 316,620 mutations at synonymous sites that have occurred in the *Ruegeria* population, with the caveat that treating the triallelic, tetrallelic, and homoplasious bi-allelic SNPs as single mutation events may underestimate the true number of point mutation events. Next, the timescale was estimated following the procedure detailed for the *Rhodobacteraceae* WWC and NMC subpopulations, with the $L$ replaced by $L_{syn}$ (the number of synonymous sites in all core genes of the *Ruegeria* population, $L_{syn}$ = 829,505).

**Text 2. Supplementary results:**

*2.1 Population differentiation at the core genomes of the Ruegeria population*

Speciation accompanies a decreased recombination frequency between differentiated populations. The $\rho/\theta$ ratio measures the relative rate of recombination to point mutation, and a threshold of 0.25-0.5 delineates the clonality of a bacterial population [68]. The ClonalFrameML v1.11 [27] analysis showed that this ratio between the two clades was only 0.05, whereas those within clade-M and within clade-S were 0.51 and 0.34, respectively (Table S1). Likewise, the r/m ratio assesses the relative effect of recombination to point mutation on genetic variation, and the result of ClonalFrameML v1.11 showed this ratio between the two clades (0.67) to be much lower than that within each clade (3.13 for clade-M and 4.63 for clade-S, Table S1). The decreased $\rho/\theta$ ratio and r/m ratio between the two clades compared to those within each clade indicates that there is a strong barrier to gene flow between clade-M and clade-S.

Without a strong cohesive force by homologous recombination, the genetic differentiation between the two clades is expected, which may have led to the fixation of different alleles. In total, we identified 502,661 single nucleotide polymorphisms (SNPs) from 3. 47 Mbp core genomes shared by all members of clade-M and clade-S (Table 1). Among them, 302,836 (60.2%) were fixed differences between the two clades. Next, we measured the level of differentiation by calculating $F_{st}$ values along the core genome in a sliding window of 10 kbp with 5 kbp steps (Fig. S2). Most of the genomic regions (96.8%) showed a high level of differentiation ($F_{st} \geq 0.5$), whereas only several patchy regions showed a lower level of differentiation ($F_{st} < 0.5$) (Fig. S2). The permutation test showed that 924 out of the 934 genomic regions were significantly differentiated ($p < 0.05$), suggesting that speciation between the two clades may have already reached completion.

*2.2 The Ruegeria population differentiation at the physiological level*

As discussed above and in the main paper, the clade-specific accessory genes and core genes that show unusually large $d_S$ values are involved in the utilization of several ecologically relevant substrates, including choline, GBT, sarcosine, TMA, TMAO, creatine, L-proline, taurine and urea. The utilization of these substrates each as a sole source of both C and N by these two clades were tested and compared. As a control, all strains did not grow without C and N sources (open circles in Fig. 4A-1), and grew equally well under optimal conditions with a replete supply of C and N (open triangles in Fig. 4A-1). This indicates that any differences of growth traits in the following assays supplemented with a specific substrate as a sole source of both C and N can be ascribed to the distinct responses of the bacteria to the added substrate. Here, we provided details of the assay results.

First, members of the two clades showed distinct responses to the addition of methylamine-related coral osmolytes including choline, GBT, DMG, sarcosine, TMA, TMAO and creatine. All six strains grew poorly when choline (Fig. 4A-2), GBT (Fig. 4A-3), TMA (Fig. 4A-6) and TMAO (Fig. 4A-7) each were used as a sole C and N source. However, for the intermediates laying downstream of GBT (Fig. 3), such as DMG (Fig. 4A-4) and sarcosine (Fig. 4A-5), the assayed bacteria generally grew and the clade-M members showed significantly higher growth rates and growth yields ($p < 0.05$, One-way Repeated-Measures ANOVA; the same test used below unless stated otherwise) on sarcosine compared to the clade-S members.

Next, we assayed the bacterial growth on other coral osmolytes including DMSP, L-proline and taurine, as well as urea mainly from the excretions of animals in coral reef ecosystems. When serving as a sole C source, DMSP supported a higher growth yield for the clade-M members than the clade-S members ($p < 0.05$; Fig. 4B-2). When growing on L-proline as a sole C and N source, the overall growth rates and growth yields of the clade-M members were significantly higher than those of the clade-S members ($p < 0.05$; Fig. 4A-9). When taurine was supplied as a sole C and N source, the clade-M members showed significantly higher growth yields than the clade-S members ($p < 0.05$; Fig. 4A-10), but no significant growth rate difference was observed. These results support the bioinformatics predictions on the additional copies specific to clade-M (i.e., *dddP*, *dmdABCD*, *tauABC* and *proVWX*) and shared genes subjected to novel allelic replacements (i.e., *dddD* and *tauABC*). When growing on urea, all tested strains grew weakly and showed no significant difference in both the growth rates and yields (Fig. 4A-11).

The physiological assays also showed that choline (Fig. 4B-2), GBT (Fig. 4B-3), TMA (Fig. 4B-6), TMAO (Fig. 4B-7) and urea (Fig. 4B-11) cannot act as a sole C and N source. We therefore further tested if they may serve as a sole C source or a sole N source separately. In the control group, all strains did not grow without C and N sources (open circles in Fig. 4B-1) or grew equally well when pyruvate and ammonium were used as C and N sources (open triangles in Fig. 4B-1), respectively. These control experiments indicate that the addition of these common C and N sources did not contribute to growth differences. In other words, when pyruvate or ammonium was replaced by the tested substrates in the experimental group, the growth differences, if any, can be ascribed to the differential responses of the bacteria to the tested substrate. When choline (Fig. 4B-3), GBT (Fig. 4B-5), TMA (Fig. 4B-7), TMAO (Fig. 4B-9) and urea (Fig. 4B-11) each was used as a sole C source, all bacteria grew poorly and did not show significant between-clade differences. However, choline (Fig. 4B-4), GBT (Fig. 4B-6), TMA (Fig. 4B-8) and TMAO (Fig. 4B-10) can serve as a sole N source to support both clade-M and clade-S strains. Using choline or GBT as a sole N source respectively, the clade-M members showed significantly higher growth yields than the clade-S members ($p < 0.05$), though the growth rates between clades showed no significant difference. Moreover, there was no significant difference in growth rates and yields between clade-M and clade-S when using TMA or TMAO as a sole N source.

### 2.3 Metabolic potential for utilizing other substrates by the mucus clade of the Ruegeria population

Carbohydrates also account for a large proportion of the osmolytes in corals [69]. L-fucose is an osmolyte present in coral secretions and also part of oligosaccharides, mucins, and other glycoconjugates in the surface mucus layer [2, 70]. A gene cluster involved in fucose catabolism was found specific to the clade-M (HKCCD4315_03759-03763, Table S3), including L-fucose mutarotase (*fucU*), L-fuconolactone hydrolase, L-fuconate dehydrogenase, ketoglutarate semialdehyde dehydrogenase and 2-keto-3-deoxy-L-fuconate dehydrogenase. This cluster was located on a plasmid, with three genes of this cluster inferred to be gained at the LCA of clade-M, and two lost at the LCA of clade-S (Table S3). Through this pathway, fucose is degraded to pyruvate and L-lactate via non-phosphorylated intermediates [71, 72]. Besides, fucose in host mucus acts as important attractants for symbiotic microbiota, and the utilization of fucose might also provide microbes with a competitive advantage in their niche colonization [72-74]. Families are related to the utilization of other unknown monosaccharides (Table S3), and these genes are located on a plasmid and were inferred to be gained at the LCA of clade-M, suggesting that the carbohydrates in mucus are another important factor driving the diversification of clade-M from clade-S.

Some of the aromatic compounds, such as polycyclic aromatic hydrocarbons (PAHs), are ubiquitous pollutants in coral reefs, and are concentrated in coral mucus due to their high lipid-solubility [75]. Hong Kong is one of the busiest seaports in the world, so oil spills occur frequently in Hong Kong coastal waters [76, 77]. The coral mucus in this region is known to contain aromatic pollutants such as PAHs [75]. Members of the *Rhodobacteraceae* play a major role in degrading aromatics including PAHs in natural communities [78]. A clade-M specific gene encoding the ring-cleaving enzyme which potentially acts on PAHs was likely acquired at the LCA of clade-M (HKCCD4315_02148, Table S3). Besides, genes (*pcaBDHG*; Table S3) in the protocatechuate pathway responsible for further degradation of the aromatic intermediates were identified exclusively in the clade-M members. These results suggest that the clade-M strains may be able to degrade aromatic pollutants like PAH and benefit the coral hosts. However, the majority of aromatics degrading genes were likely acquired after the branching of clade-M (Table S3), suggesting that these genes were a later innovation facilitating the mucus niche adaptation.

#### *2.4 Metabolic potential of the mucus clade in the Ruegeria population underlying microbial interactions in the densely-populated mucus habitat*

The coral mucus is a eutrophic niche enriched with native flora, and the associated microbial community structure is shaped by microbial interactions [79]. Bacterial quorum sensing (QS) is a widespread signaling mechanism acting at a high cell density [80]. Its canonical signaling molecules, N-acyl-homoserine lactones (AHLs), have been detected in coral mucus [81]. Bacteria respond to the AHLs signals and activate the QS circuits through the two-component system, LuxIR, and degrade the signal molecular through N-acyl homoserine lactonase [80]. The additional gene copies for the LuxIR system and N-acyl homoserine lactonase were found located in the clade-M strains specific genome region (Table S3).

In the *Rhodobacteraceae*, biofilm formation is an effective strategy to compete with other organisms for space and nutrients, which could be a response to QS signals [82]. Extracellular polymeric substances (EPS) are important components of the biofilm matrix [83]. We found that clade-M members possess several clade-specific genes involved in the synthesis of EPS such as exopolysaccharides (HKCCD4315_04016, HKCCD4315_04166, and HKCCD4315_04162, Table S3), and lipopolysaccharides (HKCCD4315_ 04159 and HKCCD4315_04161, Table S3). These compounds are the structural components of the biofilm matrix [82]. Most of the aforementioned genes (five out of eight) involved in the QS system and biofilm formation were inferred to be acquired at the LCA of the clade-M, which may have facilitated the adaptation of clade-M to the mucus niche of high cell density.

#### *2.5 Adaptation of the skeleton clade in the Ruegeria population to the periodically anoxic skeleton habitat*

The skeleton is a diurnally anoxic environment [84]. The oxygen produced by the photosynthesis of symbiotic and endolithic algae diffuses through the porous aragonite into the skeleton core in the daytime, and is continuously consumed by the coral host and associated community through respiration [84]. Thus, the skeleton undergoes a sharp decrease of dissolved oxygen at night, and resulting in anoxic in the skeleton [85]. A gene cluster associated with anaerobic respiration was identified exclusively in the clade-S members. This cluster encoded a succinate dehydrogenase/fumarate reductase (*sdh-frd*, Table S4) and L(+)-tartrate dehydratase (*ttdAB*, Table S4). The Sdh-Frd is bifunctional in some facultative anaerobes. It catalyzes succinate oxidation to support the citric acid cycle under oxic conditions. Meanwhile, it does the reverse reduction under anoxic condition, and employs fumarate as an electron acceptor to maintain the anoxic respiration in the absence of oxygen [86]. The Ttd enzyme is responsible for the fermentation of L-tartrate, which supports the anaerobic growth of bacteria as C and energy source [87, 88]. As this gene cluster is located on a plasmid that is not linked to the chromosomal genes encoding the canonical citric acid cycle-related enzymes, it is more likely related to anaerobic respiration rather than the citric acid cycle. It was inferred to be acquired at the LCA of clade-S, indicating an adaptation of clade-S strains in periodically anoxic skeleton niches. Besides, a gene cluster encoding the dimethyl sulfoxide/trimethylamine oxide reductase (*dmsABC*, Table S5) was identified as core genes with unusually large $d_S$ value. This gene is involved in anaerobic respiration and enables the bacteria to use either dimethyl sulfoxide (DMSO) or trimethylamine-N-oxide (TMAO) as a terminal electron acceptor anaerobically for oxidative phosphorylation [89]. The gene trees showed that the three gene families subject to distinct evolutionary history (Table S5). Together, the clade-S strains likely gained fitness advantages in skeleton niches through enhanced anoxic tolerance.

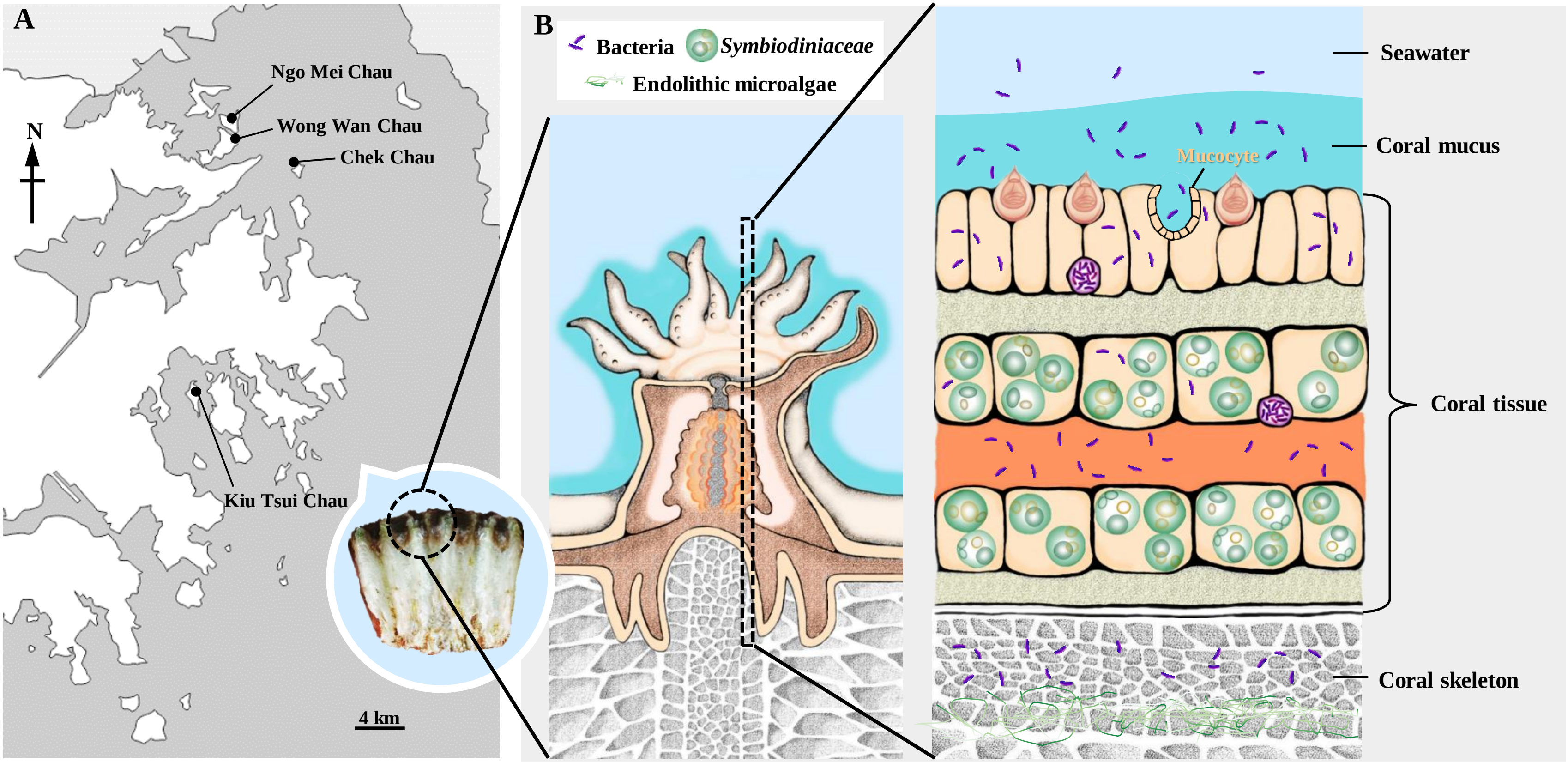


**Figure S1**. Sampling information of coral *Platygyra acuta*. **(A)** The sampling sites in Hong Kong seawater: Wong Wan Chau (N 22° 31‘31.2“ E 114° 19’00.1”), Ngo Mei Chau (N 22° 31‘47.2“ E 114° 19’02.9”), Chek Chau (N 22° 30‘03.3“ E 114° 21’22.7”) and Kiu Tsui Chau (N 22° 22‘04.4“ E 114° 17’42.0”). **(B)** A cartoon shows the compartments of coral. The associated bacteria, endosymbiont *Symbiodiniaceae* and endolithic microalgae are showed in different compartments. Part of the cartoon is adapted from Bourne et al., 2016.

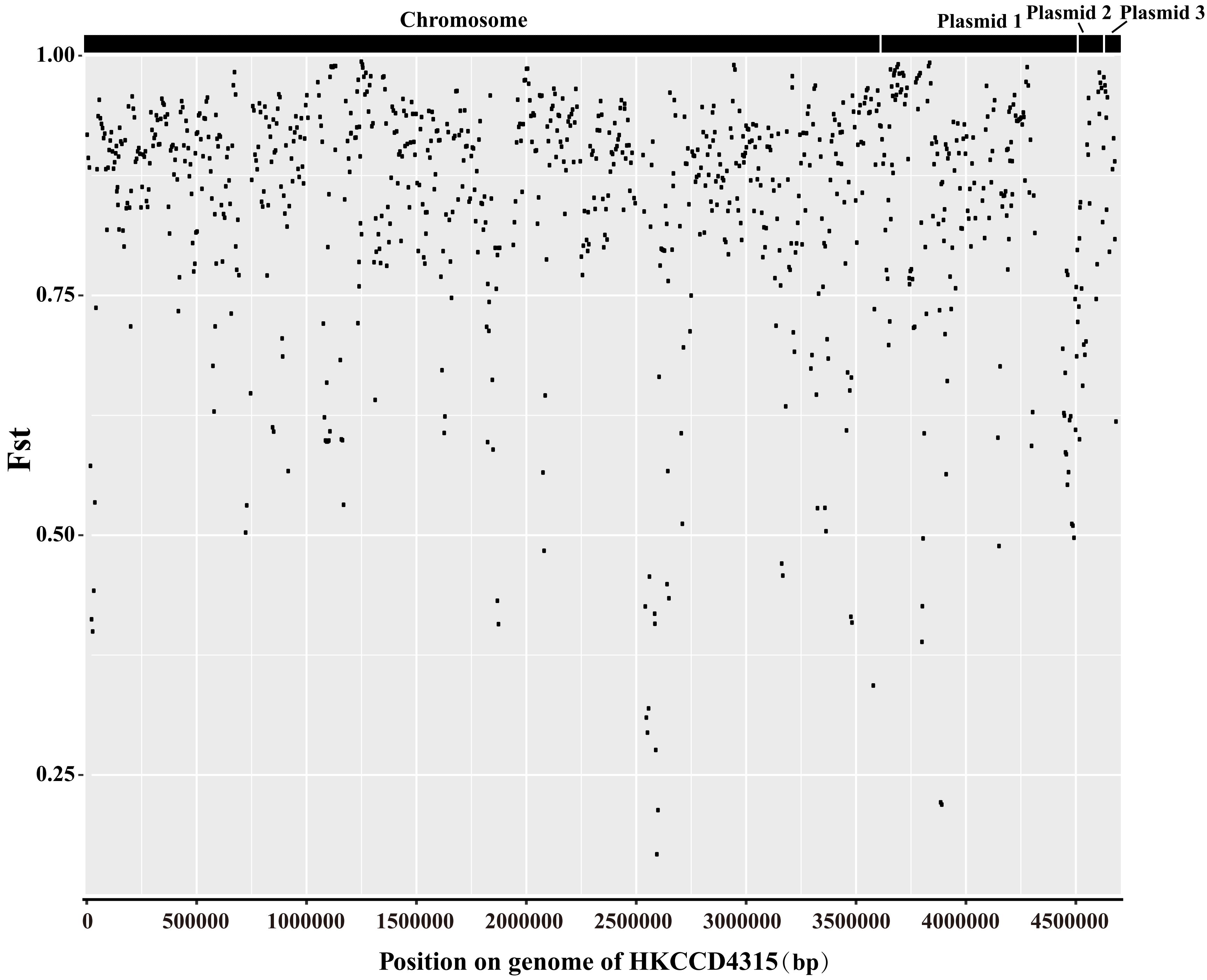


**Figure S2.** Fst values between the clade-M and clade-S strains along the closed genome of strain HKCCD4315. The upper panel showed the genomic region of the chromosome and three plasmids of strain HKCCD4315. Each dot represents an Fst value calculated for a sliding window of 10 kb moving in 5 kb steps.

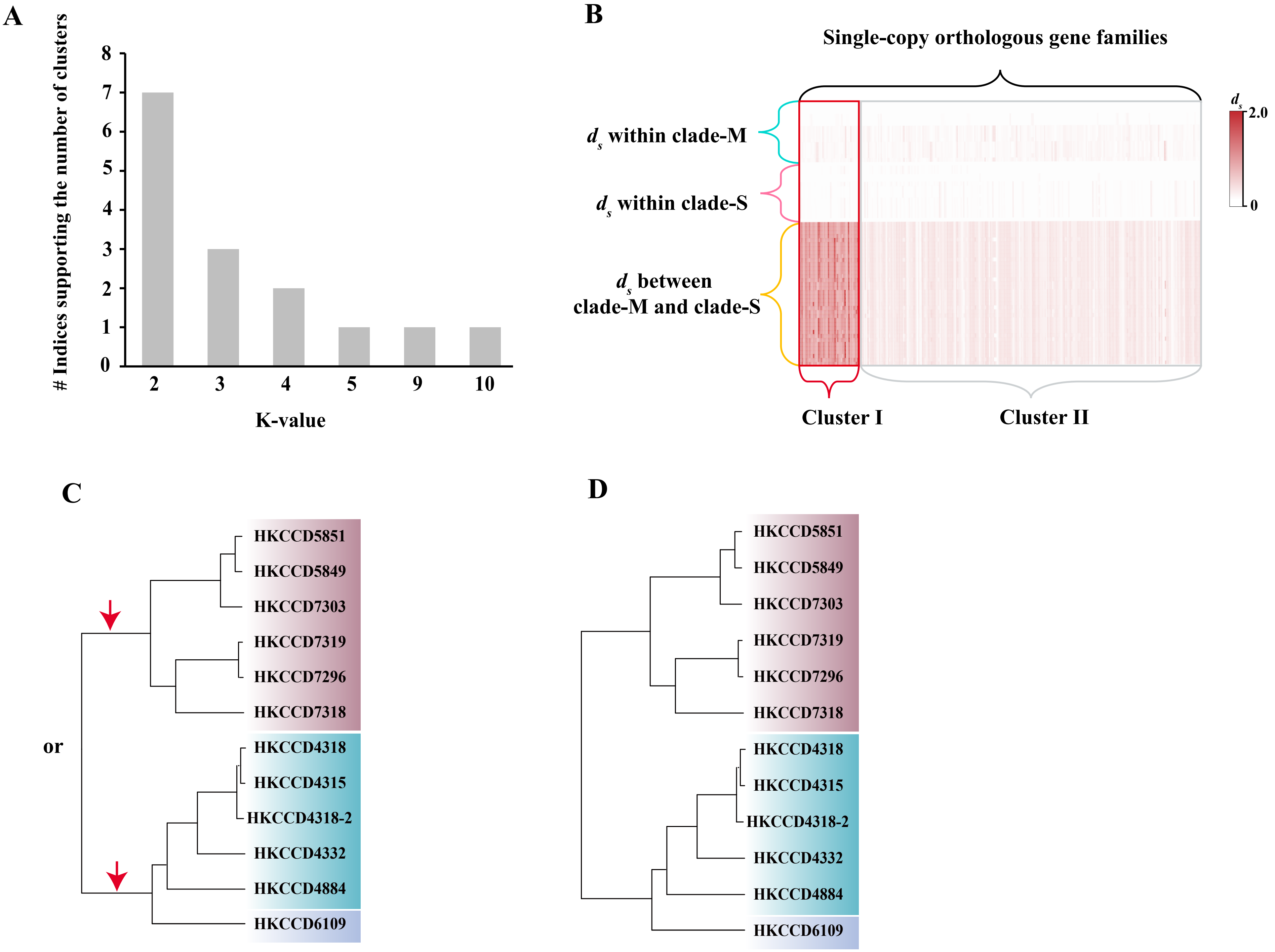


**Figure S3.** Illustration of allelic replacement inference using *ds* values. **(A)** The voting of the optimal number of cluster validity indices using R package "NbCluster". The seven indices supporting K=2 as the optimal clustering number include 'beale', 'duda', 'mcclain', 'pseudot2', 'ptbiserial', 'gap' and 'sdindex'. **(B)** The demo heatmap of the *ds* values were calculated for every possible pair of genomes across all single-copy orthologous gene families, with warmer colour indicating higher *ds* values. The gene families were grouped into two clusters using the K-means method. Cluster I shows enormously large *ds* values between clade-M and clade-S but small *ds* values within each clade, whereas all *ds* values are small in Cluster II. **(C)** The evolutionary history of an example gene from Cluster I was mapped to the genome tree. Due to the unusually large between-clade *ds* values and little diversity within each clade, the allelic replacement with distant lineages was inferred to have occurred at the LCA of clade-M or that of clade-S. **(D)** The evolutionary history of an example gene from Cluster II was mapped to the genome tree. Because of the small diversity both between- and within-clade, allelic replacement with distant lineages is less likely to occur.

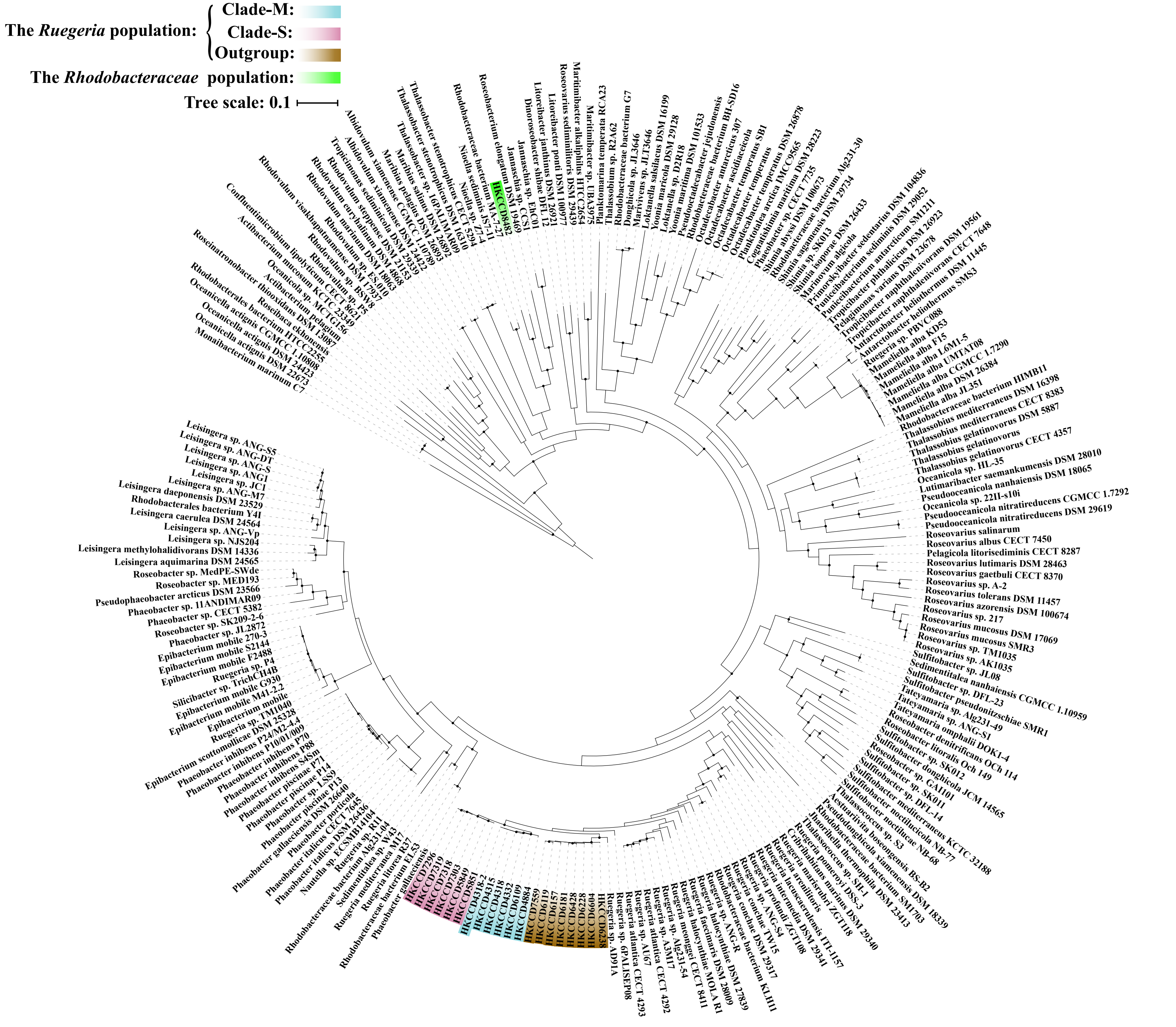


**Figure S4.** The maximum likelihood phylogeny of the *Roseobacter* group, which was constructed using 223 publicly available *Roseobacter* genomes, the 20 strains of the *Ruegeria* population and one representative of the genetically uniform *Rhodobacteraceae* population. Solid circles at the nodes indicate that the ultrafast bootstrap support value of the branch is ≥ 95%. The scale bar indicates number of substitutions per site. The members of clade-M, clade-S and outgroup clade were shadowed with blue, pink and gold, respectively.

**Figure S5.** The maximum likelihood phylogeny of the core gene families *ugpA* , *ugpB*, *ugpE*, *betA* and dmgdh involved in methylamine utilization constructed by IQ-TREE v1.6.5. Solid circles at the nodes indicate that the ultrafast bootstrap support value of the branch is ≥ 80%. The scale bar indicates number of substitutions per site. The members of clade-M, clade-S and outgroup clade were shadowed with blue, pink and gold, respectively.

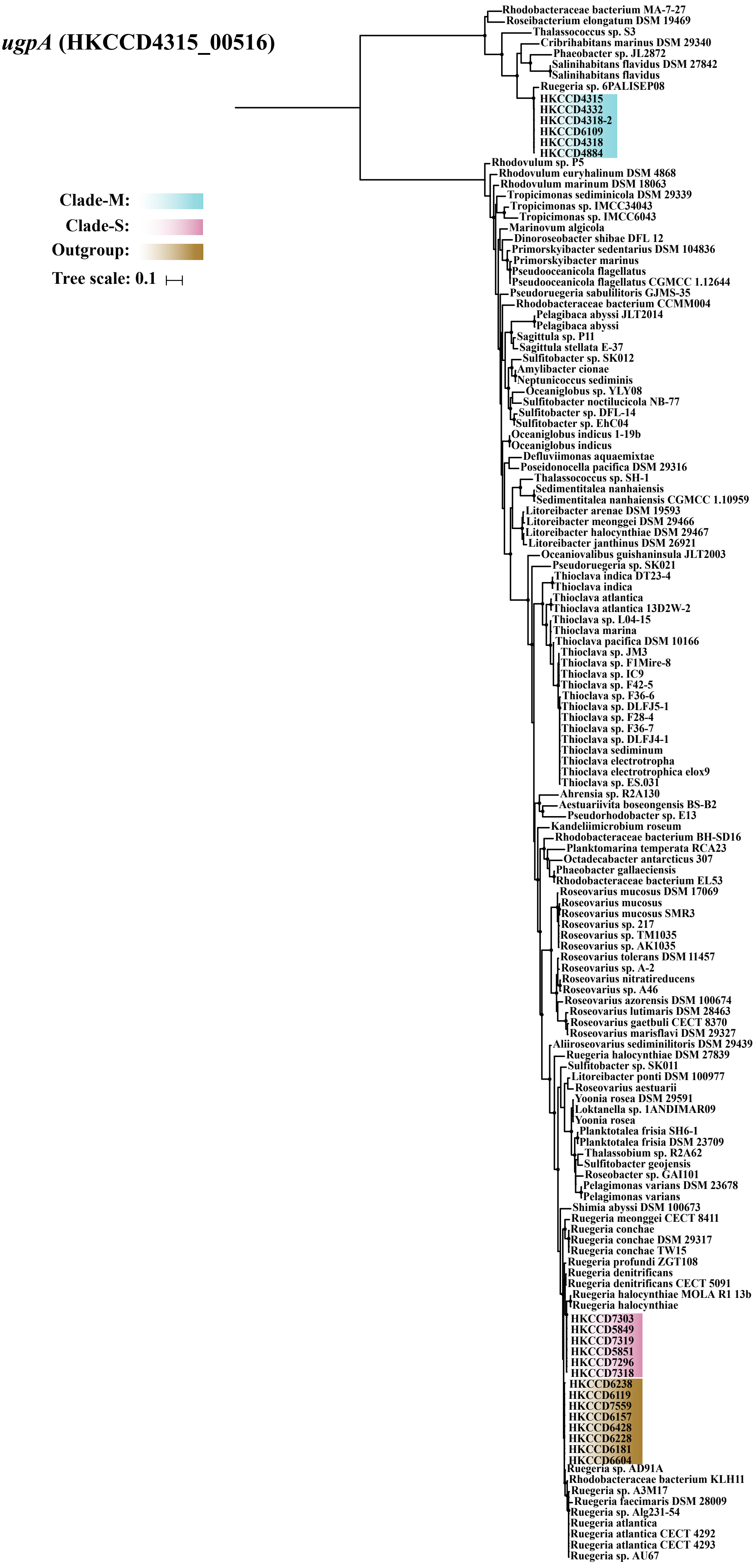
ugpA (HKCCD4315_00516)
Clade-M:
Clade-S:
Outgroup:
Tree scale: 0.1
Rhodobacteraceae bacterium MA-7-27
Roseibacterium elongatum DSM 19469
Thalassococcus sp. S3
Cribrihabitans marinus DSM 29340
Phaeobacter sp. JL2872
Salinihabitans flavidus DSM 27842
Salinihabitans flavidus
Ruegeria sp. 6PALISEP08
HKCCD4315
HKCCD4332
HKCCD4318-2
HKCCD6109
HKCCD4318
HKCCD4884
Rhodovulum sp. P5
Rhodovulum euryhalinum DSM 4868
Rhodovulum marinum DSM 18063
Tropicimonas sediminicola DSM 29339
Tropicimonas sp. IMCC34043
Tropicimonas sp. IMCC6043
Marinovum algicola
Dinoroseobacter shibae DFL 12
Primorskyibacter sedentarius DSM 104836
Primorskyibacter marinus
Pseudooceanicola flagellatus
Pseudooceanicola flagellatus CGMCC 1.12644
Pseudoruegeria sabulilitoris GJMS-35
Rhodobacteraceae bacterium CCMM004
Pelagibaca abyssi JLT2014
Pelagibaca abyssi
Sagittula sp. P11
Sagittula stellata E-37
Sulfitobacter sp. SK012
Amylibacter cionae
Neptunicoccus sediminis
Oceaniglobus sp. YLY08
Sulfitobacter noctilucicola NB-77
Sulfitobacter sp. DFL-14
Sulfitobacter sp. EhC04
Oceaniglobus indicus 1-19b
Oceaniglobus indicus
Defluviimonas aquaemixtae
Poseidonocella pacifica DSM 29316
Thalassococcus sp. SH-1
Sedimentitalea nanhaiensis
Sedimentitalea nanhaiensis CGMCC 1.10959
Litoreibacter arenae DSM 19593
Litoreibacter meonggei DSM 29466
Litoreibacter halocynthiae DSM 29467
Litoreibacter janthinus DSM 26921
Oceaniovalibus guishaninsula JLT2003
Pseudoruegeria sp. SK021
Thioclava indica DT23-4
Thioclava indica
Thioclava atlantica
Thioclava atlantica 13D2W-2
Thioclava sp. L04-15
Thioclava marina
Thioclava pacifica DSM 10166
Thioclava sp. JM3
Thioclava sp. F1Mire-8
Thioclava sp. IC9
Thioclava sp. F42-5
Thioclava sp. F36-6
Thioclava sp. DLFJ5-1
Thioclava sp. F28-4
Thioclava sp. F36-7
Thioclava sp. DLFJ4-1
Thioclava sediminum
Thioclava electrotropha
Thioclava electrotrophica elox9
Thioclava sp. ES.031
Ahrensia sp. R2A130
Aestuariivita boseongensis BS-B2
Pseudorhodobacter sp. E13
Kandeliimicrobium roseum
Rhodobacteraceae bacterium BH-SD16
Planktomarina temperata RCA23
Octadecabacter antarcticus 307
Phaeobacter gallaeciensis
Rhodobacteraceae bacterium EL53
Roseovarius mucosus DSM 17069
Roseovarius mucosus
Roseovarius mucosus SMR3
Roseovarius sp. 217
Roseovarius sp. TM1035
Roseovarius sp. AK1035
Roseovarius tolerans DSM 11457
Roseovarius sp. A-2
Roseovarius nitratireducens
Roseovarius sp. A46
Roseovarius azorensis DSM 100674
Roseovarius lutimaris DSM 28463
Roseovarius gaetbuli CECT 8370
Roseovarius marisflavi DSM 29327
Aliiroseovarius sediminilitoris DSM 29439
Ruegeria halocynthiae DSM 27839
Sulfitobacter sp. SK011
Litoreibacter ponti DSM 100977
Roseovarius aestuarii
Yoonia rosea DSM 29591
Loktanella sp. 1ANDIMAR09
Yoonia rosea
Planktotalea frisia SH6-1
Planktotalea frisia DSM 23709
Thalassobium sp. R2A62
Sulfitobacter geojensis
Roseobacter sp. GAI101
Pelagimonas varians DSM 23678
Pelagimonas varians
Shimia abyssi DSM 100673
Ruegeria meonggei CECT 8411
Ruegeria conchae
Ruegeria conchae DSM 29317
Ruegeria conchae TW15
Ruegeria profundi ZGT108
Ruegeria denitrificans
Ruegeria denitrificans CECT 5091
Ruegeria halocynthiae MOLA R1 13b
Ruegeria halocynthiae
HKCCD7303
HKCCD5849
HKCCD7319
HKCCD5851
HKCCD7296
HKCCD7318
HKCCD6238
HKCCD6119
HKCCD7559
HKCCD6157
HKCCD6428
HKCCD6228
HKCCD6181
HKCCD6604
Ruegeria sp. AD91A
Rhodobacteraceae bacterium KLH11
Ruegeria sp. A3M17
Ruegeria faecimaris DSM 28009
Ruegeria sp. Alg231-54
Ruegeria atlantica
Ruegeria atlantica CECT 4292
Ruegeria atlantica CECT 4293
Ruegeria sp. AU67

***ugpB*** **(HKCCD4315_00515)**

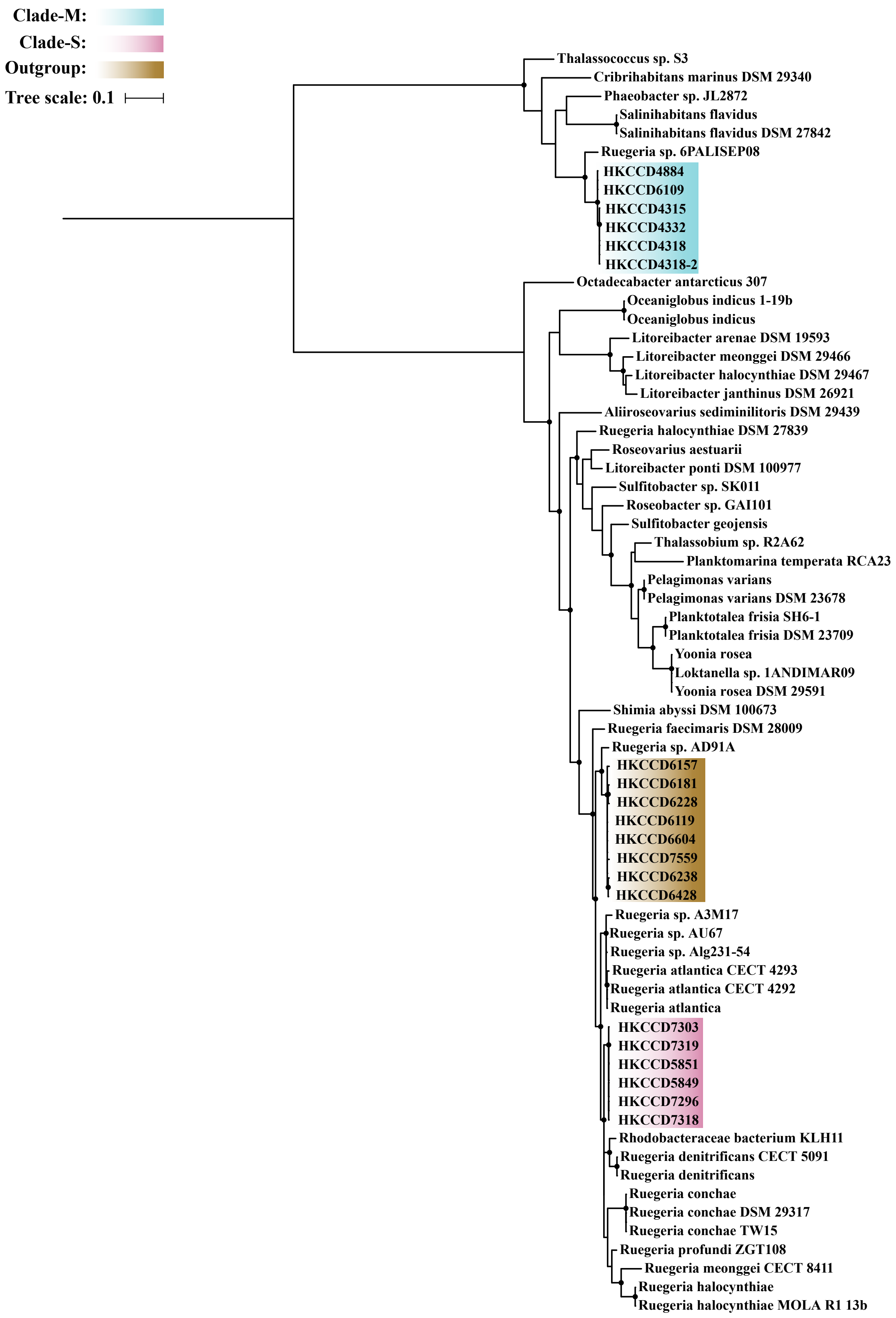

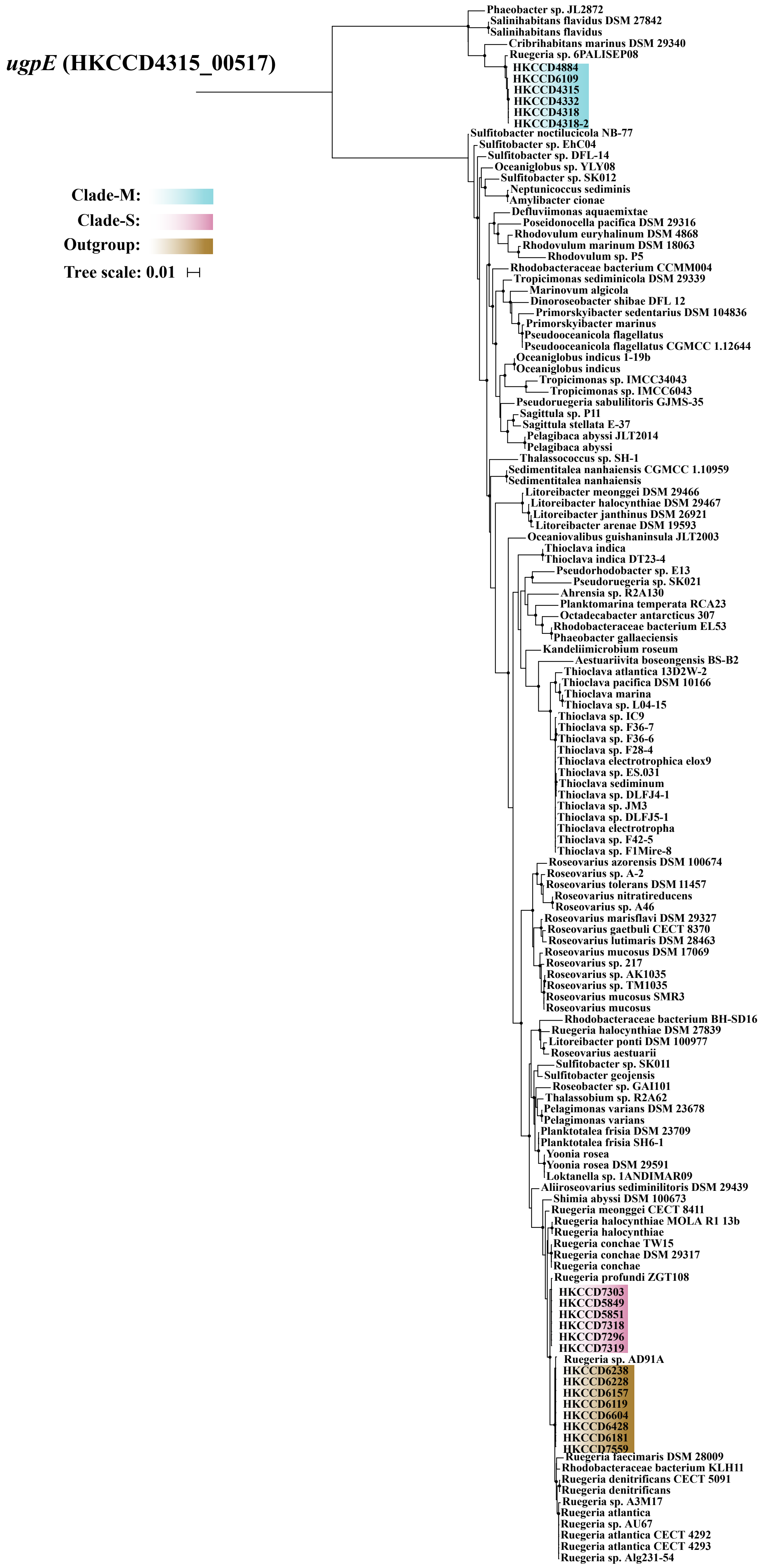
ugpE (HKCCD4315_00517)
Clade-M:
Clade-S:
Outgroup:
Tree scale: 0.01
Phaeobacter sp. JL2872
Salinihabitans flavidus DSM 27842
Salinihabitans flavidus
Cribrihabitans marinus DSM 29340
Ruegeria sp. 6PALISEP08
HKCCD4884
HKCCD6109
HKCCD4315
HKCCD4332
HKCCD4318
HKCCD4318-2
Sulfitobacter noctilucicola NB-77
Sulfitobacter sp. EhC04
Sulfitobacter sp. DFL-14
Oceaniglobus sp. YLY08
Sulfitobacter sp. SK012
Neptunicoccus sediminis
Amylibacter cionae
Defluviimonas aquaemixtae
Poseidonocella pacifica DSM 29316
Rhodovulum euryhalinum DSM 4868
Rhodovulum marinum DSM 18063
Rhodovulum sp. P5
Rhodobacteraceae bacterium CCMM004
Tropicimonas sediminicola DSM 29339
Marinovum algicola
Dinoroseobacter shibae DFL 12
Primorskyibacter sedentarius DSM 104836
Primorskyibacter marinus
Pseudooceanicola flagellatus
Pseudooceanicola flagellatus CGMCC 1.12644
Oceaniglobus indicus 1-19b
Oceaniglobus indicus
Tropicimonas sp. IMCC34043
Tropicimonas sp. IMCC6043
Pseudoruegeria sabulilitoris GJMS-35
Sagittula sp. P11
Sagittula stellata E-37
Pelagibaca abyssi JLT2014
Pelagibaca abyssi
Thalassococcus sp. SH-1
Sedimentitalea nanhaiensis CGMCC 1.10959
Sedimentitalea nanhaiensis
Litoreibacter meonggei DSM 29466
Litoreibacter halocynthiae DSM 29467
Litoreibacter janthinus DSM 26921
Litoreibacter arenae DSM 19593
Oceaniovalibus guishaninsula JLT2003
Thioclava indica
Thioclava indica DT23-4
Pseudorhodobacter sp. E13
Pseudoruegeria sp. SK021
Ahrensia sp. R2A130
Planktomarina temperata RCA23
Octadecabacter antarcticus 307
Rhodobacteraceae bacterium EL53
Phaeobacter gallaeciensis
Kandeliimicrobium roseum
Aestuariivita boseongensis BS-B2
Thioclava atlantica 13D2W-2
Thioclava pacifica DSM 10166
Thioclava marina
Thioclava sp. L04-15
Thioclava sp. IC9
Thioclava sp. F36-7
Thioclava sp. F36-6
Thioclava sp. F28-4
Thioclava electrotrophica elox9
Thioclava sp. ES.031
Thioclava sediminum
Thioclava sp. DLFJ4-1
Thioclava sp. JM3
Thioclava sp. DLFJ5-1
Thioclava electrotropha
Thioclava sp. F42-5
Thioclava sp. F1Mire-8
Roseovarius azorensis DSM 100674
Roseovarius sp. A-2
Roseovarius tolerans DSM 11457
Roseovarius nitratireducens
Roseovarius sp. A46
Roseovarius marisflavi DSM 29327
Roseovarius gaetbuli CECT 8370
Roseovarius lutimaris DSM 28463
Roseovarius mucosus DSM 17069
Roseovarius sp. 217
Roseovarius sp. AK1035
Roseovarius sp. TM1035
Roseovarius mucosus SMR3
Roseovarius mucosus
Rhodobacteraceae bacterium BH-SD16
Ruegeria halocynthiae DSM 27839
Litoreibacter ponti DSM 100977
Roseovarius aestuarii
Sulfitobacter sp. SK011
Sulfitobacter geojensis
Roseobacter sp. GAI101
Thalassobium sp. R2A62
Pelagimonas varians DSM 23678
Pelagimonas varians
Planktotalea frisia DSM 23709
Planktotalea frisia SH6-1
Yoonia rosea
Yoonia rosea DSM 29591
Loktanella sp. 1ANDIMAR09
Aliiroseovarius sediminilitoris DSM 29439
Shimia abyssi DSM 100673
Ruegeria meonggei CECT 8411
Ruegeria halocynthiae MOLA R1 13b
Ruegeria halocynthiae
Ruegeria conchae TW15
Ruegeria conchae DSM 29317
Ruegeria conchae
Ruegeria profundi ZGT108
HKCCD7303
HKCCD5849
HKCCD5851
HKCCD7318
HKCCD7296
HKCCD7319
Ruegeria sp. AD91A
HKCCD6238
HKCCD6228
HKCCD6157
HKCCD6119
HKCCD6604
HKCCD6428
HKCCD6181
HKCCD7559
Ruegeria faecimaris DSM 28009
Rhodobacteraceae bacterium KLH11
Ruegeria denitrificans CECT 5091
Ruegeria denitrificans
Ruegeria sp. A3M17
Ruegeria atlantica
Ruegeria sp. AU67
Ruegeria atlantica CECT 4292
Ruegeria atlantica CECT 4293
Ruegeria sp. Alg231-54

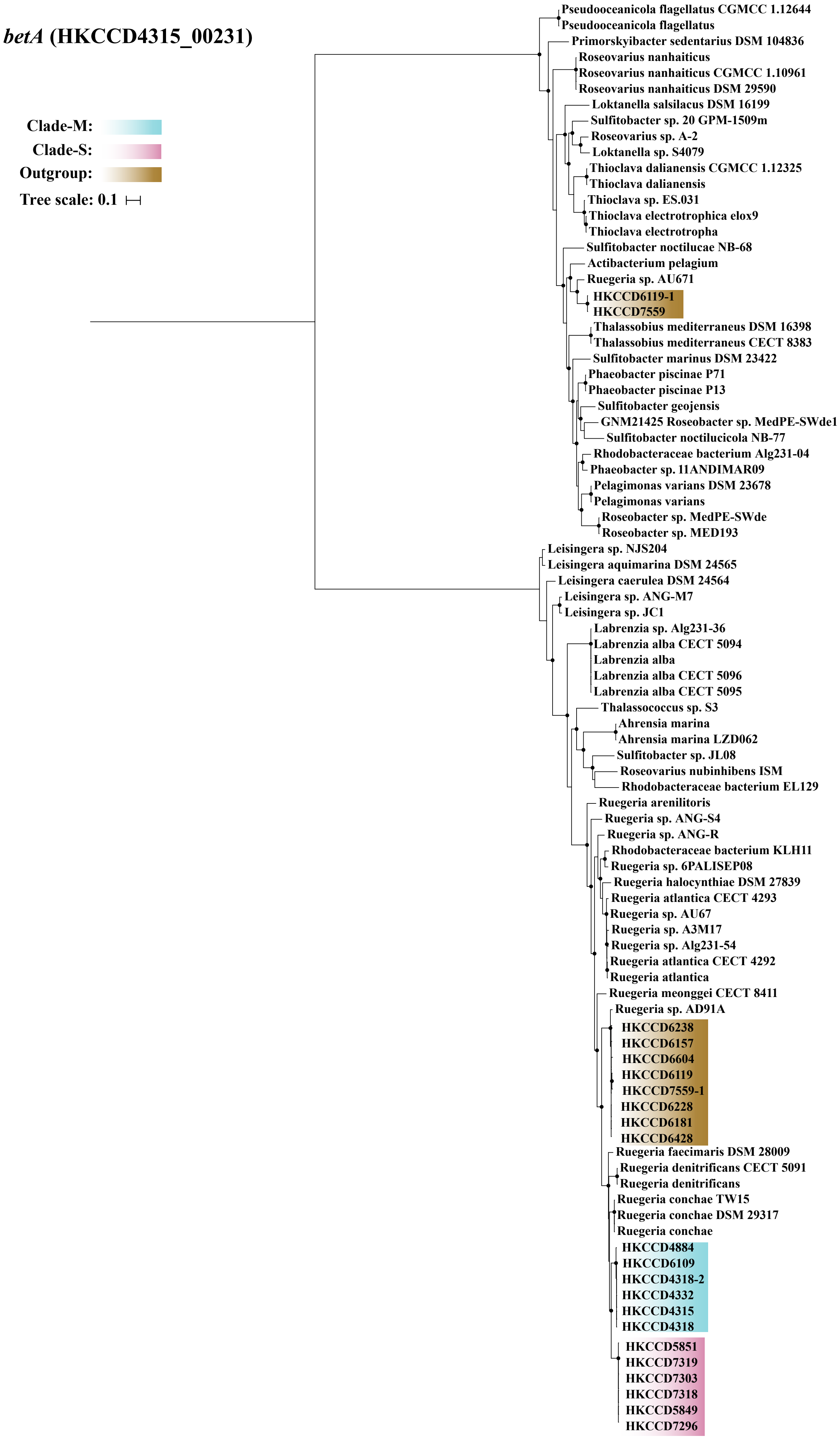

betA (HKCCD4315_00231)
Clade-M:
Clade-S:
Outgroup:
Tree scale: 0.1
Pseudooceanicola flagellatus CGMCC 1.12644
Pseudooceanicola flagellatus
Primorskyibacter sedentarius DSM 104836
Roseovarius nanhaiticus
Roseovarius nanhaiticus CGMCC 1.10961
Roseovarius nanhaiticus DSM 29590
Loktanella salsilacus DSM 16199
Sulfitobacter sp. 20 GPM-1509m
Roseovarius sp. A-2
Loktanella sp. S4079
Thioclava dalianensis CGMCC 1.12325
Thioclava dalianensis
Thioclava sp. ES.031
Thioclava electrotrophica elox9
Thioclava electrotropha
Sulfitobacter noctilucae NB-68
Actibacterium pelagium
Ruegeria sp. AU671
HKCCD6119-1
HKCCD7559
Thalassobius mediterraneus DSM 16398
Thalassobius mediterraneus CECT 8383
Sulfitobacter marinus DSM 23422
Phaeobacter piscinae P71
Phaeobacter piscinae P13
Sulfitobacter geojensis
GNM21425 Roseobacter sp. MedPE-SWde1
Sulfitobacter noctilucicola NB-77
Rhodobacteraceae bacterium Alg231-04
Phaeobacter sp. 11ANDIMAR09
Pelagimonas varians DSM 23678
Pelagimonas varians
Roseobacter sp. MedPE-SWde
Roseobacter sp. MED193
Leisingera sp. NJS204
Leisingera aquimarina DSM 24565
Leisingera caerulea DSM 24564
Leisingera sp. ANG-M7
Leisingera sp. JC1
Labrenzia sp. Alg231-36
Labrenzia alba CECT 5094
Labrenzia alba
Labrenzia alba CECT 5096
Labrenzia alba CECT 5095
Thalassococcus sp. S3
Ahrensia marina
Ahrensia marina LZD062
Sulfitobacter sp. JL08
Roseovarius nubinhibens ISM
Rhodobacteraceae bacterium EL129
Ruegeria arenilitoris
Ruegeria sp. ANG-S4
Ruegeria sp. ANG-R
Rhodobacteraceae bacterium KLH11
Ruegeria sp. 6PALISEP08
Ruegeria halocynthiae DSM 27839
Ruegeria atlantica CECT 4293
Ruegeria sp. AU67
Ruegeria sp. A3M17
Ruegeria sp. Alg231-54
Ruegeria atlantica CECT 4292
Ruegeria atlantica
Ruegeria meonggei CECT 8411
Ruegeria sp. AD91A
HKCCD6238
HKCCD6157
HKCCD6604
HKCCD6119
HKCCD7559-1
HKCCD6228
HKCCD6181
HKCCD6428
Ruegeria faecimaris DSM 28009
Ruegeria denitrificans CECT 5091
Ruegeria denitrificans
Ruegeria conchae TW15
Ruegeria conchae DSM 29317
Ruegeria conchae
HKCCD4884
HKCCD6109
HKCCD4318-2
HKCCD4332
HKCCD4315
HKCCD4318
HKCCD5851
HKCCD7319
HKCCD7303
HKCCD7318
HKCCD5849
HKCCD7296

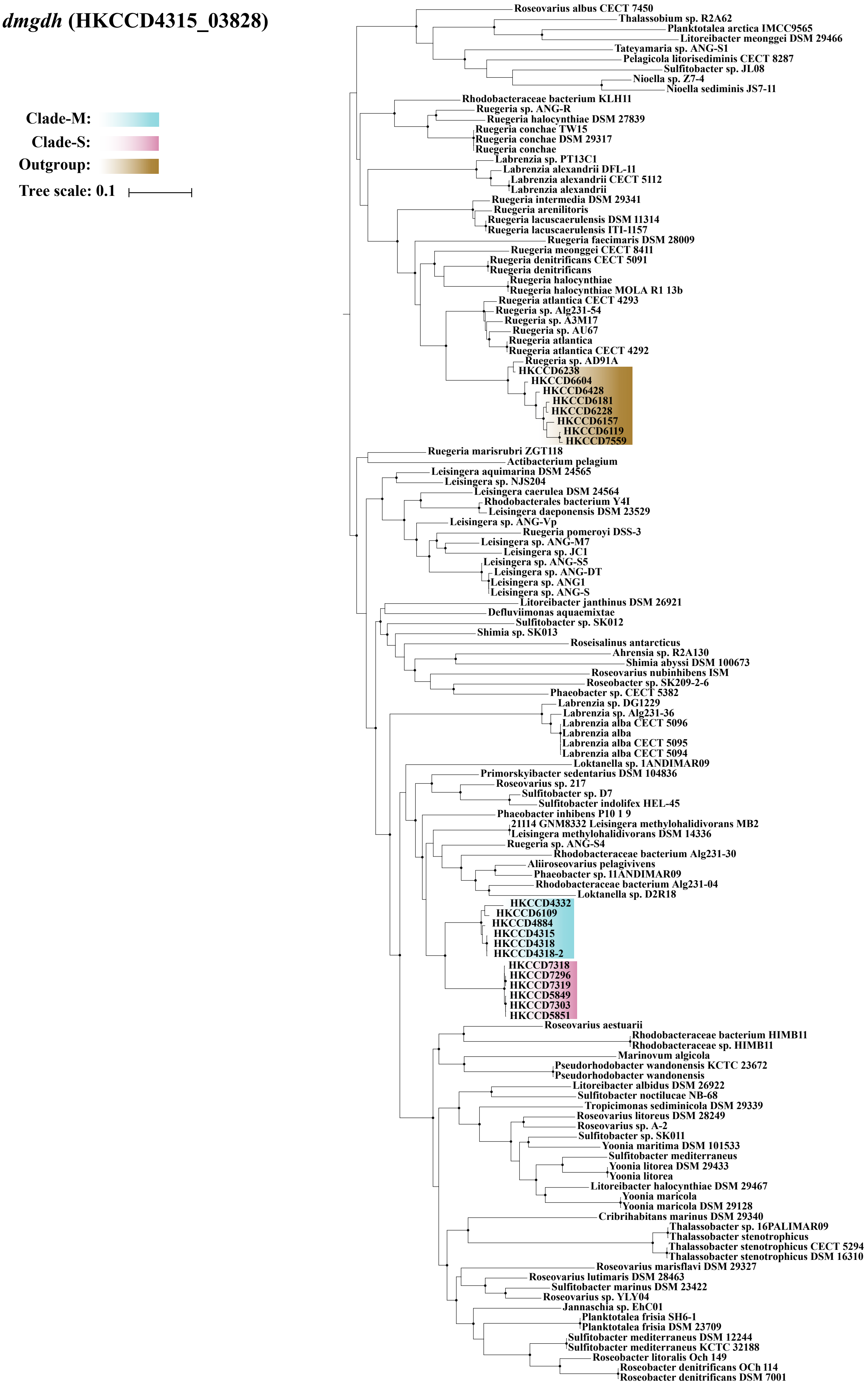
dmgdh (HKCCD4315_03828)
Clade-M:
Clade-S:
Outgroup:
Tree scale: 0.1
Roseovarius albus CECT 7450
Thalassobium sp. R2A62
Planktotalea arctica IMCC9565
Litoreibacter meonggei DSM 29466
Tateyamaria sp. ANG-S1
Pelagicola litorisediminis CECT 8287
Sulfitobacter sp. JL08
Nioella sp. Z7-4
Nioella sediminis JS7-11
Rhodobacteraceae bacterium KLH11
Ruegeria sp. ANG-R
Ruegeria halocynthiae DSM 27839
Ruegeria conchae TW15
Ruegeria conchae DSM 29317
Ruegeria conchae
Labrenzia sp. PT13C1
Labrenzia alexandrii DFL-11
Labrenzia alexandrii CECT 5112
Labrenzia alexandrii
Ruegeria intermedia DSM 29341
Ruegeria arenilitoris
Ruegeria lacuscaerulensis DSM 11314
Ruegeria lacuscaerulensis ITI-1157
Ruegeria faecimaris DSM 28009
Ruegeria meonggei CECT 8411
Ruegeria denitrificans CECT 5091
Ruegeria denitrificans
Ruegeria halocynthiae
Ruegeria halocynthiae MOLA R1 13b
Ruegeria atlantica CECT 4293
Ruegeria sp. Alg231-54
Ruegeria sp. A3M17
Ruegeria sp. AU67
Ruegeria atlantica
Ruegeria atlantica CECT 4292
Ruegeria sp. AD91A
HKCCD6238
HKCCD6604
HKCCD6428
HKCCD6181
HKCCD6228
HKCCD6157
HKCCD6119
HKCCD7559
Ruegeria marisrubri ZGT118
Actibacterium pelagium
Leisingera aquimarina DSM 24565
Leisingera sp. NJS204
Leisingera caerulea DSM 24564
Rhodobacterales bacterium Y4I
Leisingera daeponensis DSM 23529
Leisingera sp. ANG-Vp
Ruegeria pomeroyi DSS-3
Leisingera sp. ANG-M7
Leisingera sp. JC1
Leisingera sp. ANG-S5
Leisingera sp. ANG-DT
Leisingera sp. ANG1
Leisingera sp. ANG-S
Litoreibacter janthinus DSM 26921
Defluviimonas aquaemixtae
Sulfitobacter sp. SK012
Shimia sp. SK013
Roseisalinus antarcticus
Ahrensia sp. R2A130
Shimia abyssi DSM 100673
Roseovarius nubinhibens ISM
Roseobacter sp. SK209-2-6
Phaeobacter sp. CECT 5382
Labrenzia sp. DG1229
Labrenzia sp. Alg231-36
Labrenzia alba CECT 5096
Labrenzia alba
Labrenzia alba CECT 5095
Labrenzia alba CECT 5094
Loktanella sp. 1ANDIMAR09
Primorskyibacter sedentarius DSM 104836
Roseovarius sp. 217
Sulfitobacter sp. D7
Sulfitobacter indolifex HEL-45
Phaeobacter inhibens P10 1 9
21114 GNM8332 Leisingera methylohalidivorans MB2
Leisingera methylohalidivorans DSM 14336
Ruegeria sp. ANG-S4
Rhodobacteraceae bacterium Alg231-30
Aliiroseovarius pelagivivens
Phaeobacter sp. 11ANDIMAR09
Rhodobacteraceae bacterium Alg231-04
Loktanella sp. D2R18
HKCCD4332
HKCCD6109
HKCCD4884
HKCCD4315
HKCCD4318
HKCCD4318-2
HKCCD7318
HKCCD7296
HKCCD7319
HKCCD5849
HKCCD7303
HKCCD5851
Roseovarius aestuarii
Rhodobacteraceae bacterium HIMB11
Rhodobacteraceae sp. HIMB11
Marinovum algicola
Pseudorhodobacter wandonensis KCTC 23672
Pseudorhodobacter wandonensis
Litoreibacter albidus DSM 26922
Sulfitobacter noctilucae NB-68
Tropicimonas sediminicola DSM 29339
Roseovarius litoreus DSM 28249
Roseovarius sp. A-2
Sulfitobacter sp. SK011
Yoonia maritima DSM 101533
Sulfitobacter mediterraneus
Yoonia litorea DSM 29433
Yoonia litorea
Litoreibacter halocynthiae DSM 29467
Yoonia maricola
Yoonia maricola DSM 29128
Cribrihabitans marinus DSM 29340
Thalassobacter sp. 16PALIMAR09
Thalassobacter stenotrophicus
Thalassobacter stenotrophicus CECT 5294
Thalassobacter stenotrophicus DSM 16310
Roseovarius marisflavi DSM 29327
Roseovarius lutimaris DSM 28463
Sulfitobacter marinus DSM 23422
Roseovarius sp. YLY04
Jannaschia sp. EhC01
Planktotalea frisia SH6-1
Planktotalea frisia DSM 23709
Sulfitobacter mediterraneus DSM 12244
Sulfitobacter mediterraneus KCTC 32188
Roseobacter litoralis Och 149
Roseobacter denitrificans OCh 114
Roseobacter denitrificans DSM 7001

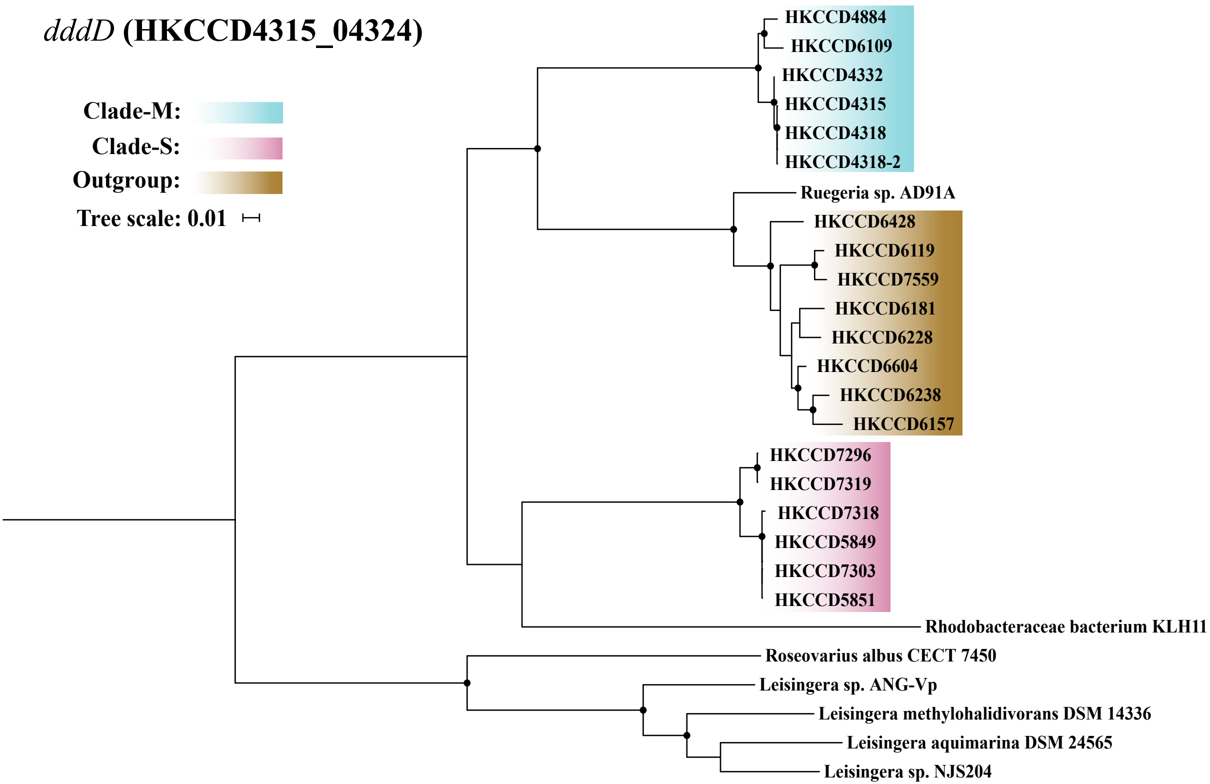


**Figure S6.** The maximum likelihood phylogeny of the core gene families *dddD* involved in DMSP utilization constructed by IQ-TREE v1.6.5. Solid circles at the nodes indicate that the ultrafast bootstrap support value of the branch is ≥ 80%. The scale bar indicates number of substitutions per site. The members of clade-M, clade-S and outgroup clade were shadowed with blue, pink and gold, respectively.

**Figure S7.** The maximum likelihood phylogeny of the core gene families *tauA*, *tauB* and *tauC* involved in taurine utilization constructed by IQ-TREE v1.6.5. Solid circles at the nodes indicate that the ultrafast bootstrap support value of the branch is ≥ 80%. The scale bar indicates number of substitutions per site. The members of clade-M, clade-S and outgroup clade were shadowed with blue, pink and gold, respectively.

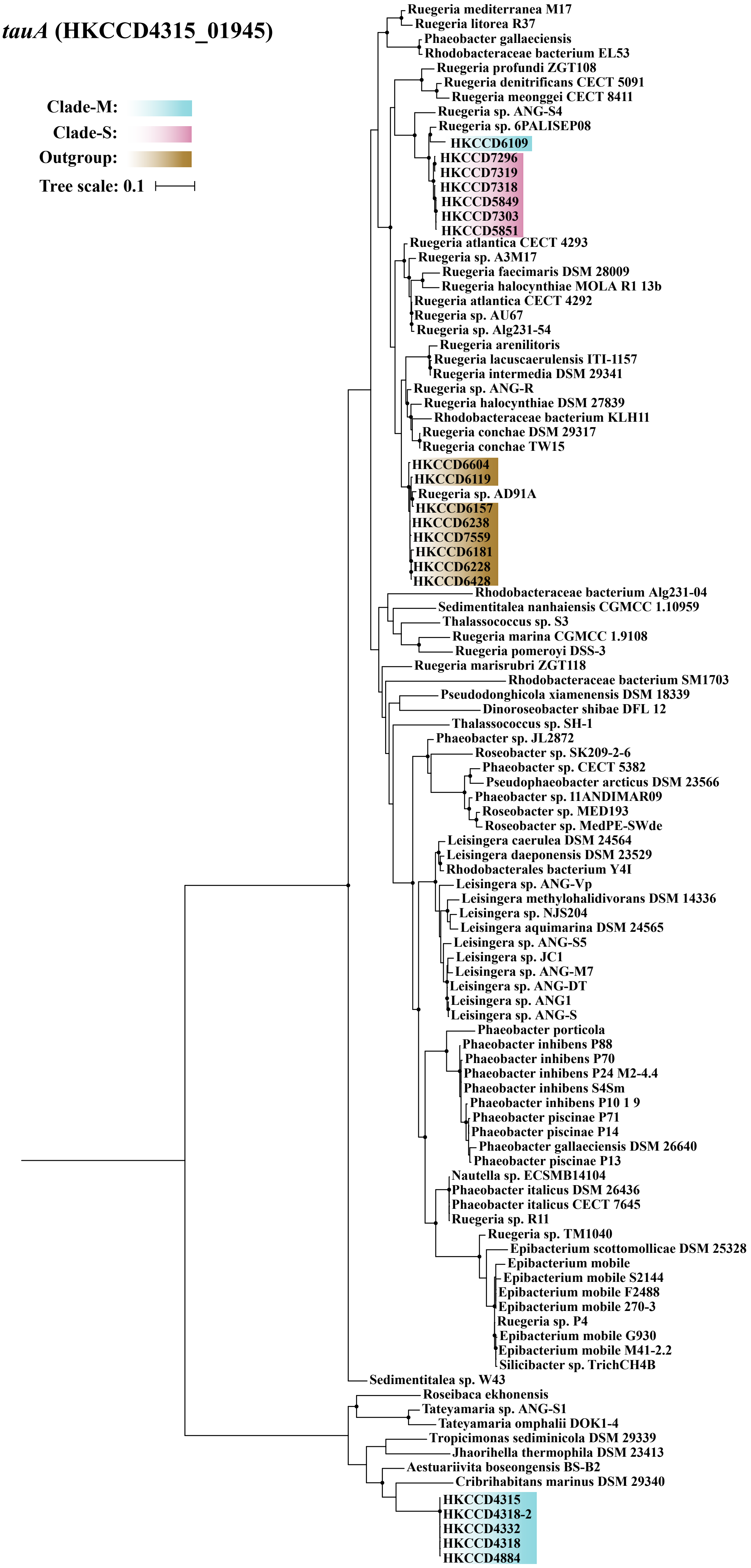

tauA (HKCCD4315_01945)
Clade-M:
Clade-S:
Outgroup:
Tree scale: 0.1
Ruegeria mediterranea M17
Ruegeria litorea R37
Phaeobacter gallaeciensis
Rhodobacteraceae bacterium EL53
Ruegeria profundi ZGT108
Ruegeria denitrificans CECT 5091
Ruegeria meonggei CECT 8411
Ruegeria sp. ANG-S4
Ruegeria sp. 6PALISEP08
HKCCD6109
HKCCD7296
HKCCD7319
HKCCD7318
HKCCD5849
HKCCD7303
HKCCD5851
Ruegeria atlantica CECT 4293
Ruegeria sp. A3M17
Ruegeria faecimaris DSM 28009
Ruegeria halocynthiae MOLA R1 13b
Ruegeria atlantica CECT 4292
Ruegeria sp. AU67
Ruegeria sp. Alg231-54
Ruegeria arenilitoris
Ruegeria lacuscaerulensis ITI-1157
Ruegeria intermedia DSM 29341
Ruegeria sp. ANG-R
Ruegeria halocynthiae DSM 27839
Rhodobacteraceae bacterium KLH11
Ruegeria conchae DSM 29317
Ruegeria conchae TW15
HKCCD6604
HKCCD6119
Ruegeria sp. AD91A
HKCCD6157
HKCCD6238
HKCCD7559
HKCCD6181
HKCCD6228
HKCCD6428
Rhodobacteraceae bacterium Alg231-04
Sedimentitalea nanhaiensis CGMCC 1.10959
Thalassococcus sp. S3
Ruegeria marina CGMCC 1.9108
Ruegeria pomeroyi DSS-3
Ruegeria marisrubri ZGT118
Rhodobacteraceae bacterium SM1703
Pseudodonghicola xiamenensis DSM 18339
Dinoroseobacter shibae DFL 12
Thalassococcus sp. SH-1
Phaeobacter sp. JL2872
Roseobacter sp. SK209-2-6
Phaeobacter sp. CECT 5382
Pseudophaeobacter arcticus DSM 23566
Phaeobacter sp. 11ANDIMAR09
Roseobacter sp. MED193
Roseobacter sp. MedPE-SWde
Leisingera caerulea DSM 24564
Leisingera daeponensis DSM 23529
Rhodobacterales bacterium Y4I
Leisingera sp. ANG-Vp
Leisingera methylohalidivorans DSM 14336
Leisingera sp. NJS204
Leisingera aquimarina DSM 24565
Leisingera sp. ANG-S5
Leisingera sp. JC1
Leisingera sp. ANG-M7
Leisingera sp. ANG-DT
Leisingera sp. ANG1
Leisingera sp. ANG-S
Phaeobacter porticola
Phaeobacter inhibens P88
Phaeobacter inhibens P70
Phaeobacter inhibens P24 M2-4.4
Phaeobacter inhibens S4Sm
Phaeobacter inhibens P10 1 9
Phaeobacter piscinae P71
Phaeobacter piscinae P14
Phaeobacter gallaeciensis DSM 26640
Phaeobacter piscinae P13
Nautella sp. ECSMB14104
Phaeobacter italicus DSM 26436
Phaeobacter italicus CECT 7645
Ruegeria sp. R11
Ruegeria sp. TM1040
Epibacterium scottomollicae DSM 25328
Epibacterium mobile
Epibacterium mobile S2144
Epibacterium mobile F2488
Epibacterium mobile 270-3
Ruegeria sp. P4
Epibacterium mobile G930
Epibacterium mobile M41-2.2
Silicibacter sp. TrichCH4B
Sedimentitalea sp. W43
Roseibaca ekhonensis
Tateyamaria sp. ANG-S1
Tateyamaria omphalii DOK1-4
Tropicimonas sediminicola DSM 29339
Jhaorihella thermophila DSM 23413
Aestuariivita boseongensis BS-B2
Cribrihabitans marinus DSM 29340
HKCCD4315
HKCCD4318-2
HKCCD4332
HKCCD4318
HKCCD4884

### *tauB* (HKCCD4315_01947)

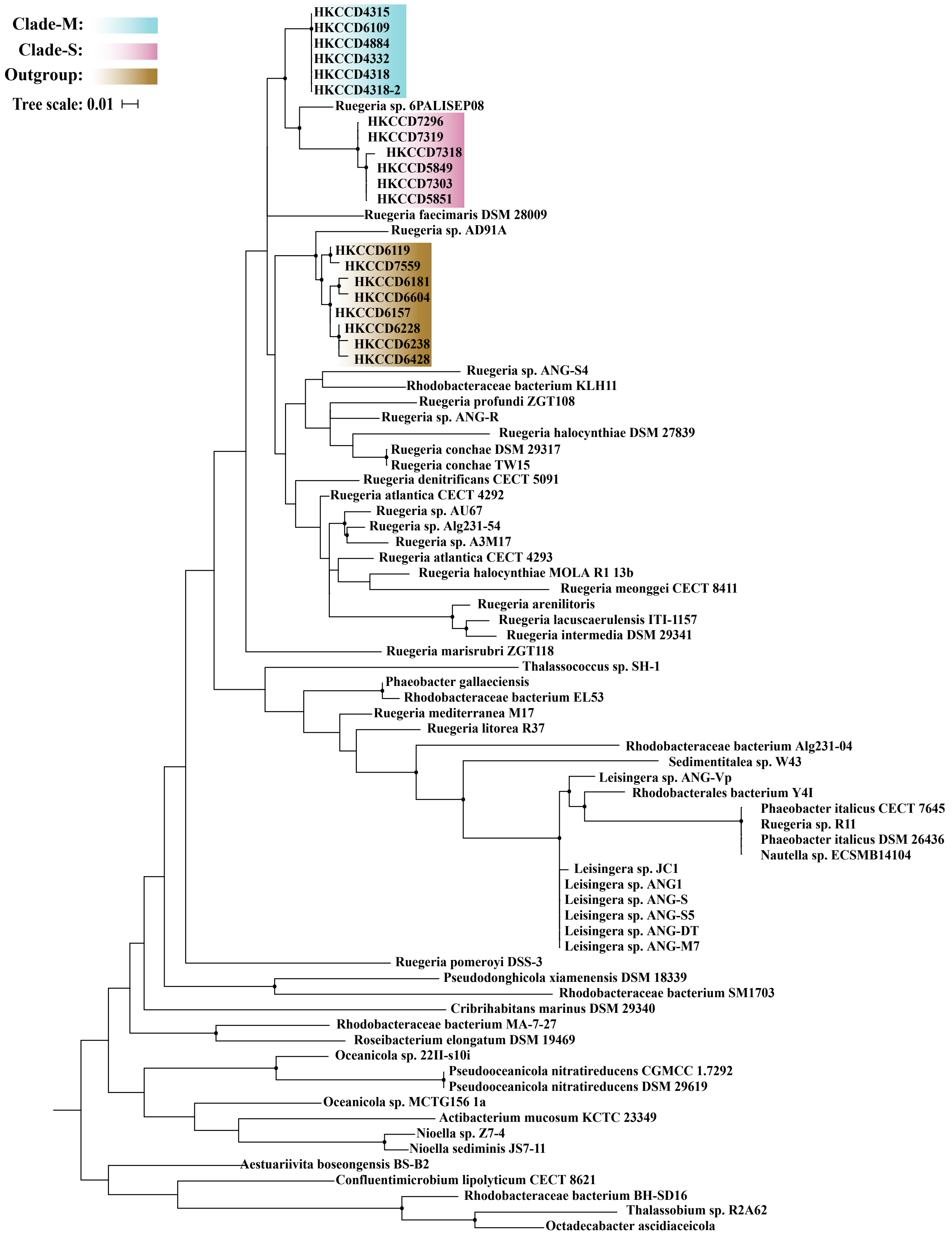

### *tauC* (HKCCD4315_01944)

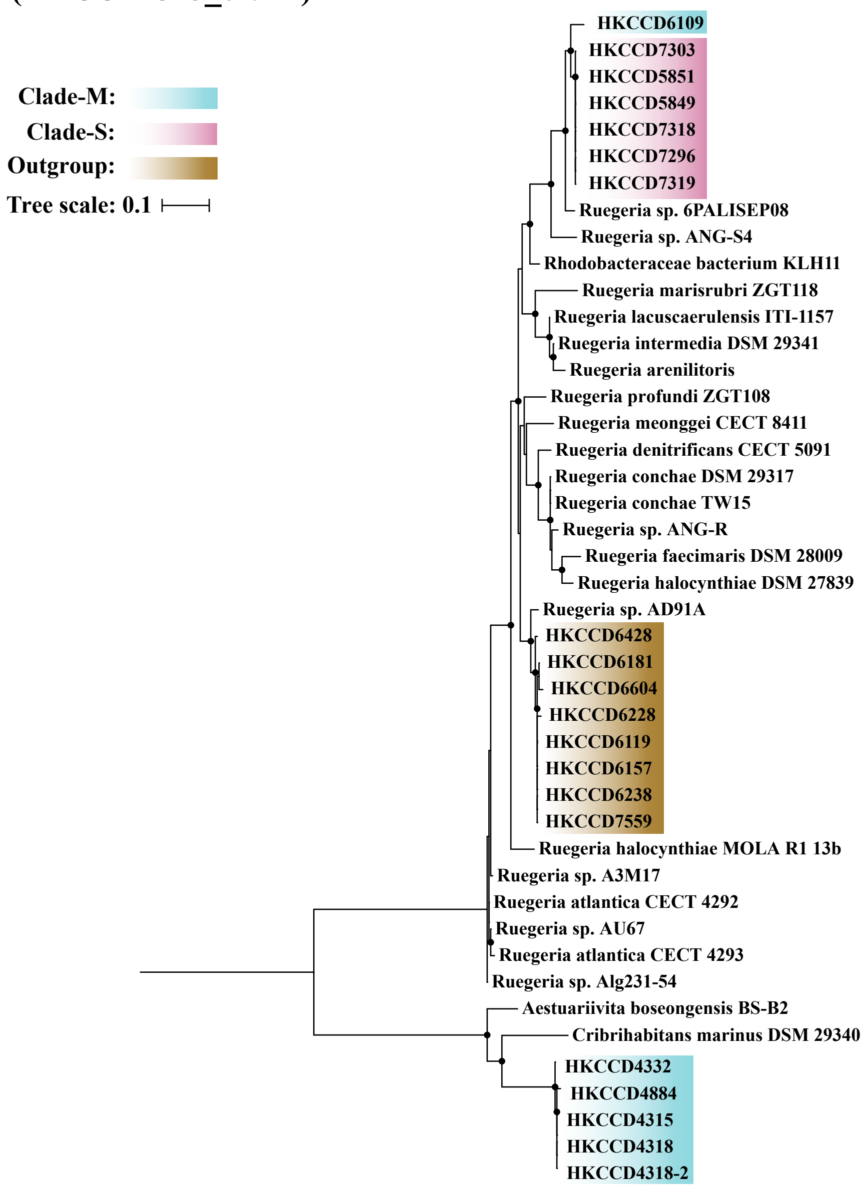

**Figure S8.** The maximum likelihood phylogeny of the core gene families *coxG*, *coxF*, *coxE*, *coxD*, *coxL*, *coxS*, *coxM* and *coxC* involved in CO oxidation constructed by IQ-TREE v1.6.5. Solid circles at the nodes indicate that the ultrafast bootstrap support value of the branch is ≥ 80%. The scale bar indicates number substitutions per site. The members of clade-M, clade-S and outgroup clade were shadowed with blue, pink and gold, respectively.

*coxG* (HKCCD4315_03676)

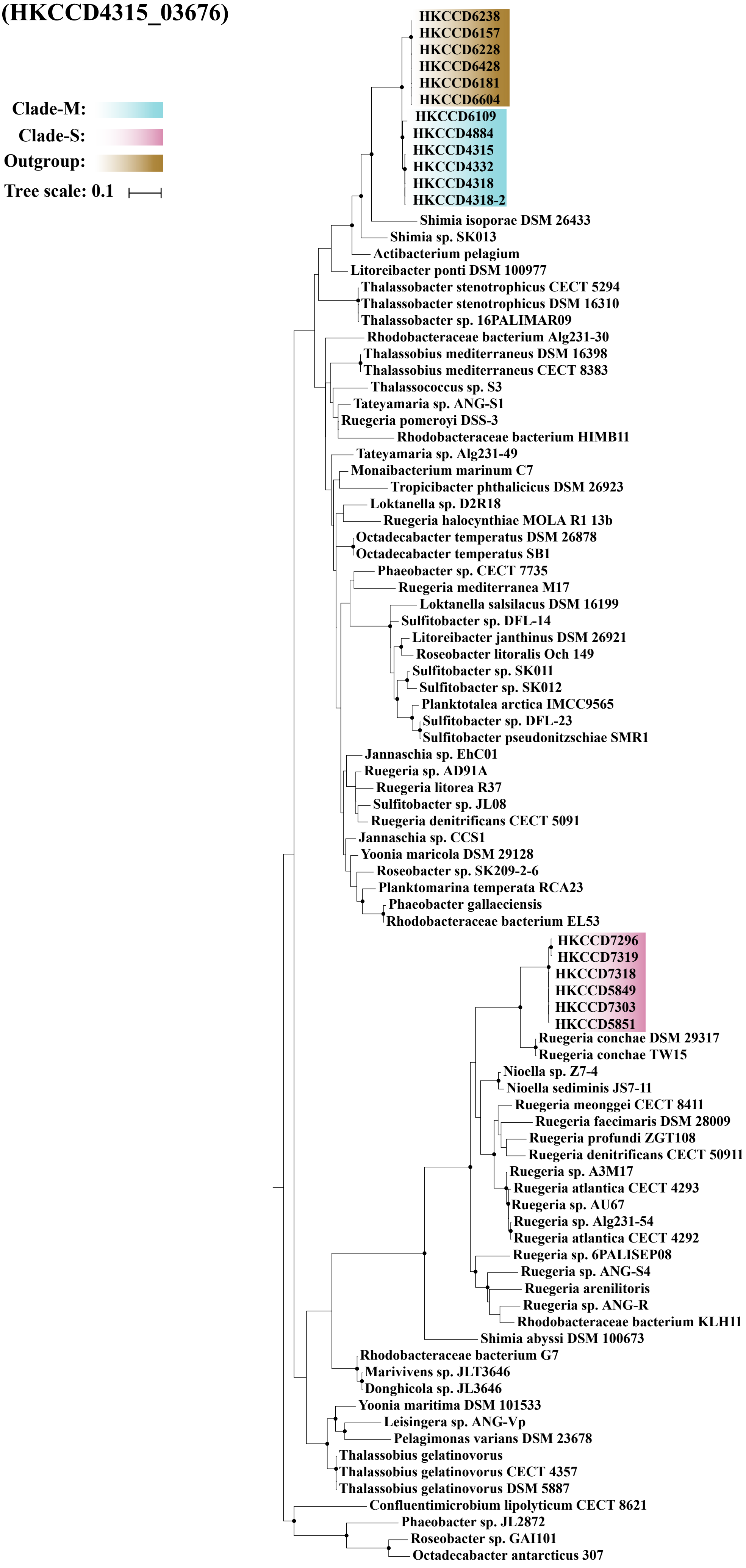

### *coxF* (HKCCD4315_03678)

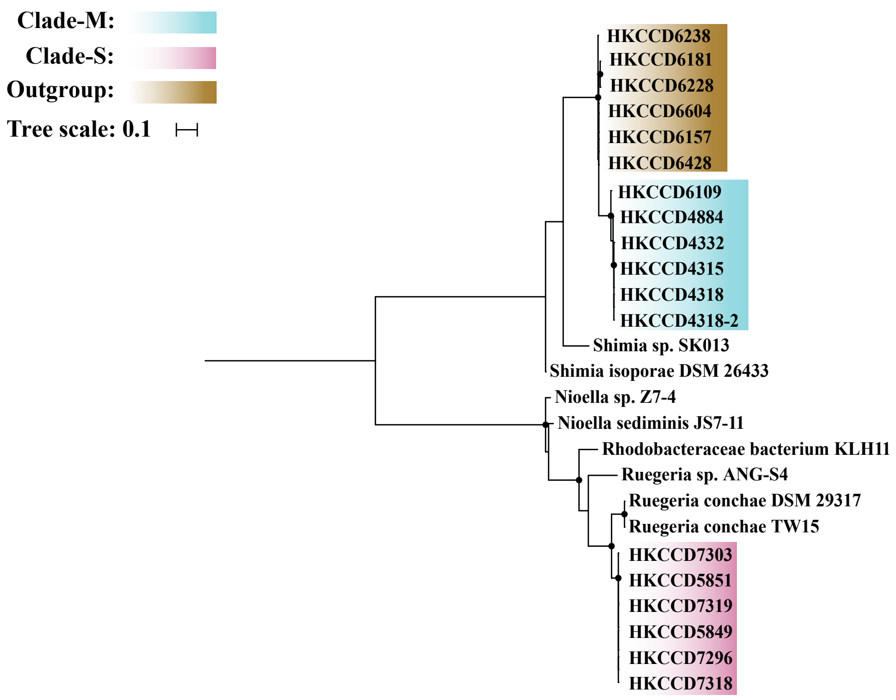


### *coxE* (HKCCD4315_03679)

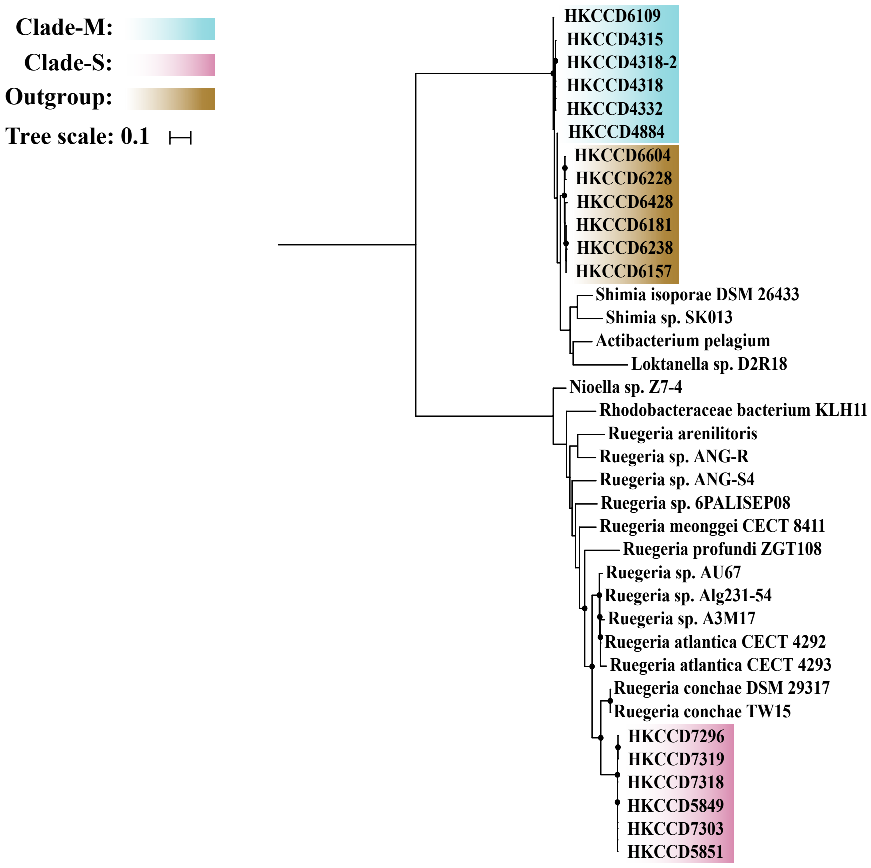

*coxD* (HKCCD4315_03680)

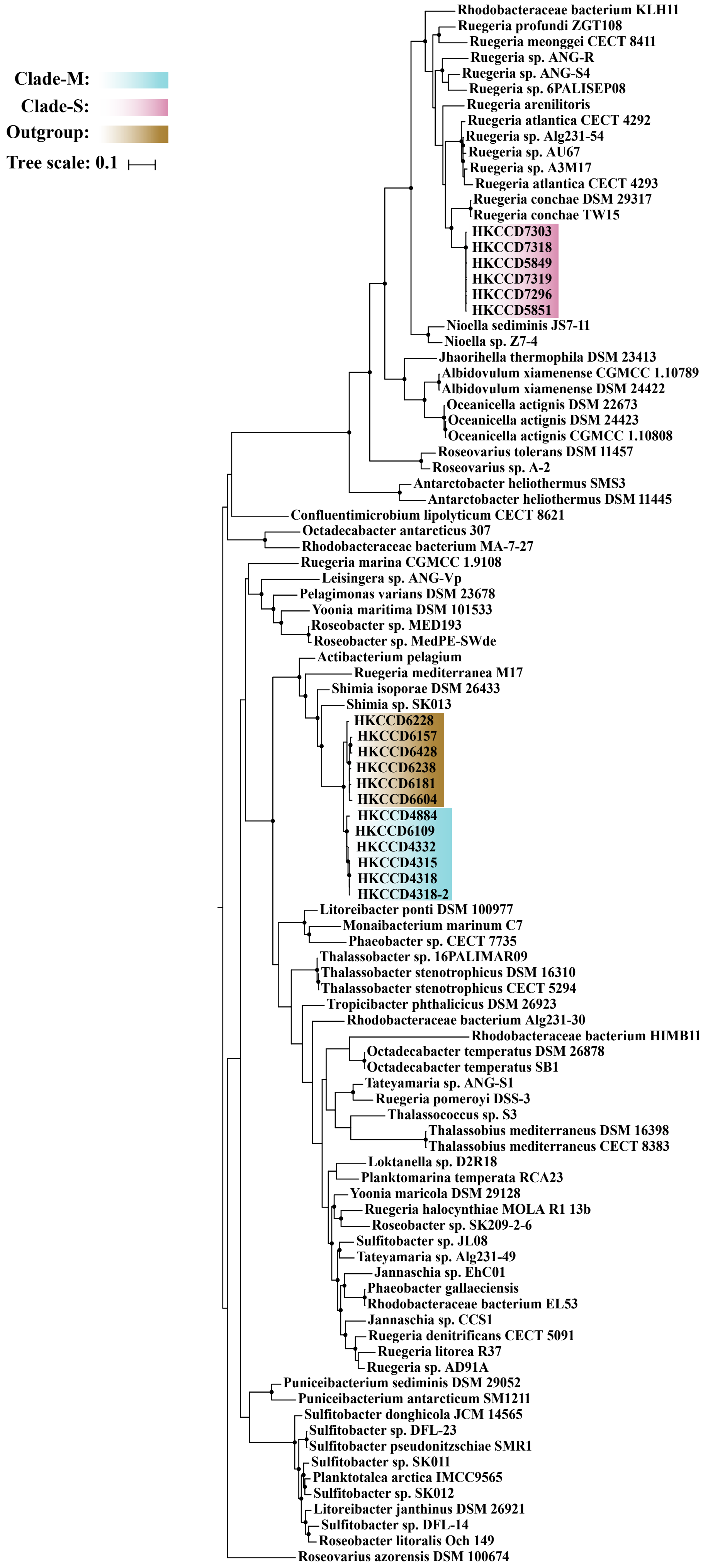

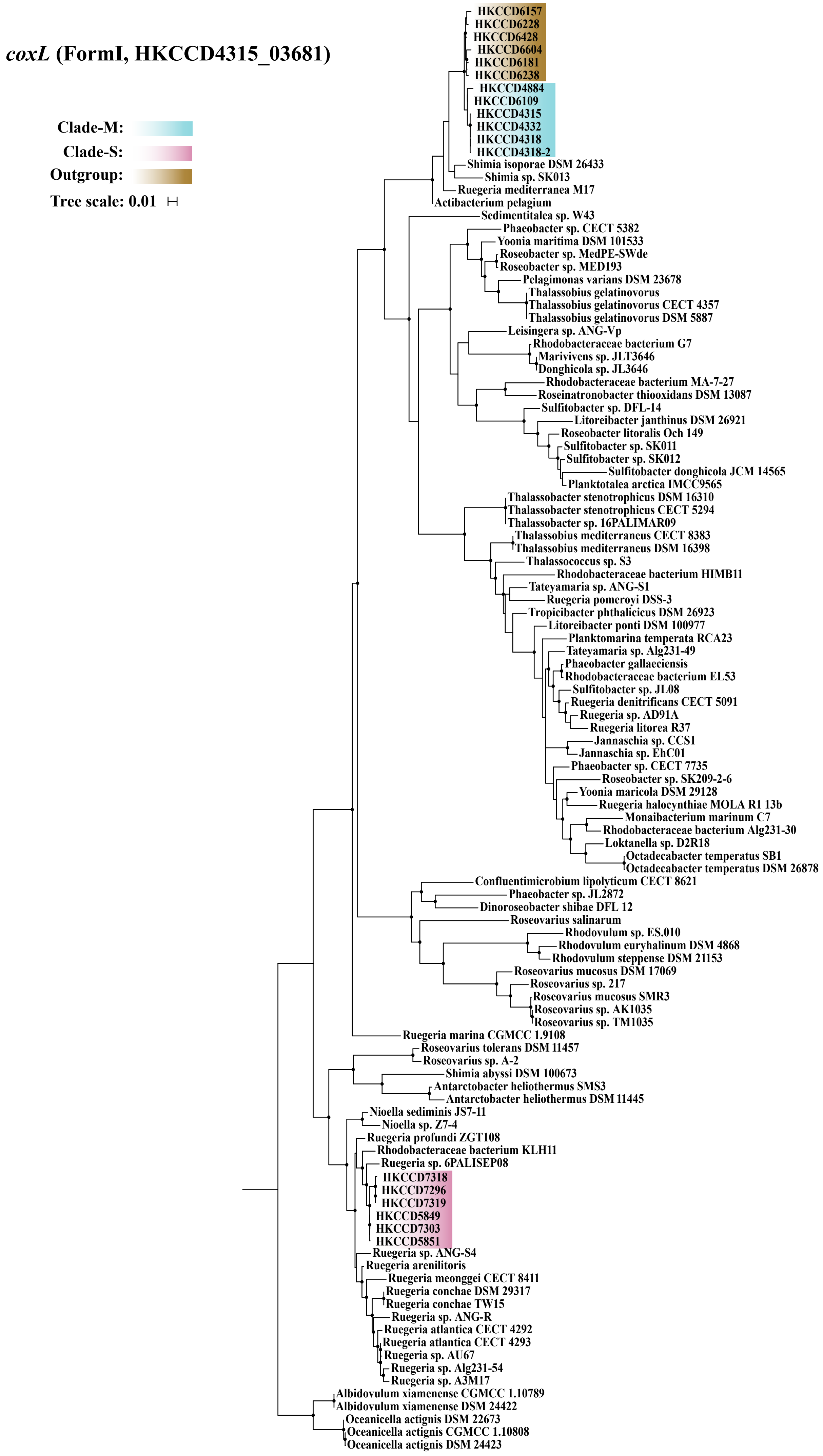
coxL (FormI, HKCCD4315_03681)
Clade-M:
Clade-S:
Outgroup:
Tree scale: 0.01
HKCCD6157
HKCCD6228
HKCCD6428
HKCCD6604
HKCCD6181
HKCCD6238
HKCCD4884
HKCCD6109
HKCCD4315
HKCCD4332
HKCCD4318
HKCCD4318-2
Shimia isoporae DSM 26433
Shimia sp. SK013
Ruegeria mediterranea M17
Actibacterium pelagium
Sedimentitalea sp. W43
Phaeobacter sp. CECT 5382
Yoonia maritima DSM 101533
Roseobacter sp. MedPE-SWde
Roseobacter sp. MED193
Pelagimonas varians DSM 23678
Thalassobius gelatinovorus
Thalassobius gelatinovorus CECT 4357
Thalassobius gelatinovorus DSM 5887
Leisingera sp. ANG-Vp
Rhodobacteraceae bacterium G7
Marivivens sp. JLT3646
Donghicola sp. JL3646
Rhodobacteraceae bacterium MA-7-27
Roseinatronobacter thiooxidans DSM 13087
Sulfitobacter sp. DFL-14
Litoreibacter janthinus DSM 26921
Roseobacter litoralis Och 149
Sulfitobacter sp. SK011
Sulfitobacter sp. SK012
Sulfitobacter donghicola JCM 14565
Planktotalea arctica IMCC9565
Thalassobacter stenotrophicus DSM 16310
Thalassobacter stenotrophicus CECT 5294
Thalassobacter sp. 16PALIMAR09
Thalassobius mediterraneus CECT 8383
Thalassobius mediterraneus DSM 16398
Thalassococcus sp. S3
Rhodobacteraceae bacterium HIMB11
Tateyamaria sp. ANG-S1
Ruegeria pomeroyi DSS-3
Tropicibacter phthalicicus DSM 26923
Litoreibacter ponti DSM 100977
Planktomarina temperata RCA23
Tateyamaria sp. Alg231-49
Phaeobacter gallaeciensis
Rhodobacteraceae bacterium EL53
Sulfitobacter sp. JL08
Ruegeria denitrificans CECT 5091
Ruegeria sp. AD91A
Ruegeria litorea R37
Jannaschia sp. CCS1
Jannaschia sp. EhC01
Phaeobacter sp. CECT 7735
Roseobacter sp. SK209-2-6
Yoonia maricola DSM 29128
Ruegeria halocynthiae MOLA R1 13b
Monaibacterium marinum C7
Rhodobacteraceae bacterium Alg231-30
Loktanella sp. D2R18
Octadecabacter temperatus SB1
Octadecabacter temperatus DSM 26878
Confluentimicrobium lipolyticum CECT 8621
Phaeobacter sp. JL2872
Dinoroseobacter shibae DFL 12
Roseovarius salinarum
Rhodovulum sp. ES.010
Rhodovulum euryhalinum DSM 4868
Rhodovulum steppense DSM 21153
Roseovarius mucosus DSM 17069
Roseovarius sp. 217
Roseovarius mucosus SMR3
Roseovarius sp. AK1035
Roseovarius sp. TM1035
Ruegeria marina CGMCC 1.9108
Roseovarius tolerans DSM 11457
Roseovarius sp. A-2
Shimia abyssi DSM 100673
Antarctobacter heliothermus SMS3
Antarctobacter heliothermus DSM 11445
Nioella sediminis JS7-11
Nioella sp. Z7-4
Ruegeria profundi ZGT108
Rhodobacteraceae bacterium KLH11
Ruegeria sp. 6PALISEP08
HKCCD7318
HKCCD7296
HKCCD7319
HKCCD5849
HKCCD7303
HKCCD5851
Ruegeria sp. ANG-S4
Ruegeria arenilitoris
Ruegeria meonggei CECT 8411
Ruegeria conchae DSM 29317
Ruegeria conchae TW15
Ruegeria sp. ANG-R
Ruegeria atlantica CECT 4292
Ruegeria atlantica CECT 4293
Ruegeria sp. AU67
Ruegeria sp. Alg231-54
Ruegeria sp. A3M17
Albidovulum xiamenense CGMCC 1.10789
Albidovulum xiamenense DSM 24422
Oceanicella actignis DSM 22673
Oceanicella actignis CGMCC 1.10808
Oceanicella actignis DSM 24423

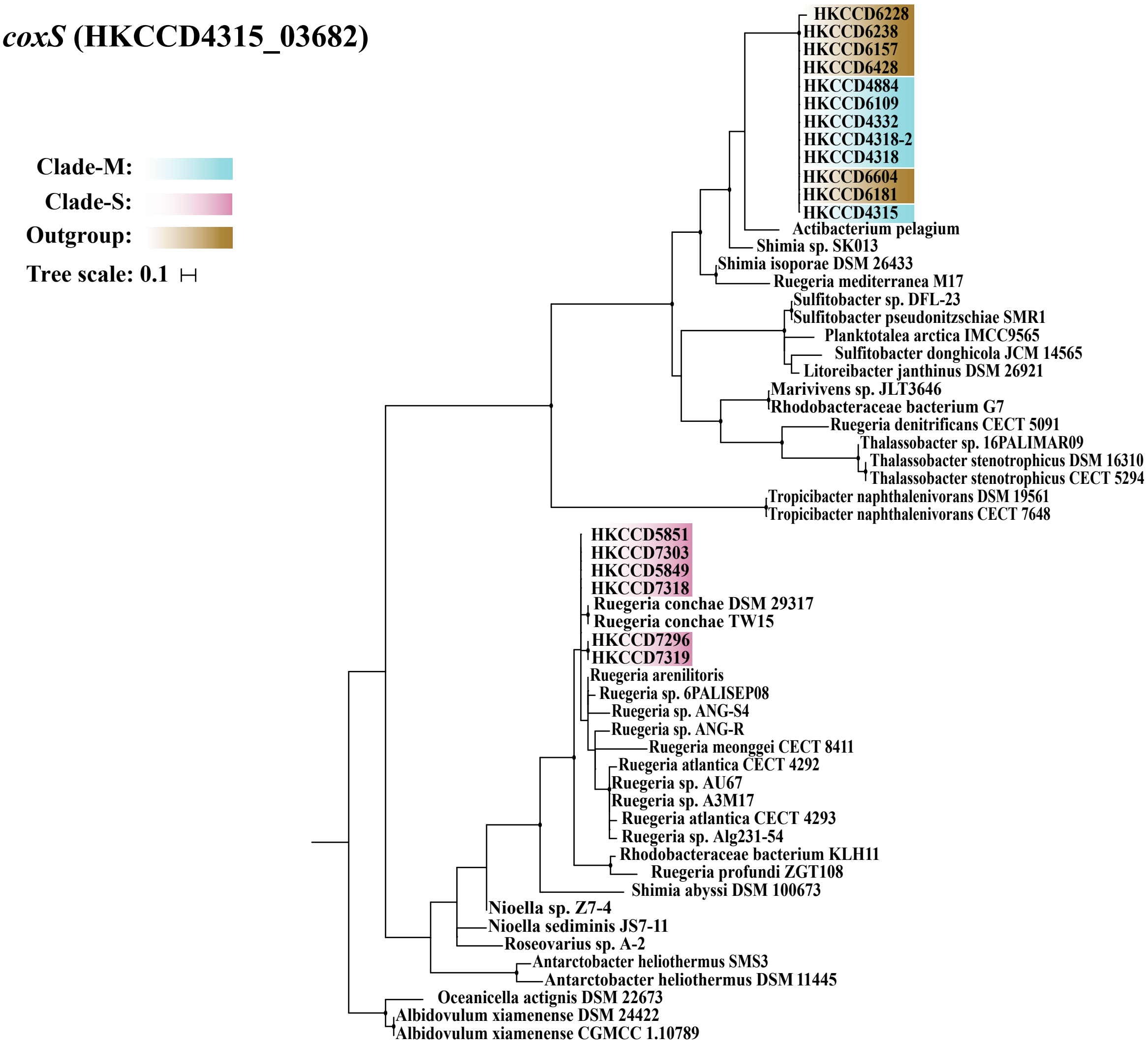


### *coxC* (HKCCD4315_03684)

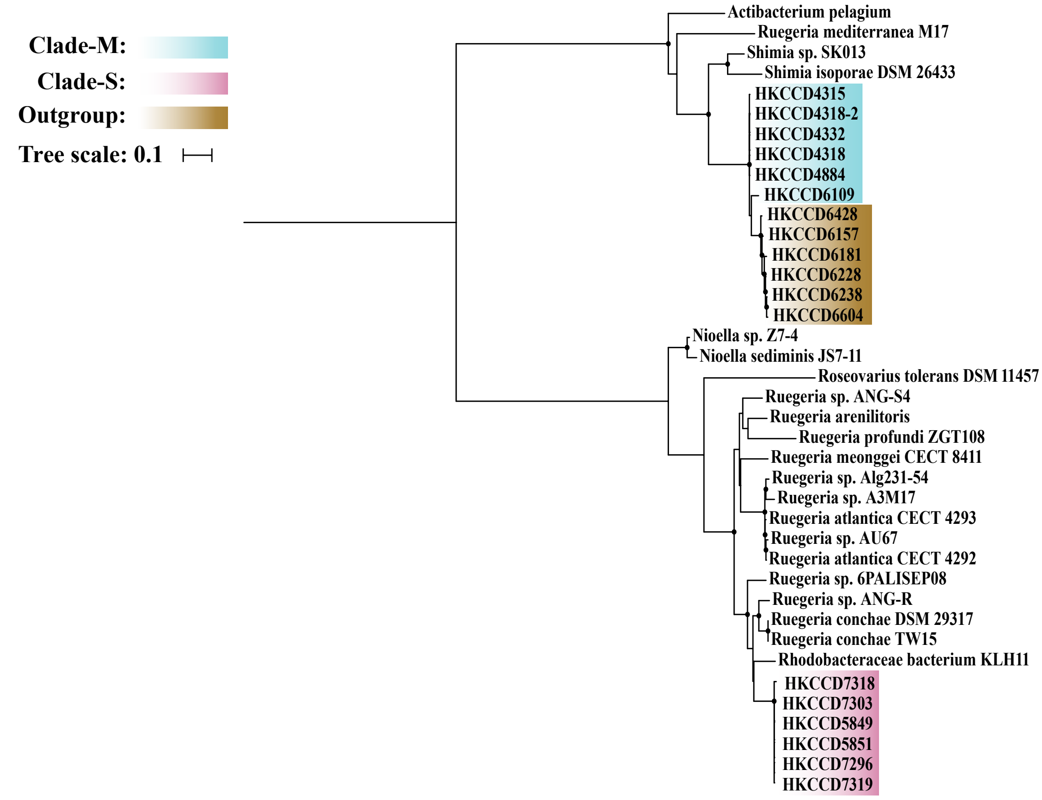

### *coxM* (HKCCD4315_03683)

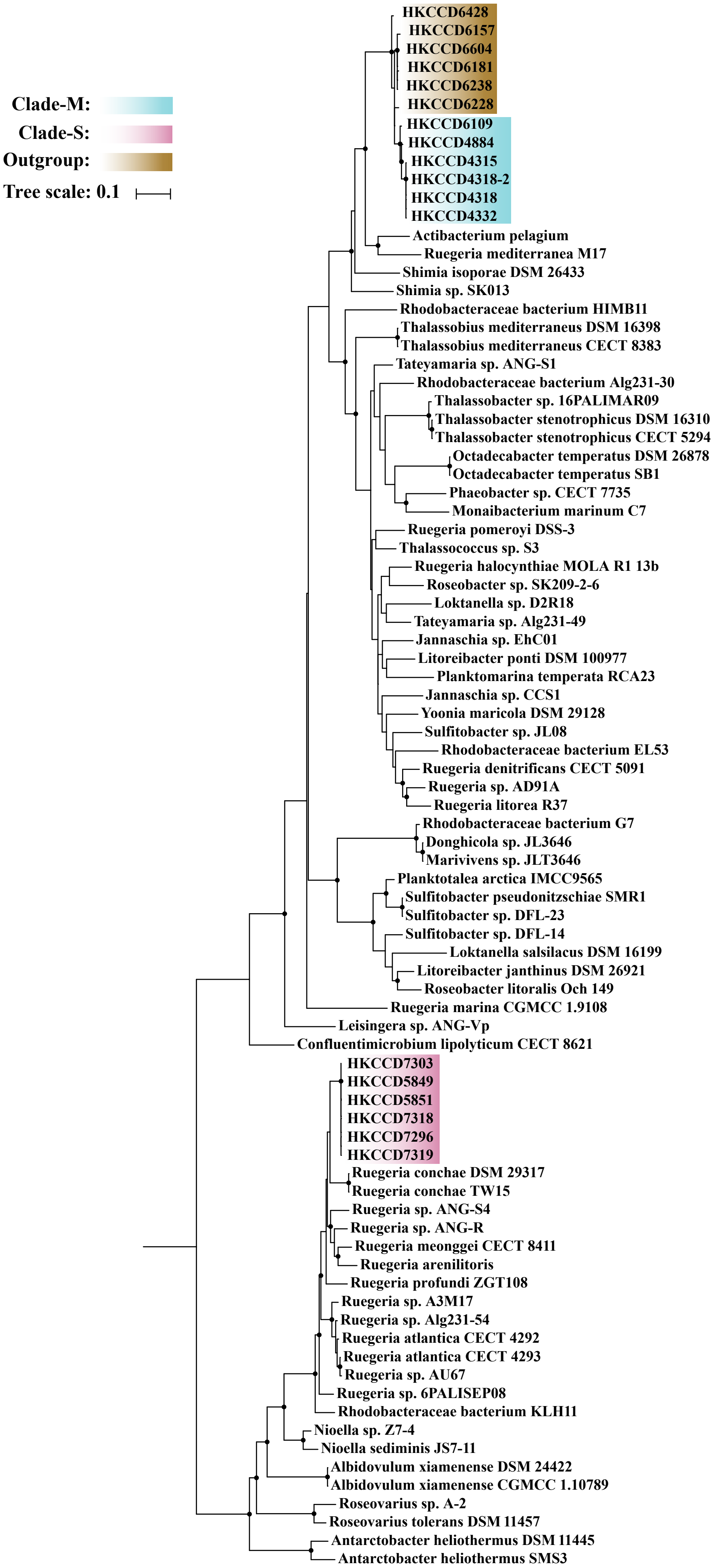

**Figure S9.** The maximum likelihood phylogeny of the core gene families *dmsA*, *dmsB* and *dmsC* involved in anaerobic respiration constructed by IQ-TREE v1.6.5. Solid circles at the nodes indicate that the ultrafast bootstrap support value of the branch is ≥ 80%. The scale bar indicates number of substitutions per site. The members of clade-M, clade-S and outgroup clade were shadowed with blue, pink and gold, respectively.

### *dmsA* (HKCCD4315_03090)

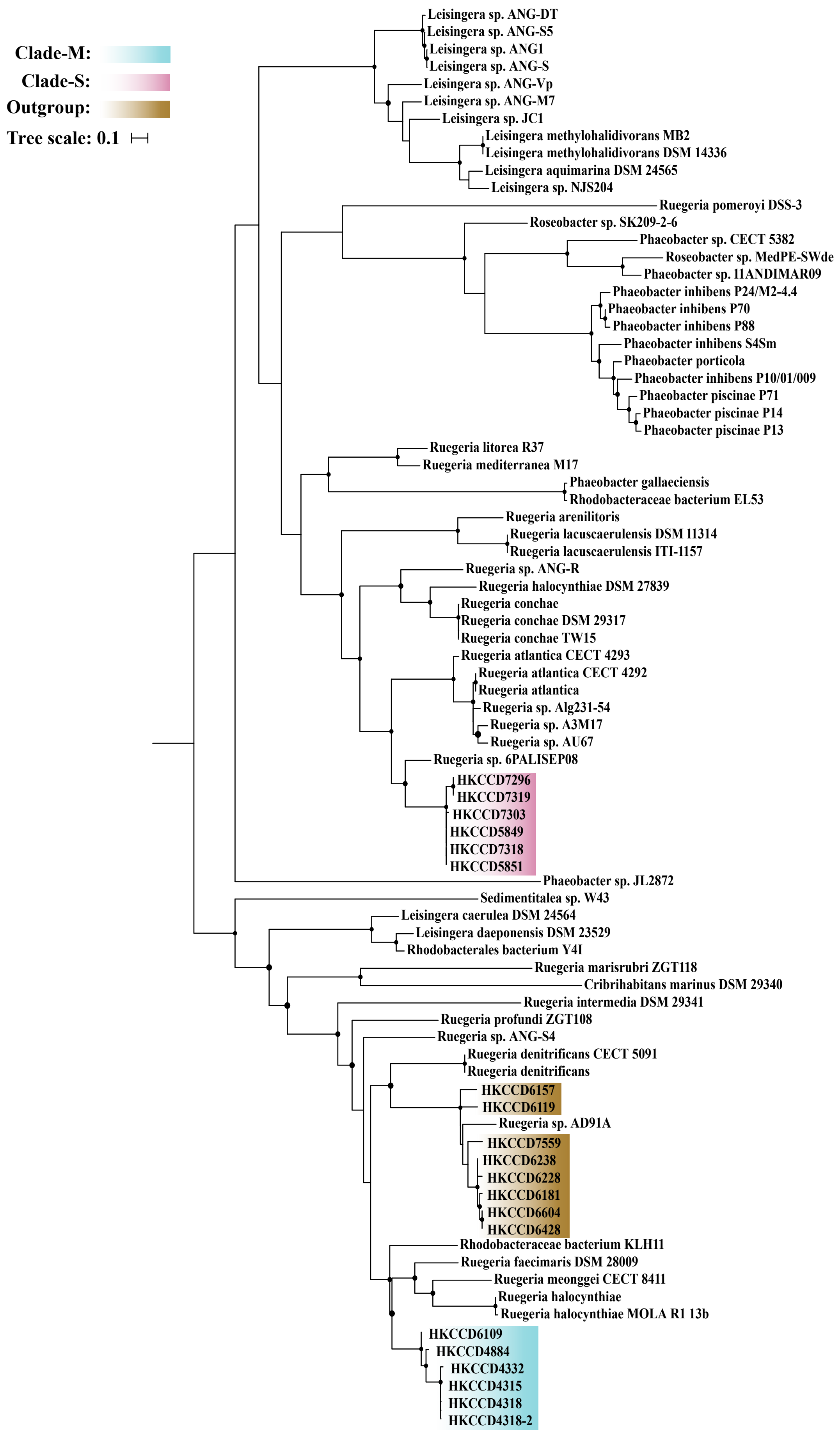

*dmsB* (HKCCD4315_03091)

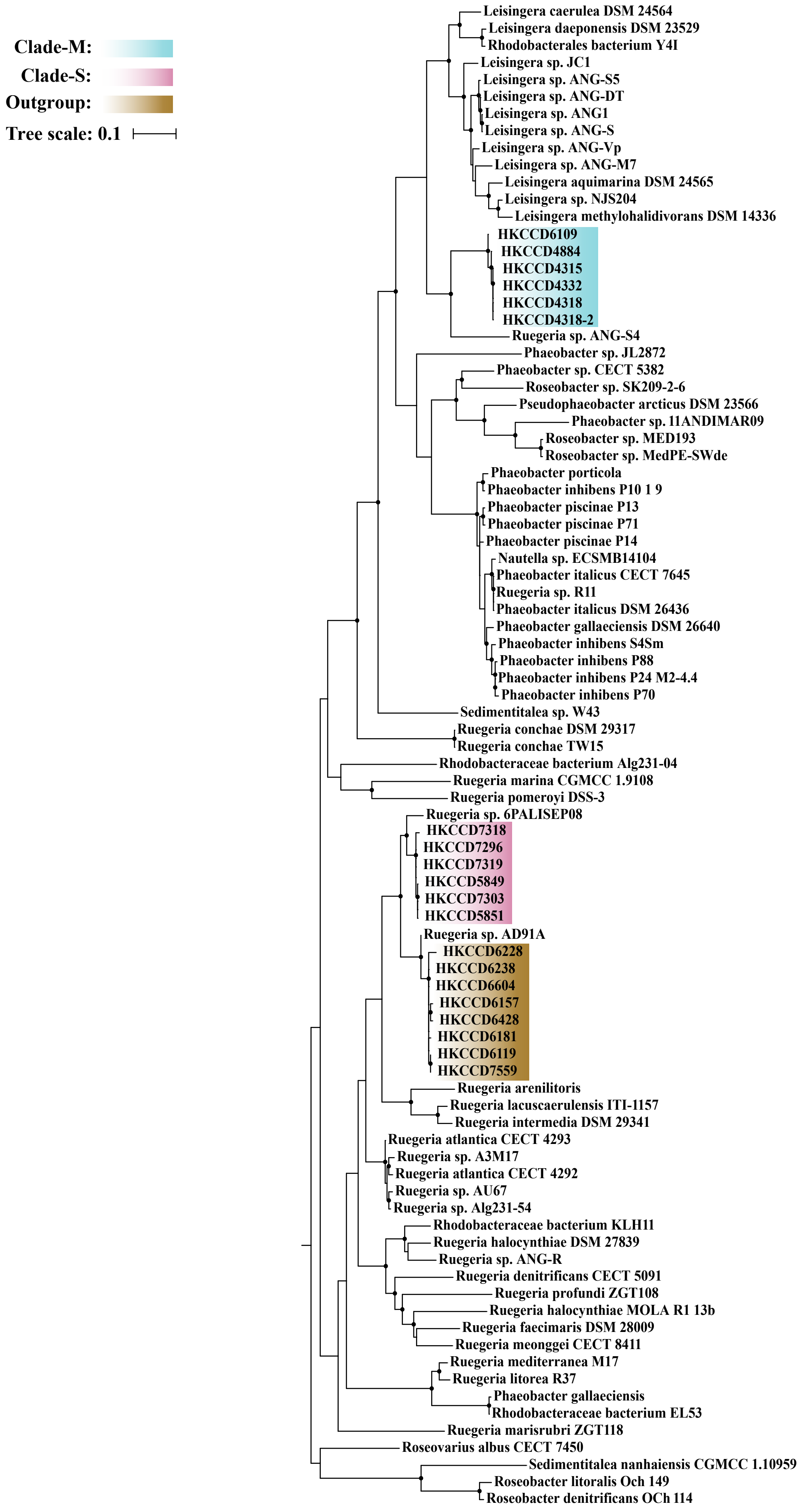

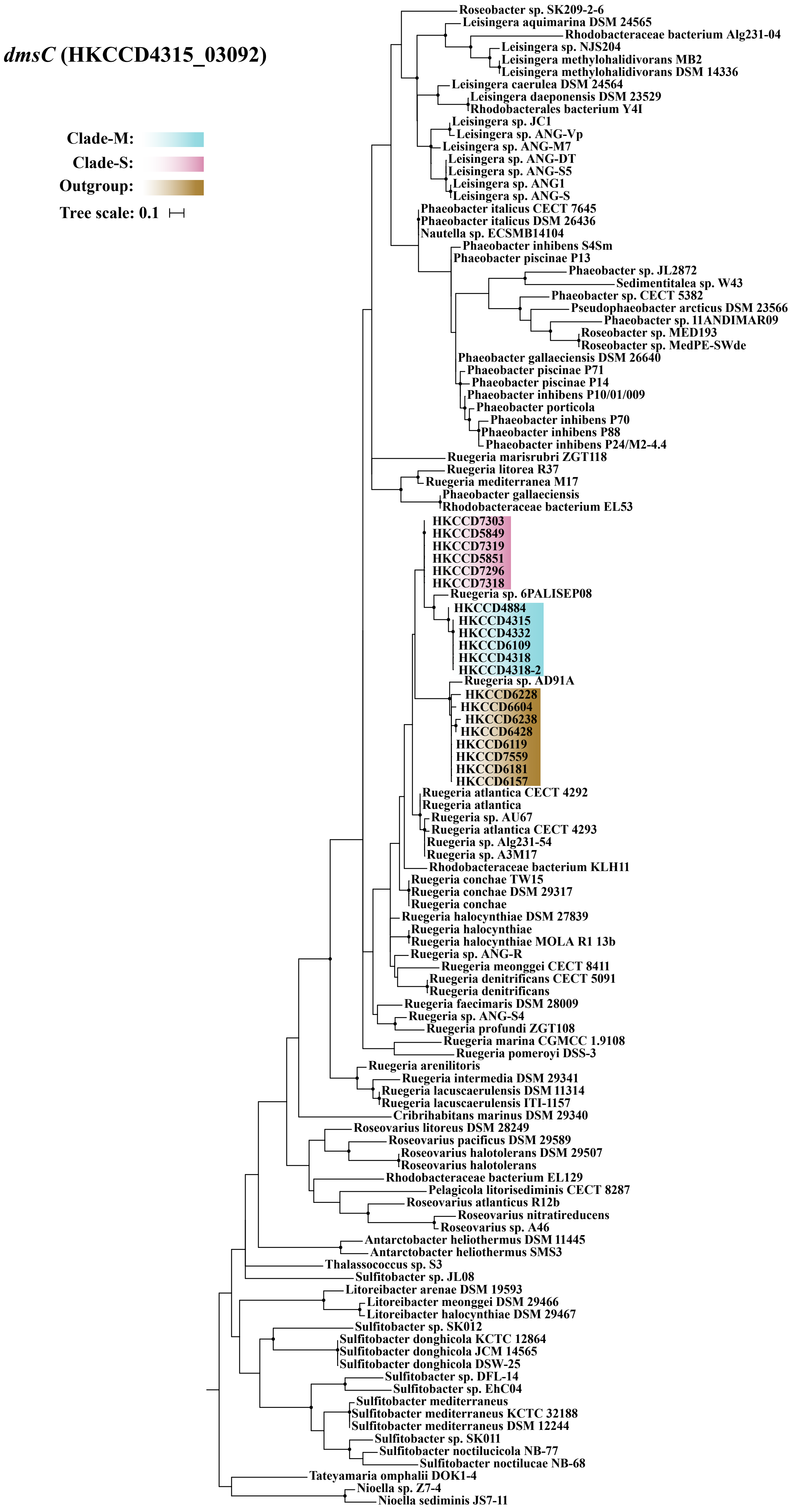
dmsC (HKCCD4315_03092)
Clade-M:
Clade-S:
Outgroup:
Tree scale: 0.1
Roseobacter sp. SK209-2-6
Leisingera aquimarina DSM 24565
Rhodobacteraceae bacterium Alg231-04
Leisingera sp. NJS204
Leisingera methylohalidivorans MB2
Leisingera methylohalidivorans DSM 14336
Leisingera caerulea DSM 24564
Leisingera daeponensis DSM 23529
Rhodobacterales bacterium Y4I
Leisingera sp. JC1
Leisingera sp. ANG-Vp
Leisingera sp. ANG-M7
Leisingera sp. ANG-DT
Leisingera sp. ANG-S5
Leisingera sp. ANG1
Leisingera sp. ANG-S
Phaeobacter italicus CECT 7645
Phaeobacter italicus DSM 26436
Nautella sp. ECSMB14104
Phaeobacter inhibens S4Sm
Phaeobacter piscinae P13
Phaeobacter sp. JL2872
Sedimentitalea sp. W43
Phaeobacter sp. CECT 5382
Pseudophaeobacter arcticus DSM 23566
Phaeobacter sp. 11ANDIMAR09
Roseobacter sp. MED193
Roseobacter sp. MedPE-SWde
Phaeobacter gallaeciensis DSM 26640
Phaeobacter piscinae P71
Phaeobacter piscinae P14
Phaeobacter inhibens P10/01/009
Phaeobacter porticola
Phaeobacter inhibens P70
Phaeobacter inhibens P88
Phaeobacter inhibens P24/M2-4.4
Ruegeria marisrubri ZGT118
Ruegeria litorea R37
Ruegeria mediterranea M17
Phaeobacter gallaeciensis
Rhodobacteraceae bacterium EL53
HKCCD7303
HKCCD5849
HKCCD7319
HKCCD5851
HKCCD7296
HKCCD7318
Ruegeria sp. 6PALISEP08
HKCCD4884
HKCCD4315
HKCCD4332
HKCCD6109
HKCCD4318
HKCCD4318-2
Ruegeria sp. AD91A
HKCCD6228
HKCCD6604
HKCCD6238
HKCCD6428
HKCCD6119
HKCCD7559
HKCCD6181
HKCCD6157
Ruegeria atlantica CECT 4292
Ruegeria atlantica
Ruegeria sp. AU67
Ruegeria atlantica CECT 4293
Ruegeria sp. Alg231-54
Ruegeria sp. A3M17
Rhodobacteraceae bacterium KLH11
Ruegeria conchae TW15
Ruegeria conchae DSM 29317
Ruegeria conchae
Ruegeria halocynthiae DSM 27839
Ruegeria halocynthiae
Ruegeria halocynthiae MOLA R1 13b
Ruegeria sp. ANG-R
Ruegeria meonggei CECT 8411
Ruegeria denitrificans CECT 5091
Ruegeria denitrificans
Ruegeria faecimaris DSM 28009
Ruegeria sp. ANG-S4
Ruegeria profundi ZGT108
Ruegeria marina CGMCC 1.9108
Ruegeria pomeroyi DSS-3
Ruegeria arenilitoris
Ruegeria intermedia DSM 29341
Ruegeria lacuscaerulensis DSM 11314
Ruegeria lacuscaerulensis ITI-1157
Cribrihabitans marinus DSM 29340
Roseovarius litoreus DSM 28249
Roseovarius pacificus DSM 29589
Roseovarius halotolerans DSM 29507
Roseovarius halotolerans
Rhodobacteraceae bacterium EL129
Pelagicola litorisediminis CECT 8287
Roseovarius atlanticus R12b
Roseovarius nitratireducens
Roseovarius sp. A46
Antarctobacter heliothermus DSM 11445
Antarctobacter heliothermus SMS3
Thalassococcus sp. S3
Sulfitobacter sp. JL08
Litoreibacter arenae DSM 19593
Litoreibacter meonggei DSM 29466
Litoreibacter halocynthiae DSM 29467
Sulfitobacter sp. SK012
Sulfitobacter donghicola KCTC 12864
Sulfitobacter donghicola JCM 14565
Sulfitobacter donghicola DSW-25
Sulfitobacter sp. DFL-14
Sulfitobacter sp. EhC04
Sulfitobacter mediterraneus
Sulfitobacter mediterraneus KCTC 32188
Sulfitobacter mediterraneus DSM 12244
Sulfitobacter sp. SK011
Sulfitobacter noctilucicola NB-77
Sulfitobacter noctilucae NB-68
Tateyamaria omphalii DOK1-4
Nioella sp. Z7-4
Nioella sediminis JS7-11

**Figure S10.** The maximum likelihood phylogeny of the core gene families *ureA*, *ureB*, *ureC*, *ureD*, *ureE* and *ureF* involved in urea utilization constructed by IQ-TREE v1.6.5. Solid circles at the nodes indicate that the ultrafast bootstrap support value of the branch is ≥ 80%. The scale bar indicates number of substitutions per site. The members of clade-M, clade-S and outgroup clade were shadowed with blue, pink and gold, respectively.

### *ureA* (HKCCD4315_03756)

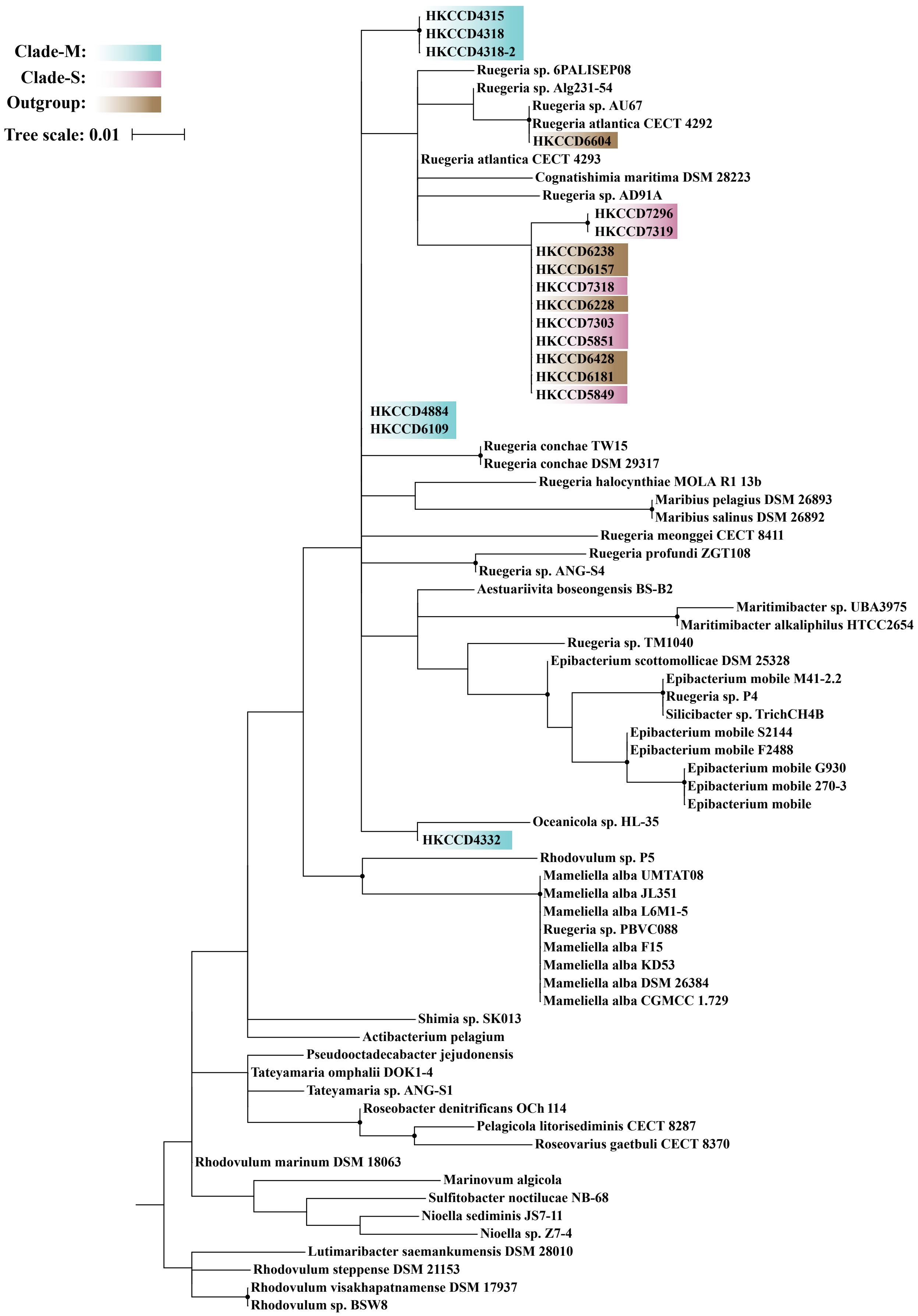

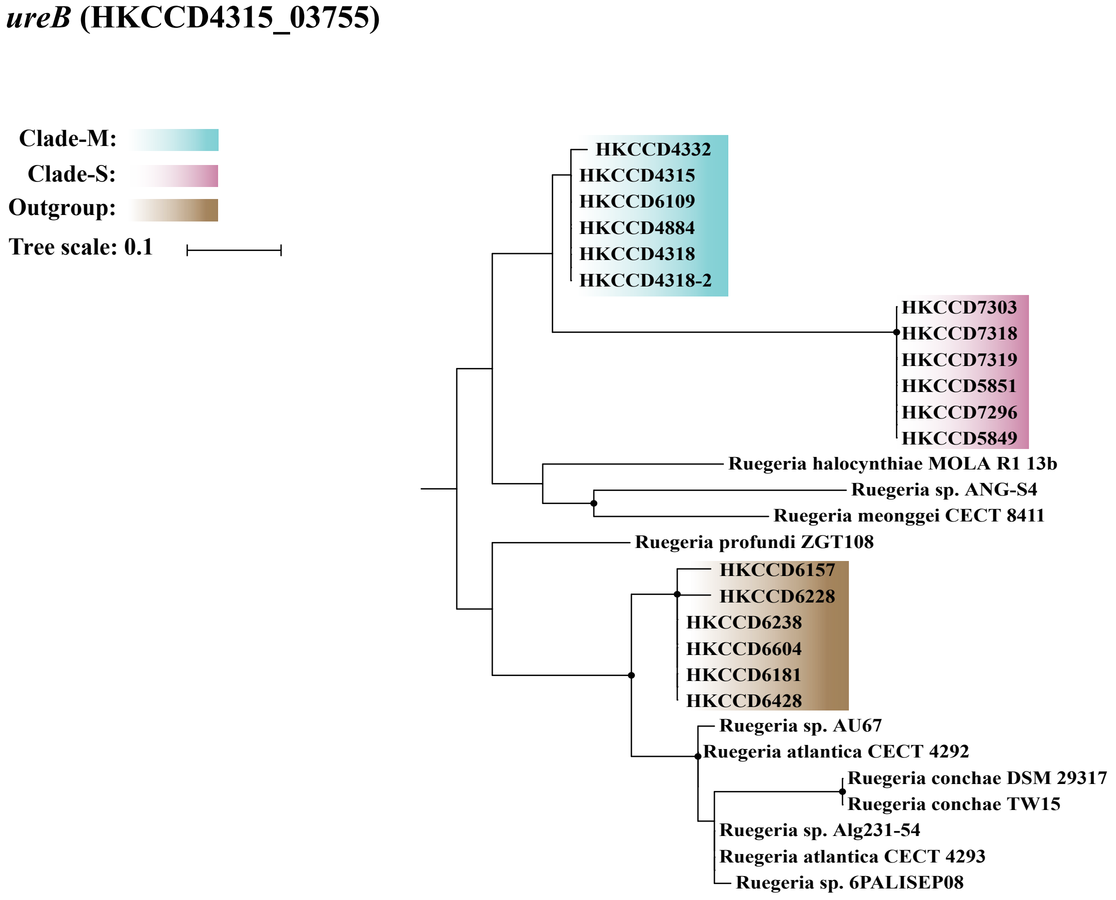

ureB (HKCCD4315_03755)
Clade-M:
Clade-S:
Outgroup:
Tree scale: 0.1
HKCCD4332
HKCCD4315
HKCCD6109
HKCCD4884
HKCCD4318
HKCCD4318-2
HKCCD7303
HKCCD7318
HKCCD7319
HKCCD5851
HKCCD7296
HKCCD5849
Ruegeria halocynthiae MOLA R1 13b
Ruegeria sp. ANG-S4
Ruegeria meonggei CECT 8411
Ruegeria profundi ZGT108
HKCCD6157
HKCCD6228
HKCCD6238
HKCCD6604
HKCCD6181
HKCCD6428
Ruegeria sp. AU67
Ruegeria atlantica CECT 4292
Ruegeria conchae DSM 29317
Ruegeria conchae TW15
Ruegeria sp. Alg231-54
Ruegeria atlantica CECT 4293
Ruegeria sp. 6PALISEP08


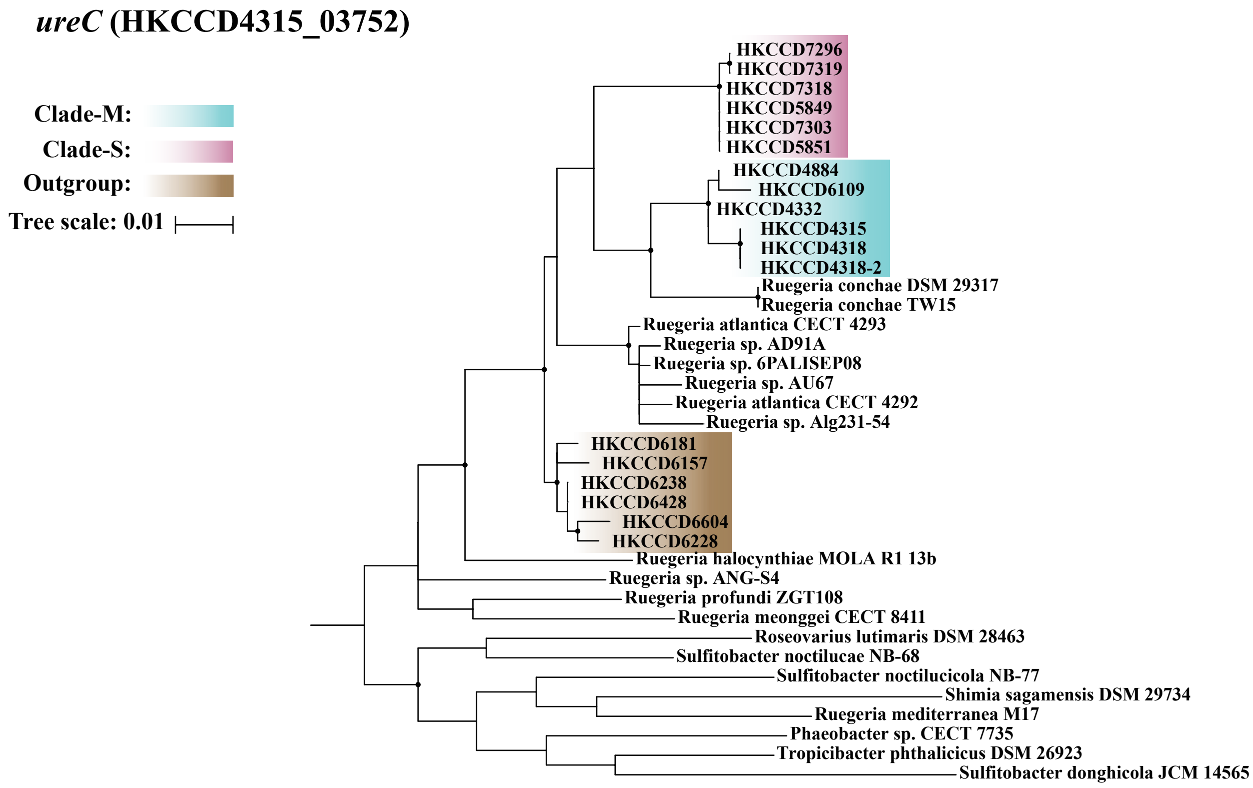

ureC (HKCCD4315_03752)
Clade-M:
Clade-S:
Outgroup:
Tree scale: 0.01
HKCCD7296
HKCCD7319
HKCCD7318
HKCCD5849
HKCCD7303
HKCCD5851
HKCCD4884
HKCCD6109
HKCCD4332
HKCCD4315
HKCCD4318
HKCCD4318-2
Ruegeria conchae DSM 29317
Ruegeria conchae TW15
Ruegeria atlantica CECT 4293
Ruegeria sp. AD91A
Ruegeria sp. 6PALISEP08
Ruegeria sp. AU67
Ruegeria atlantica CECT 4292
Ruegeria sp. Alg231-54
HKCCD6181
HKCCD6157
HKCCD6238
HKCCD6428
HKCCD6604
HKCCD6228
Ruegeria halocynthiae MOLA R1 13b
Ruegeria sp. ANG-S4
Ruegeria profundi ZGT108
Ruegeria meonggei CECT 8411
Roseovarius lutimaris DSM 28463
Sulfitobacter noctilucae NB-68
Sulfitobacter noctilucicola NB-77
Shimia sagamensis DSM 29734
Ruegeria mediterranea M17
Phaeobacter sp. CECT 7735
Tropicibacter phthalicicus DSM 26923
Sulfitobacter donghicola JCM 14565

***ureD* (HKCCD4315_03757)**

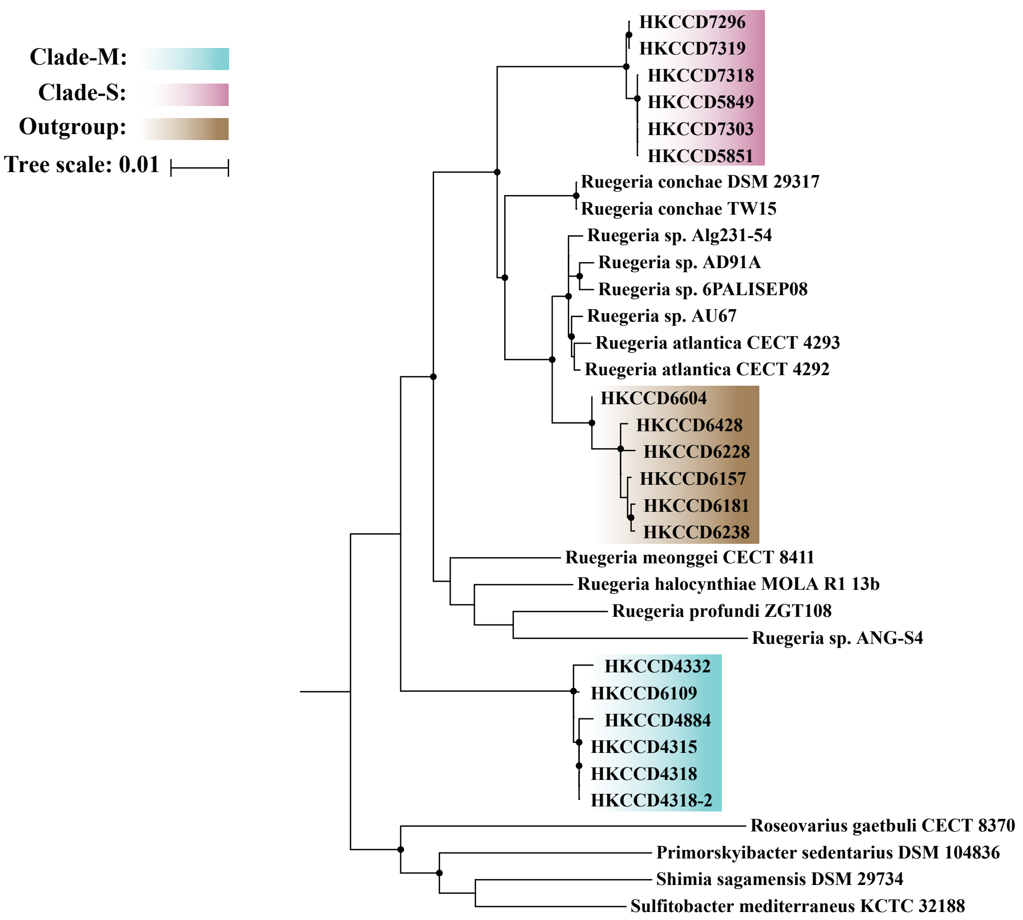


***ureE* (HKCCD4315_03750)**

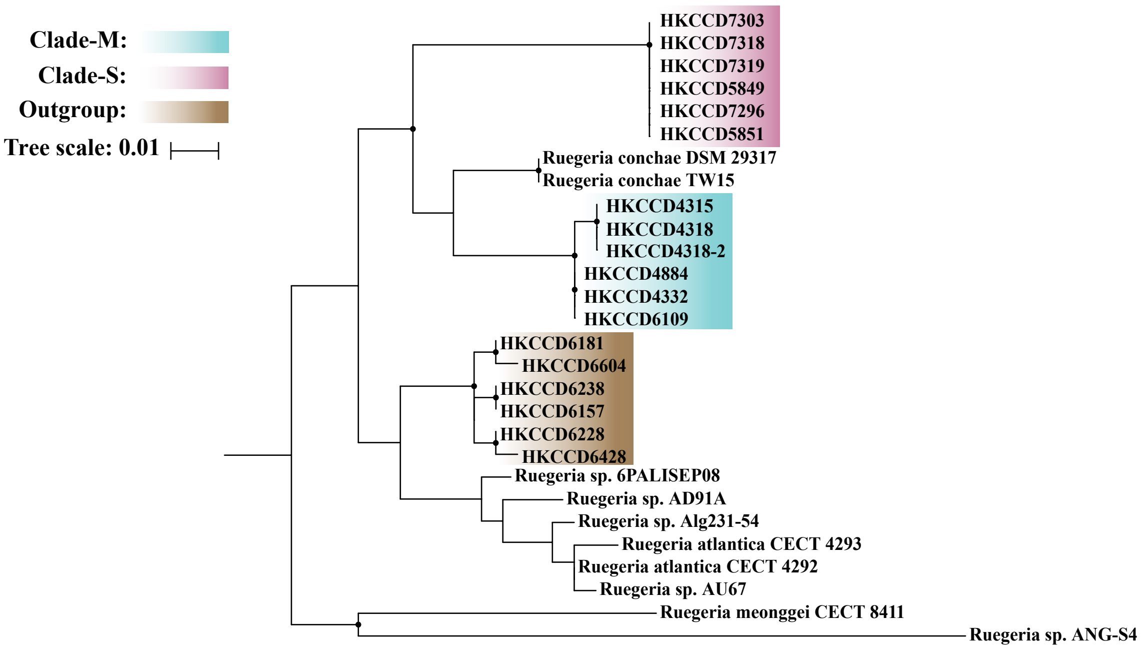

***ureF*** **(HKCCD4315_03749)**

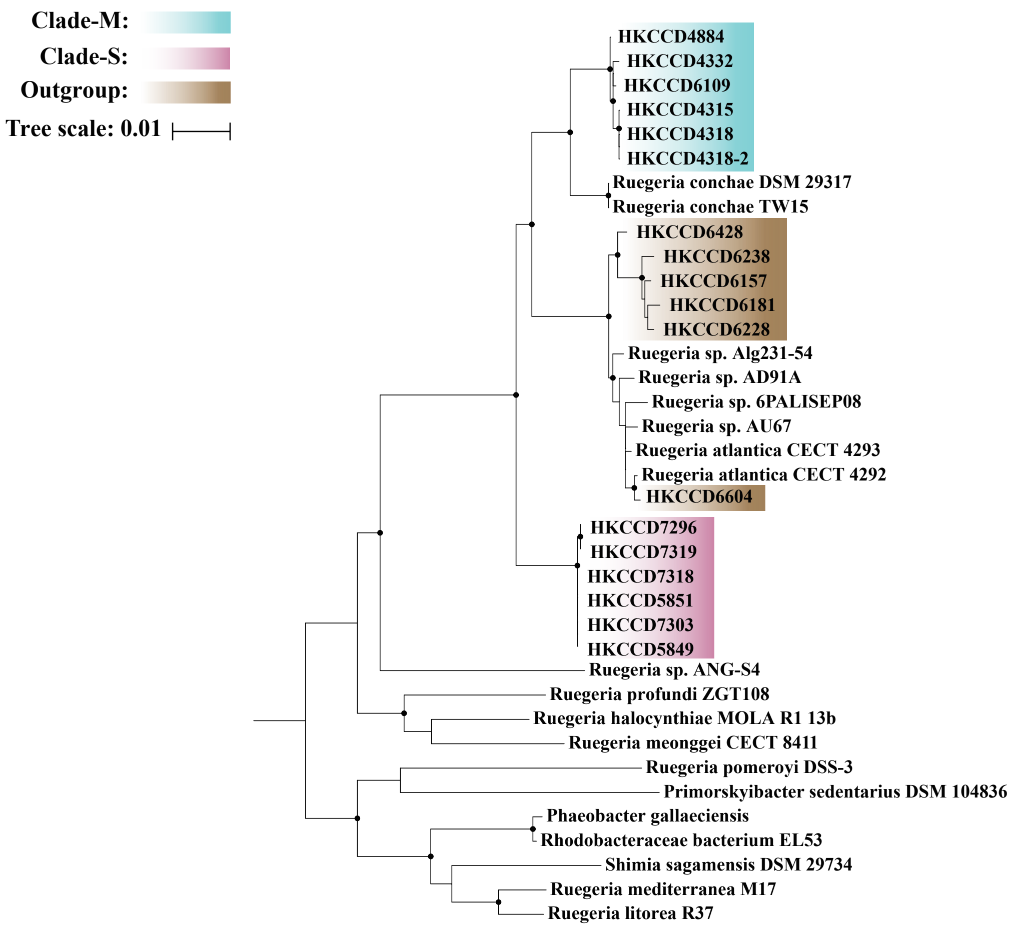

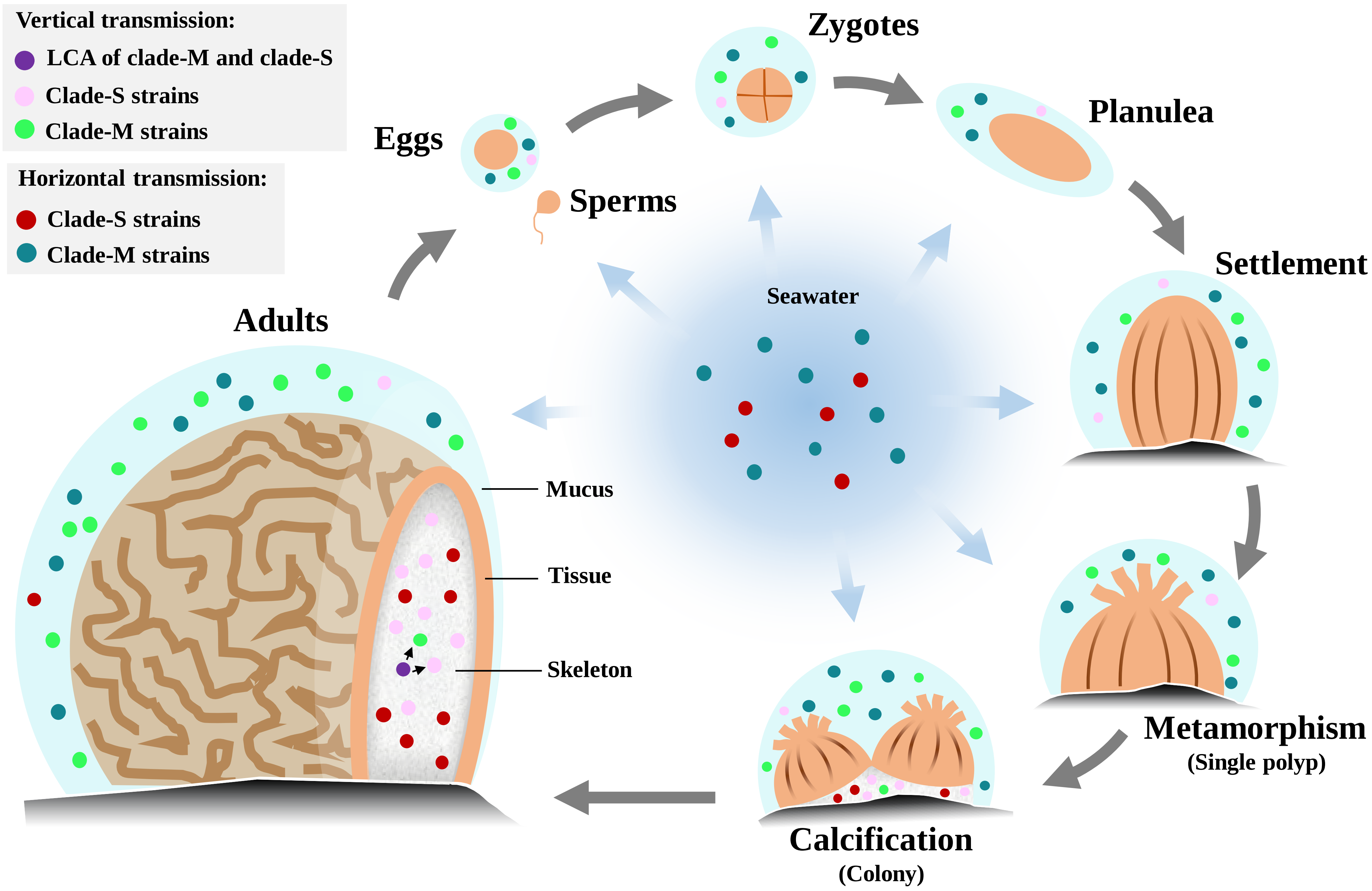


**Figure S11.** The life cycle of *Platygyra acuta* showing the origin of mucus and skeleton associated bacteria. The grey arrows indicate the vertical transmission of bacteria. The cyan arrows indicate the horizontal transmission of bacteria.

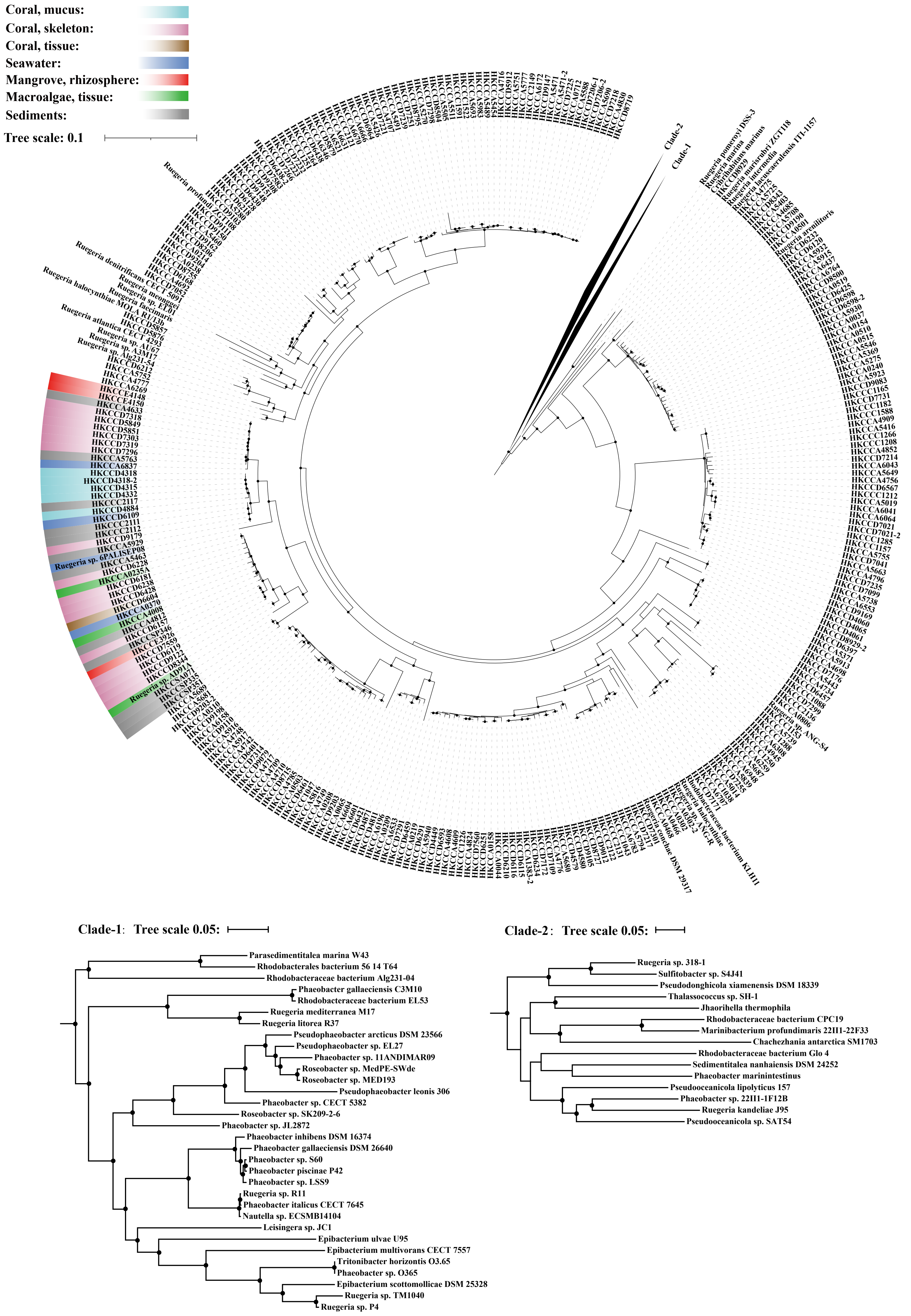


**Figure S12.** The maximum likelihood phylogeny of 454 *Ruegeria* and related strains. Solid circles at the nodes indicate that the ultrafast bootstrap support value of the branch is ≥ 95%. The scale bar indicates number of substitutions per site. The habitats of the expanded *Ruegeria* population were shadowed with different colours.

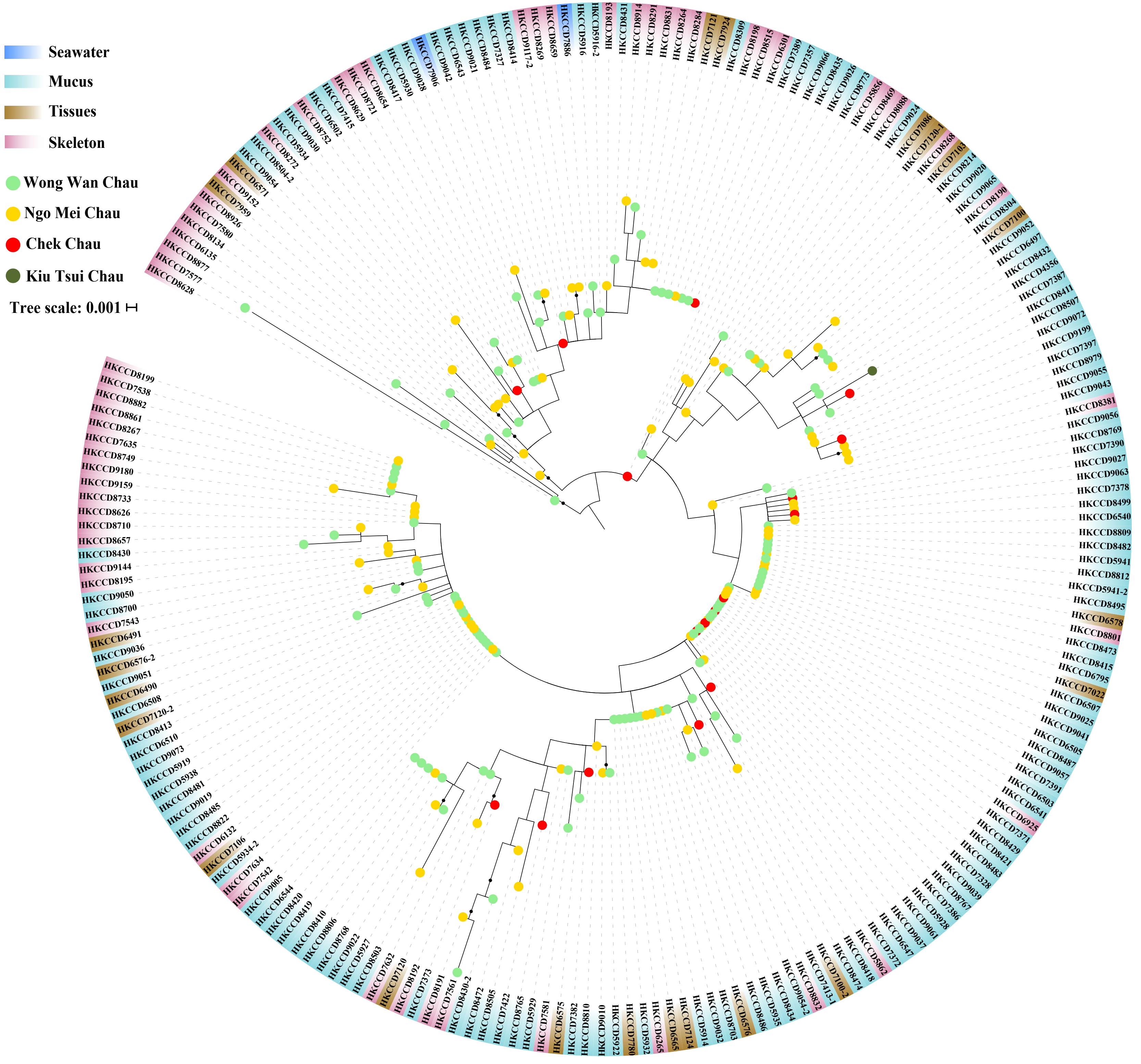


**Figure S13.** The maximum-likelihood phylogenomic tree of the genetically uniform *Rhodobacteraceae* population consisting of 214 strains isolated from four coral individuals of the same coral species *Platygyra acuta*, each collected from a distinct location in Hong Kong. Solid black dots at the nodes indicate that the ultrafast bootstrap support value of the branch is ≥ 80%. The scale bar indicates number of substitutions per site. Strain names are coloured with coral compartment information. The coloured dots at tree tips represent the sampling sites of associated coral individuals.
